# Repurposing antiviral drugs as a new avenue for *Klebsiella pneumoniae* decolonization

**DOI:** 10.64898/2026.05.14.725135

**Authors:** Nicole L. Anderson, Krista Gunter, Katlyn Todd, Valerie Velázquez-Colón, Maria Casiano, Nithyashree Maheswaran, Amanda Blankenberger, Arnav Singh, Ryan F. Relich, Natasha L. Tilston, Jay Vornhagen

## Abstract

*Klebsiella pneumoniae* (Kp) is a common antibiotic-resistant pathogen that colonizes the gastrointestinal tract and can disseminate to peripheral sites, causing a range of infections including bacteremia, urinary tract infections, and pneumonia. Intestinal colonization with Kp is a risk factor for subsequent infection, as the colonizing strain frequently corresponds to the infecting isolate. Accordingly, targeting Kp prior to dissemination at the site of colonization through decolonization strategies offers a promising approach to mitigate infection risk. In this study, we evaluated the repurposing of existing drugs not indicated for antibacterial use as candidates for Kp decolonization. To this end, we screened an antiviral compound library for their activity against Kp. We identified and validated six compounds with poorly characterized activity against Kp. Then, we screened a library of clinical Kp strains against a subset of these compounds and found that their activity was strain-specific to degrees that differed based on the compound. We also tested the activity of these compounds in conditions relevant to the human gut and determined the activity of these candidates was dependent on biological context. Focusing in on the most successful target drug from the human *ex vivo* experiment, 3′-azido-3′-deoxythymidine (zidovudine or AZT), a transposon mutant screen identified multiple genetic determinants of AZT susceptibility, with genes involved in DNA maintenance and central metabolic processes contributing to AZT sensitivity. Serial passage of Kp in AZT resulted in the repeated emergence of mutations in *tdk*, a gene that encodes thymidine kinase, suggesting that alterations in thymidine metabolism contribute to the development of AZT resistance. Finally, we show that development of AZT resistance leads to collateral sensitivity to trimethoprim, suggesting promise for combination treatment. Collectively, these findings support further investigation of antiviral drugs as potential gut decolonization therapies for Kp.

## Introduction

*Klebsiella pneumoniae* (Kp) is a leading contributor to antimicrobial resistance-associated mortality, as infections caused by multidrug resistant Kp strains (MDR-Kp) result in >50% mortality and $40K in healthcare costs per case.^1^ Projections estimate antibiotic-resistant bacteria may cause 10 million deaths annually by 2050 and surpass cancer and diabetes as global leading causes of death, leading to an economic loss of up to $100 trillion.^2^ Additionally, hypervirulent strains of Kp (hvKp) have independently emerged and are disseminating globally.^3^ Disturbingly, there is reported genetic convergence of MDR- and hvKp; thus, there is an urgent need to develop novel strategies to combat these dangerous emergent strains.^4,5^

Kp is a common opportunistic pathogen that causes respiratory tract, urinary tract, and bloodstream infections. The primary reservoir for infectious Kp is the gut, and most (∼80%) of infections are caused by the original colonizing strain.^6^ Colonization is generally a prerequisite for infection, making it an attractive intervention point. Further, the risk of infection increases upwards of 400% when Kp colonizes the gut. Additionally, infection risk also increases if Kp becomes a dominant member of the gut microbial community.^6^ As such, there is increasing interest in developing therapies to reduce infection risk in colonized patients through targeted decolonization. Decolonization is a medical strategy that involves “removing or reducing the burden of a pathogen, either temporarily or permanently” from a non-sterile site such as the skin, reproductive tract or gut.^7^ Gentamicin is an FDA approved antibiotic for treatment of Kp infection that can be used for gut decontamination or decolonization of Kp to prevent infection. Although this approach has been successful in clinical reports, it requires a prolonged high-dose treatment, is effective only against gentamicin-susceptible strains, and such extended antibiotic use significantly reduces gut microbiota diversity which makes this decolonization method suboptimal for preserving gut microbiome health.^8,9^ An alternative treatment is decolonization through bacteriophages, which can selectively target Kp while preserving gut microbiome diversity.

Thousands of Kp-specific phages have been identified, with growing evidence of efficacy against multidrug-resistant strains.^10^ However, therapeutic application is limited by the highly variable polysaccharide capsule (K antigen), a primary phage receptor. Diversity in capsule types and the ability of Kp to alter or lose capsule expression thus restricts phage-host range and can lead to treatment failure.^11–13^ Another option that shows promise is the reduction of Kp gut density through restoration of gut diversity via fecal microbiota transplantation (FMT), which has been successful in clinical case reports. FMT is currently used as an experimental therapy, as the risks include variation in success over a longer period of time and regulation/safety concerns with transferring undesirable pathogens to the host.^14–16^ Given the challenges of these therapies and the impending outpacing of antibiotics to combat rising converging strains, it is clear that there is a need for the development of additional alternative Kp decolonization methods or to adopt a combined approach integrating multiple methods.

These limitations highlight the need for alternative strategies that are both targeted and clinically feasible. Advances in pharmacology, genomics, and computational screening have transformed drug repurposing to identify new uses for approved drugs.^17,18^ In this vein, antiviral drugs are intended to target viral replication; however, antivirals can have off-target effects, including the inhibition of bacterial growth.^19,20^ The viability of using antiviral drugs against Kp and whether these drugs could be repurposed for gut decolonization is underexplored. This work aims to identify novel drug therapies for decolonization of Kp by targeting the bacteria with novel approaches such as antiviral drugs, which will ideally lead to a reduction in gut density and decreased risk of infection dissemination.

## Results

### Antiviral compounds are bactericidal against Kp

We screened a library of 509 antiviral compounds for their ability to restrict Kp growth. We used a KPPR1 strain encoding the pJL1-sfGFP plasmid for this screen. Growth assays were performed under defined growth conditions at 37 °C with shaking (220 rpm), using a standardized initial inoculum (OD_600_ = 0.01) and compounds applied at a final concentration of 1 mM. Measurements were taken at the 24-hour timepoint, and all experiments were conducted in triplicate biological replicates. Bacterial growth was measured by both OD_600_ and relative fluorescent units (RFUs). RFUs were measured to account for compounds that interfered with OD_600_ readings, such as crystal violet. *z*-score normalization was applied to RFU and OD_600_ measurements to account for experimental variability across the 509-compound screen, with hits defined as values deviating more than 2.5 standard deviations below the mean for OD_600_ and an RFU-based *z-*score that was directionally concordant (*z-*score ≤ −1). The output of RFUs and OD_600_ for the 509 compounds tested revealed 19 hits that restricted KPPR1 growth (**Figure 1A-B**, **Table 1, Supplemental Dataset 1**). 12/19 hits were antibiotic and anti-septic compounds with known anti-Kp activity (**Table 1**); therefore, we selected seven compounds (bithionol, hinokitiol, fingolimod, NH125, prulifloxacin, AZT, and montelukast), with previously uncharacterized activity against Kp, as well as two controls (gentamicin and ivermectin) for validation.

**Table 1.** Anti-Kp drug hits identified in screening.

| COMPOUND | CAS NUMBER | KNOWN ACTIVITY AGAINST *KLEBSIELLA* | ACTIVITY | Z-SCORE (OD600) | Z-SCORE (RFU) |
| --- | --- | --- | --- | --- | --- |
| Cetylpyridinium chloride | 123-03-5 | Yes | Antiseptic | -5.47 | -5.33 |
| Ciprofloxacin HCL | 93107-08-5 | Yes | Fluoroquinolone antibiotic | -5.47 | -4.96 |
| Ciprofloxacin HCL hydrate | 86393-32-0 | Yes | Fluoroquinolone antibiotic | -5.47 | -4.83 |
| Gentamycin sulfate | 1405-41-0 | Yes | Aminoglycoside antibiotic | -5.41 | -5.18 |
| Chlorhexidine HCL | 3697-42-5 | Yes | Antiseptic | -5.38 | -5.00 |
| Tetracycline HCL | 64-75-5 | Yes | Tetracycline antibiotic | -5.36 | -5.11 |
| Oxytetracycline HCL | 2058-46-0 | Yes | Tetracycline antibiotic | -5.31 | -5.11 |
| Chlorhexidine diacetate | 56-95-1 | Yes | Antiseptic | -5.29 | -5.28 |
| Bithionol | 97-18-7 | No | Inhibitor of soluble adenylyl cyclase | -4.95 | -4.38 |
| Oxytetracycline (Terramycin) | 79-57-2 | Yes | Tetracycline antibiotic | -4.60 | -2.40 |
| Azithromycin dihydrate | 117772-70-0 | Yes | Macrolide antibiotic | -4.55 | -4.13 |
| **Azithromycin** | 83905-01-5 | Yes | Macrolide antibiotic | -4.31 | -3.67 |
| **AZT** | 30516-87-1 | No | Reverse transcriptase inhibitor | -4.18 | -1.62 |
| **NH125** | 278603-08-0 | No | Selective eEF-2 kinase inhibitor | -4.02 | -4.27 |
| **Prulifloxacin (Pruvel)** | 123447-62-1 | No | Fluoroquinolone antibiotic | -3.69 | -3.64 |
| **Fingolimod (FTY720)** | 162359-56-0 | No | S1P receptors agonist | -3.49 | -3.47 |
| **Hinokitiol** | 499-44-5 | No | Apoptosis inducer and iron chelator | -2.92 | -1.22 |
| **Methacycline HCL** | 3963-95-9 | Yes | Tetracycline antibiotic | -2.61 | -4.84 |
| **Montelukast sodium** | 151767-02-1 | No | Leukotriene receptor antagonist | -2.55 | -2.32 |
| **Ivermectin** | 70288-86-7 | No | P2X purinergic receptor | 0.13 | 0.18 |

**Figure 1.**
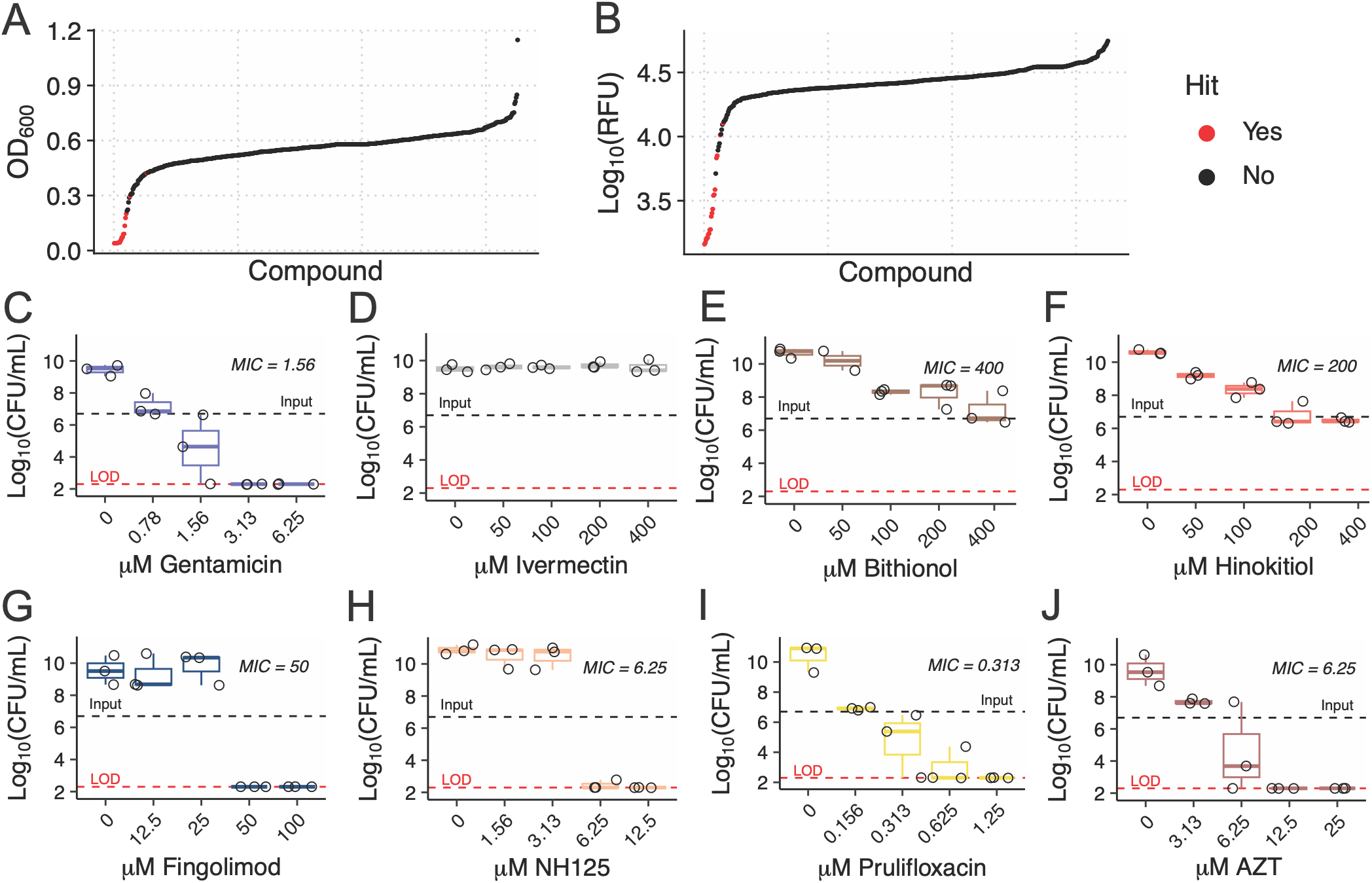
Antiviral drugs are bactericidal against Kp strain KPPR1. KPPR1pJL1-sfGFP was grown overnight with the arrayed antiviral library of 509 compounds and resulting growth determined via the OD_600_ (**A**) and relative fluorescent units (RFUs, **B**). The minimum inhibitory concentrations (MIC) and minimum bactericidal concentrations (MBC) were determined for positive control gentamicin (**C**), negative control ivermectin (**D**) and candidate compounds: bithionol (**E**), hinokitiol (**F**), fingolimod (**G**), NH125 (**H**), prulifloxacin (**I**) and AZT (**J**). For **A-B**, the mean RFU and OD_600_ values from three biological replicates are shown. For **C-J**, the black dashed line represents the input CFU/mL concentration, the red dashed line represents the limit of detection (LOD) for the assay, each datapoint represents a single biological replicate, and the boxplots shown the mean and interquartile range, with tails indicating the range.

To validate our screen, we determined the minimum inhibitory concentration (MIC) and minimum bactericidal concentration (MBC) of each of the selected compounds. These two phenotypes can be compared to determine bacteriostatic or bactericidal activity of a given compound. Bacteriostatic compounds have an MBC greater than a 2-fold higher concentration than the MIC concentration and bactericidal compounds have an MBC concentration no greater than 2-fold higher than the MIC concentration. Gentamicin, an antibiotic with known activity against Kp, was used as a positive control. As expected, this compound exhibited bactericidal activity, wherein the MIC was 1.56 μM and the MBC was 1.56-3.13 μM (**Figure 1C**). Ivermectin, an anti-parasitic with no expected anti-Kp activity, was chosen as a negative control. As expected, this compound showed no inhibitory activity (**Figure 1D**). The candidate compounds bithionol and hinokitiol were determined to be bacteriostatic (**Figure 1E-F**), which aligns with the literature as these are known iron chelators/metal-interacting compounds that sequester iron away from bacteria to prevent growth.^21,22^ Fingolimod, NH125, prulifloxacin, and AZT all exhibited bactericidal activity (**Figure 1G-J**). Prulifloxacin exhibited the lowest MIC (0.313 μM), which is expected given that it is a known fluoroquinolone; however, to our knowledge, it has not been tested against Kp which is why we included it in our validation studies. The only compound that failed validation (no activity against Kp) was montelukast (**Supplemental Figure 1**). Collectively, we validated 8/9 candidate compounds, indicating that our original screening assay was high quality. Although the mechanisms of action for each of these bactericidal drugs vary, our results indicate that there is potential for antiviral compounds to have success in limiting Kp.

### Kp clinical isolates from various infection sites are susceptible to AZT, fingolimod and NH125

To assess the generalizability of our findings across Kp strains, we selected the bactericidal compounds AZT, fingolimod and NH125. We cultured an array of 87 clinical Kp isolates obtained from various infection sites in the presence of gentamicin, AZT, fingolimod, or NH125 at 2X the concentration of the KPPR1 MIC. As anticipated, all clinical Kp isolates grew in LB broth and 83/87 (95.4%) were susceptible to gentamicin (**Figure 2**). Interestingly, 80/87 of Kp clinical isolates were susceptible to fingolimod and NH125 (91.9%), whereas 39/87 (44.8%) were susceptible to AZT. Select clinical isolates that varied in sample origin, JV94, JV101, and JV132 (**Supplemental Table 1**), were selected for MIC determination to validate these findings. We found that the calculated MICs for these compounds were concordant with our clinical strain screen (**Supplemental Table 2**). Additionally, we stratified these isolates by site of infection and the presence of *rmpA*, which is a marker of hypervirulence.^23^ Resistant strains were distributed across sites of infection (**Supplemental Figure 2**). Only two isolates in our dataset were PCR-confirmed *rmpA* positive and were hypermucoviscous by string test. These strains were susceptible to all compounds we tested. Collectively, these data indicate that Kp are broadly susceptible to fingolimod and NH125, but variably susceptible to AZT with no obvious biases to infection site or hypervirulence.

**Figure 2.**
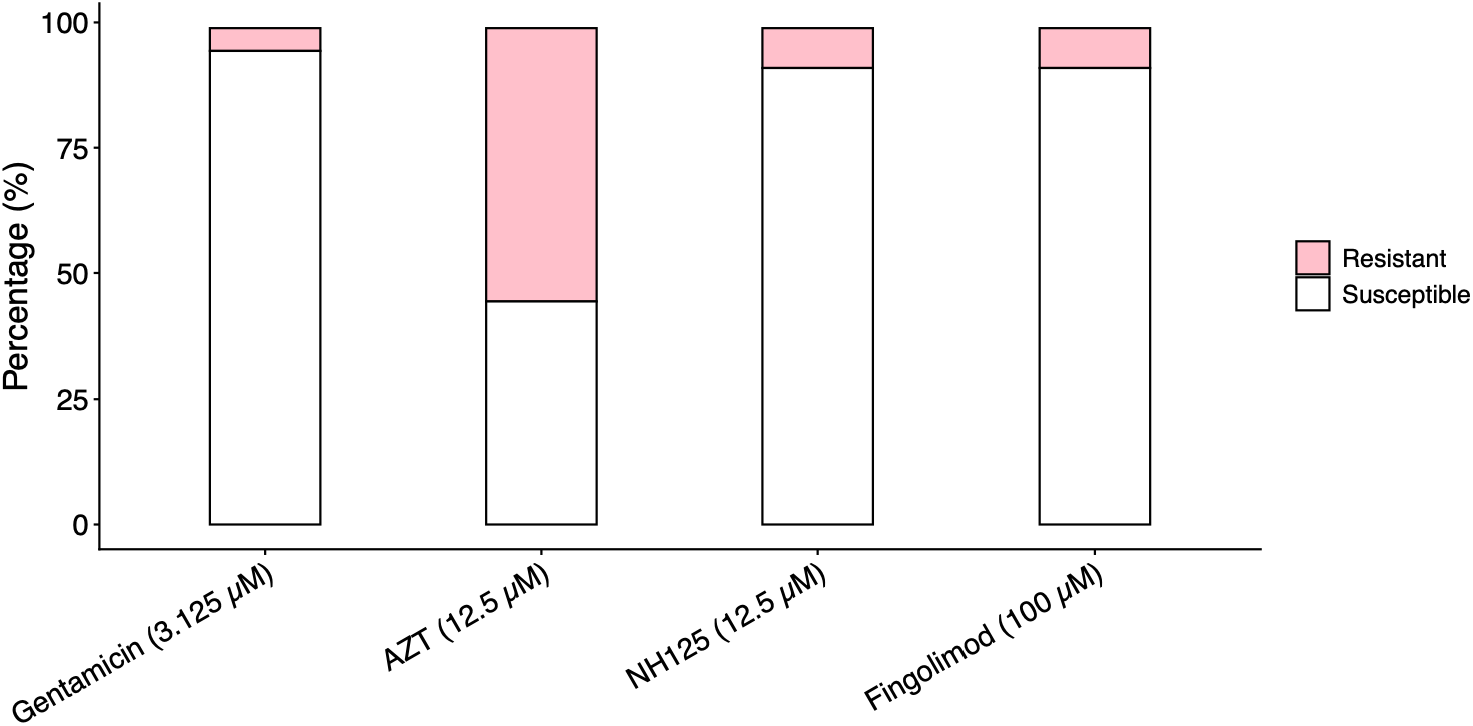
Various Kp clinical isolates are susceptible to novel antiviral drugs. Kp isolates obtained from infected patients were grown overnight in the presence of bactericidal compounds at 2X the KPPR1 MIC and determined visually to be susceptible or resistant.

### KPPR1 was susceptible to AZT, but not fingolimod or NH125, when cultured with human fecal material

While AZT, fingolimod, and NH125 showed promising activity in initial *in vitro* assays, we next evaluated their efficacy in a more physiologically relevant context for gut colonization. To assess how these anti-Kp compounds perform within a human microbiome– associated environment, KPPR1 was cultured in a human donor fecal slurry from three independent donors under both aerobic and anaerobic conditions. Each compound was tested at 16X its respective MIC to replicate clinical settings where drug administration far exceeds MIC. KPPR1 was directly inoculated into fecal samples with or without each compound and CFUs were quantified after 48 hours of incubation in anaerobic or aerobic conditions. Data were summarized as fold change (output CFUs/input CFUs). All mock-treated samples sustained robust KPPR1 growth (**Supplemental Figure 3**). All compounds effectively inhibited KPPR1 growth in LB broth under both aerobic and anaerobic conditions, confirming their intrinsic antibacterial activity. Gentamicin consistently suppressed growth across all three human donors in both oxygen conditions. AZT reduced KPPR1 growth relative to the mock-treated (1X PBS) control in a donor-dependent and oxygen-dependent fashion, with more pronounced activity under aerobic conditions, suggesting potential oxygen-modulated activity. Nonetheless, KPPR1 growth was strongly restricted in the presence of AZT in feces under anaerobic conditions (68.89-99.97% restriction compared to “Mock”). By comparison, fingolimod and NH125 retained activity in LB but showed a marked loss of efficacy in the human donor slurry in both aerobic and anaerobic conditions, indicating that components of the microbiome-derived matrix likely inactivate or sequester these compounds (**Figure 3**). Overall, these findings suggest that AZT maintains measurable activity in microbiome-relevant conditions and may represent a clinically relevant anti-Kp candidate, however further work is needed to understand the impact of complex microbial communities and oxygen tension on antibacterial activity.

**Figure 3.**
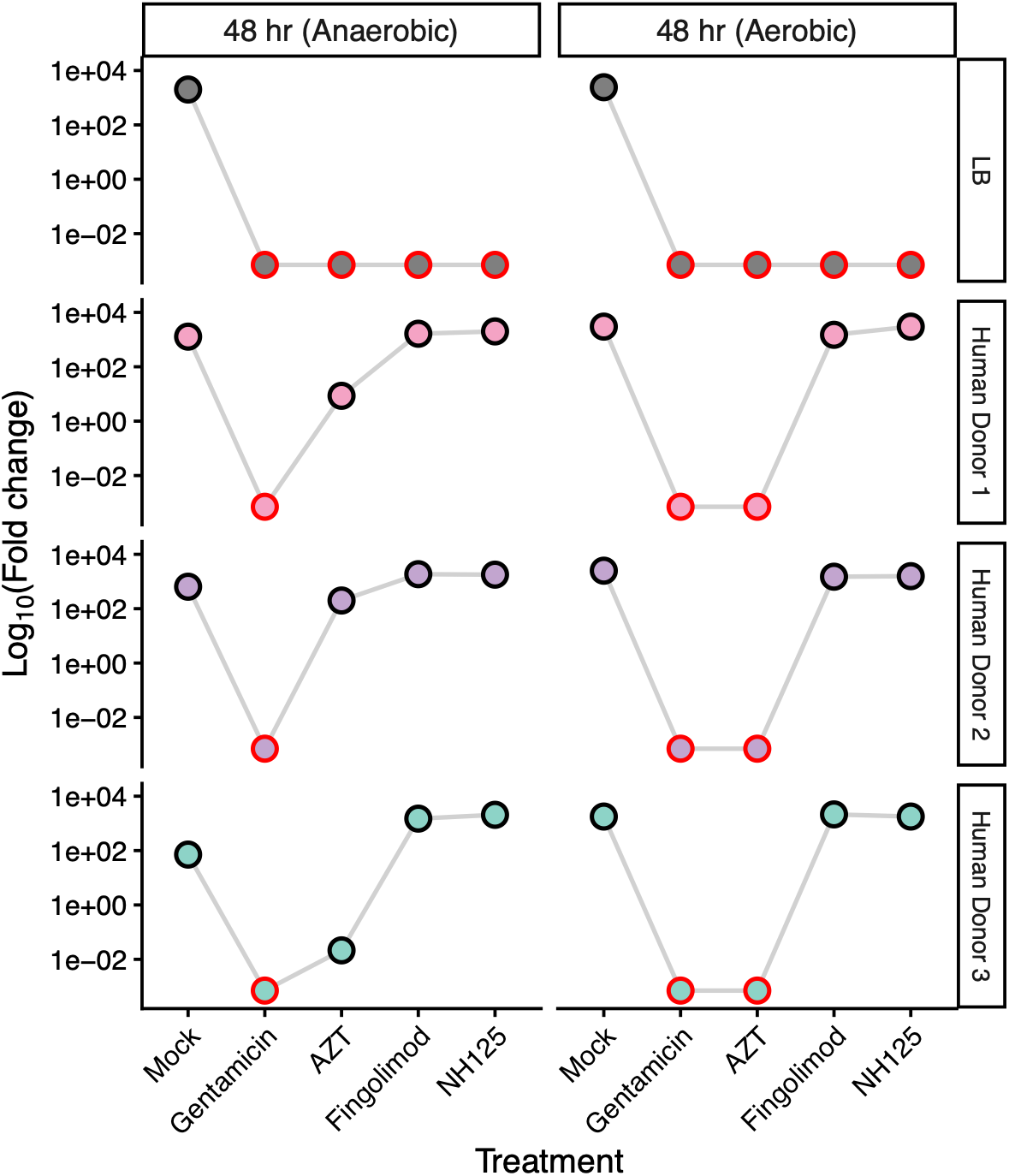
Treatment of KPPR1 with select compounds in *ex vivo* human feces results in decreased Kp growth. KPPR1 was grown overnight at 37 °C with shaking (220 rpm) and used the following day for *ex vivo* culture with fecal material from three human donors or in LB medium, in the presence of compounds at 16X their respective MIC values. Samples were inoculated with ∼10^5^ CFU. Fold change = (output CFUs)/(input CFUs). Cultures were incubated under aerobic or anaerobic conditions and sampled at 48 hours. A red circle indicates the data point is at the limit of detection for this assay.

### A transposon screen of KPPR1 mutants identified multiple genetic determinants for AZT susceptibility or resistance

Since AZT emerged as the most effective compound at limiting KPPR1 growth in our *ex vivo* human fecal cultures, we next investigated the bacterial genes underlying susceptibility and resistance. We profiled 3,719 transposon mutants with single insertions and assessed their fitness in the presence of one-quarter of the predetermined MIC (1.56 µM) of AZT. Mutant fitness was quantified using a *z*-score, and a −1.4 cutoff in all trials was applied to identify potential hits. Using this approach, we identified 132 susceptible mutants (**Figure 4A**). A complete list of *z*-scores is provided in **Supplemental Table 3**. KEGG BRITE functional hierarchy analysis was used to determine whether mutant *z*-score patterns were associated with specific functional categories. KEGG BRITE is a hierarchical classification system that groups genes with related biological functions into categories with increasing specificity. Each mapped mutant was assigned to KEGG BRITE functional hierarchies, and mean trial-level *z*-scores were summarized within the most specific functional category (Level C). Level C categories with an absolute mean *z*-score greater than 0.3 are shown in (**Figure 4B**). To reduce the influence of categories represented by few genes, only categories containing at least two genes were included in the final summary. Functional categories with negative mean *z*-scores were interpreted as groups in which gene disruption was associated with reduced growth during AZT exposure.

**Figure 4.**
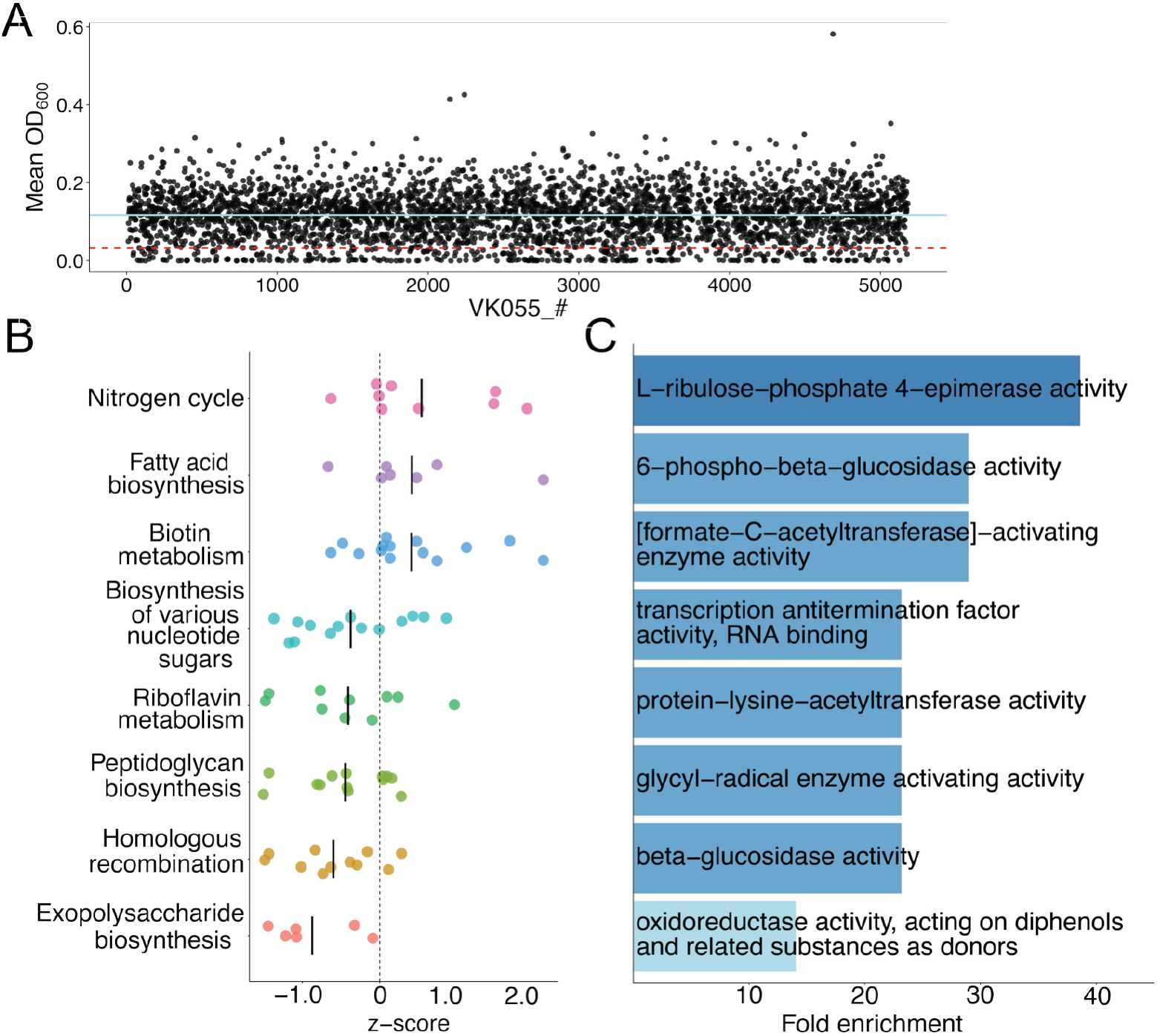
Functional analysis of AZT susceptibility in the *K. pneumoniae* transposon mutant screen. (**A**) Mean OD_600_ values for mapped transposon mutants by VK055 gene locus following AZT exposure. Each point represents an individual mutant. The solid blue line indicates the mean OD_600_ across all mutants, and the red dashed line indicates the cutoff for potential AZT-susceptibility hits. (**B)** KEGG BRITE Level C pathway analysis of mutant *z*-scores. Each point represents an individual gene-level mean trial *z*-score within the indicated Level C pathway, and the black horizontal line indicates the mean *z*-score for that pathway. Pathways shown contain more than three genes and had an absolute mean *z*-score ≥ 0.3. Negative *z*-scores indicate reduced growth during AZT exposure, while positive *z*-scores indicate increased growth relative to the trial mean. (**C**) PANTHER GO molecular function overrepresentation analysis of AZT-susceptibility hits (N = 132) mapped to putative *E. coli* orthologs. Bars show fold enrichment for significantly overrepresented GO molecular function terms with fold enrichment >10 using the *E. coli* whole-genome reference background.

Among the most negatively scoring categories were exopolysaccharide (EPS) synthesis, DNA repair, and peptidoglycan biosynthesis. The identification of DNA repair among the most negatively scoring categories is consistent with AZT’s activity as a thymidine analog and suggests that impaired DNA maintenance increases susceptibility to AZT exposure. EPS synthesis requires nucleotide-sugar precursors, suggesting that defects in EPS biosynthesis may alter nucleotide-associated metabolism and increase susceptibility to AZT.^24–26^ Similarly, peptidoglycan biosynthesis depends on nucleotide-activated precursors, which may explain why disruption of cell wall biosynthetic pathways was associated with enhanced AZT susceptibility.^27^ In contrast, mutants with disruptions in nitrogen cycle pathways, biotin metabolism, and fatty acid biosynthesis showed higher mean *z*-scores, suggesting reduced susceptibility to AZT exposure. This may be due to altered nitrogen availability reducing nucleotide synthesis, creating a slower replication rate that is less susceptible to AZT treatment; however, this interpretation will require future study.

Next, we used PANTHER Gene Ontology (GO) overrepresentation analysis to identify functional activities enriched among AZT-susceptibility hits. GO analysis organizes genes and proteins into functional categories based on biological processes, molecular function, and cellular components. AZT-susceptibility hits were mapped to *E. coli* orthologs using shared KEGG Orthology identifiers. The corresponding *E. coli* UniProt identifiers were analyzed using the PANTHER Overreprentation test using the *E. coli* whole genome as the reference list. Significant enrichment was observed in the GO molecular function category using Fisher’s exact test with false discovery rate (FDR) correction for multiple testing (**Figure 4C**). GO terms with an FDR of <0.05 and a positive fold enrichment were considered significantly overrepresented. Enriched molecular functions clustered into three broad functional groups: DNA-associated functions, metabolic enzyme activities, and carbohydrate/glycan metabolism. Representative enriched functions included single-stranded DNA binding, oxidoreductase activity, glycyl-radical enzyme activating activity, transcription antitermination factor activity, protein-lysine-acetyltransferase activity, and carbohydrate/glycoside-metabolizing enzymes such as beta-glucosidase, 6-phospho-beta-glucosidase, and L-ribulose-phosphate 4-epimerase. Overall, these findings suggest that genes involved in DNA maintenance and central metabolic processes contribute to AZT susceptibility, consistent with the KEGG BRITE analysis.

### Evolution of KPPR1 in response to AZT

We next wanted to determine whether KPPR1 could develop antimicrobial resistance to AZT. To test this, three KPPR1 lineages were serially passaged in the presence of 1.56 µM AZT (¼ the KPPR1 MIC) for 5 passages and 3.125 µM AZT (½ the KPPR1 MIC) for another 5 passages, and the resulting mutations were annotated in evolved lineages compared to the starting culture. As shown in **Table 2**, all three evolved lineages contained mutations in the thymidine kinase family protein encoded by *tdk*. Lineage 1 contained a premature stop codon (L61*), lineage 2 contained a missense mutation (E28A), and lineage 3 contained a 1-bp deletion within the *tdk* coding region, which is predicted to disrupt the resulting protein. Notably, lineage 3 also acquired mutations in *ftsX* and *wcaJ*, although these mutations were not observed in the other lineages. Since thymidine kinases are involved in the metabolism and phosphorylation of thymidine and thymidine analogs, mutations that disrupt Tdk function could reduce the ability of AZT to exert its bactericidal effect. The repeated identification of *tdk* mutations across independently evolved lineages therefore suggests that alterations in thymidine kinase may contribute to AZT resistance in KPPR1. Further, these findings support the thymidine salvage pathway as a potential mechanism of AZT resistance, whereby reduced thymidine kinase activity may limit AZT sensitivity.^28^

**Table 2.**
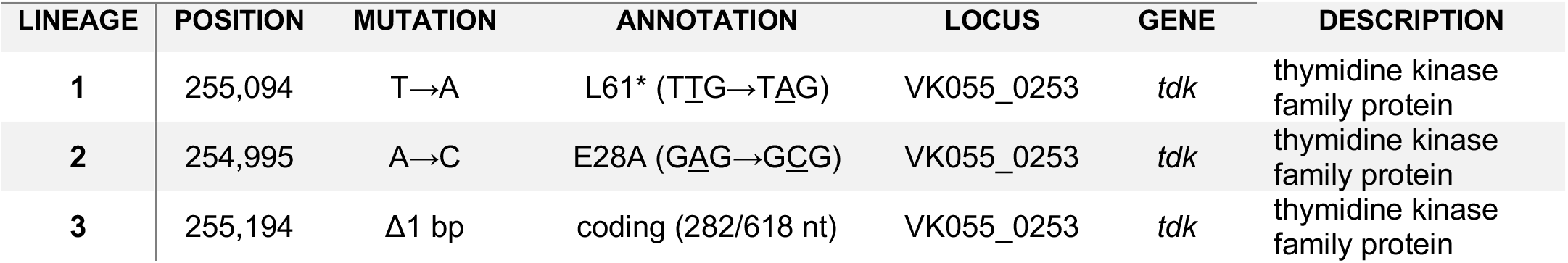
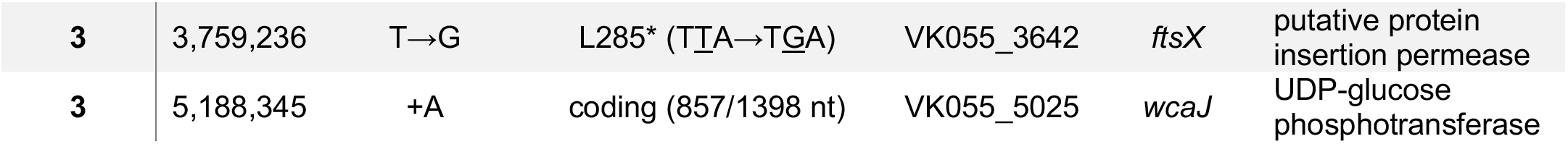
Mutations in AZT-evolved lineages.

### Development of AZT resistance induces collateral sensitivity to TMP

Given the AZT resistance observed in AZT-evolved KPPR1 lineages (**Table 2**), we next wanted to determine whether this resistance altered susceptibility to trimethoprim (TMP). Kp can obtain thymidine nucleotides through two major pathways: salvage of extracellular thymidine and *de novo* synthesis. The salvage pathway involves the uptake of extracellular thymidine followed by its phosphorylation by thymidine kinase to generate thymidine monophosphate (dTMP), whereas the *de novo* pathway generates dTMP from deoxyuridine monophosphate (dUMP) through a folate-dependent reaction. AZT activity is dependent on the thymidine salvage pathway because AZT must be phosphorylated by thymidine kinase to become active, whereas TMP disrupts *de novo* dTMP synthesis. Therefore, alterations in thymidine utilization that confer AZT resistance may increase dependence on *de novo* dTMP synthesis and consequently increase susceptibility to TMP. TMP is an antimicrobial that inhibits dihydrofolate reductase (DHFR), disrupting folate metabolism and ultimately inhibiting *de novo* dTMP synthesis.^28^ To test this, we performed a TMP killing assay on the KPPR1 ancestral and evolved lineages. As shown in **Figure 5**, the AZT-evolved lineages demonstrated enhanced sensitivity to TMP compared to the ancestral KPPR1 strain. Taken together, these results suggest that the development of AZT resistance induces collateral sensitivity, or the evolution of resistance to one antimicrobial results in increased susceptibility to TMP.

**Figure 5.**
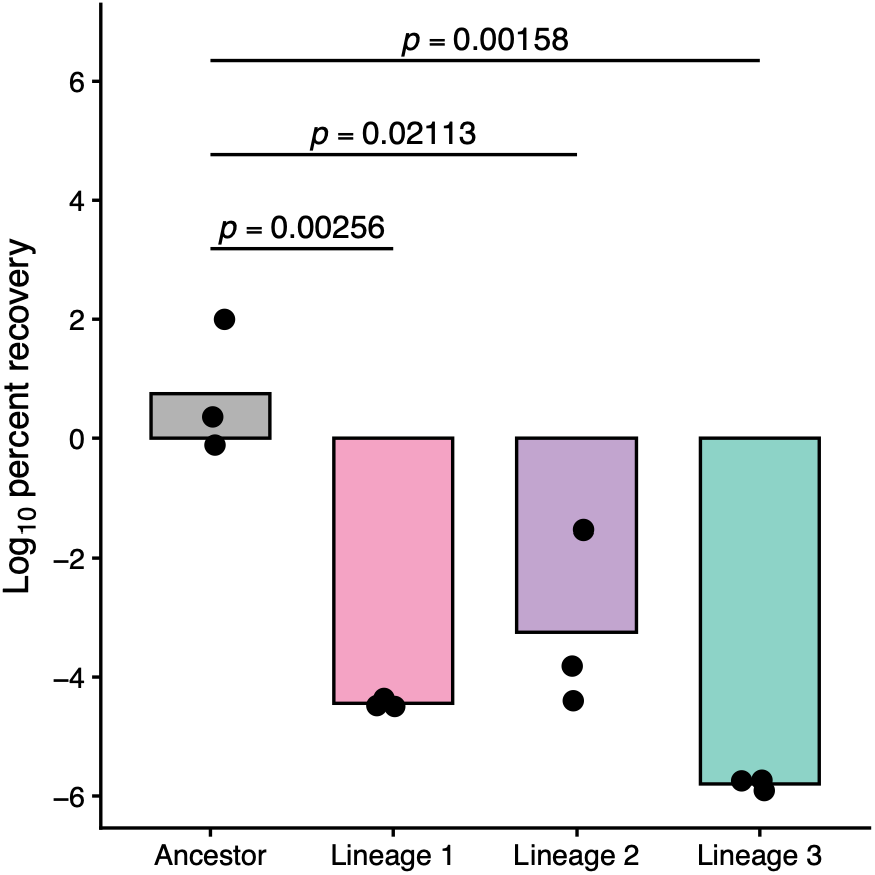
AZT-evolved KPPR1 lineages exhibit collateral sensitivity to TMP *in vitro*. Ancestral KPPR1 and AZT-evolved lineages were grown overnight at 37 °C with shaking (220 rpm) and subsequently cultured with TMP at 524 µg/mL. Following overnight TMP exposure, samples were plated on LB rifampicin (30 µg/mL) for CFU enumeration and determination of percent recovery. Percent recovery values were log_10_-transformed prior to analysis. Individual points represent independent biological replicates, and bars indicate the mean. Statistical comparisons were performed between each AZT-evolved lineage and the ancestral strain using Student’s *t*-tests with Holm correction for multiple comparisons; adjusted *p*-values are shown.

## Discussion

In this study, we investigate the potential use of antiviral drugs against Kp. We demonstrate that the novel anti-Kp drug candidates NH125, fingolimod, and AZT are bactericidal *in vitro* against hypervirulent KPPR1 and exhibit differential activity against a panel of clinical Kp isolates. Additionally, we show that AZT is active against Kp in settings that mimic gut colonization. Conversely, NH125 and fingolimod lost activity in these conditions, likely because of their dependency on biological context (fecal material source, oxygen, etc.). Overall, these findings support antiviral drugs as a promising strategy for controlling Kp and justify further evaluation in Kp-colonized or infected individuals.

Future research should explore the broader potential of repurposed antiviral drugs as antimicrobial agents beyond their activity against Kp, including their ability to inhibit multiple microbial species and potentially function as broad-spectrum or “pan-inhibitor” compounds. For example, AZT has been shown to inhibit other members of the Enterobacteriaceae family, including *Escherichia coli*, suggesting possible utility against organisms involved in co-infections and polymicrobial colonization.^19,29–31^ Extending these findings into *in vivo* models will be important to evaluate host and microbiome off-target effects, including toxicity, pharmacokinetics, route of administration, dosing, timing, and mechanisms of action, all of which are necessary to assess safety and translational potential. More broadly, this work highlights the promise of drug repurposing as a rapid strategy for identifying anti-Kp activity from FDA-approved compounds, potentially accelerating clinical translation while expanding the antimicrobial arsenal. In the context of increasing antimicrobial resistance, antiviral repurposing represents one of several complementary approaches that may help reduce reliance on traditional antibiotic discovery and provide alternative strategies to prevent or reduce Kp colonization (i.e., decolonization) and progression to invasive infection.

AZT is a synthetic thymidine analogue and was the first FDA-approved nucleos(t)ide reverse transcriptase inhibitor (NRTI) for the treatment of HIV.^32,33^ As an inactive nucleoside analogue, AZT requires sequential phosphorylation by host cellular kinases to generate the mono-, di-, and ultimately triphosphate metabolite, the active form responsible for inhibiting DNA synthesis.^34^ Following oral administration, AZT is rapidly absorbed, with an oral bioavailability of approximately 60–75%, which is lower than that of other commonly used NRTIs such as emtricitabine (∼93%)^35^, suggesting that a relatively large fraction of orally administered AZT remains within the gastrointestinal tract before systemic absorption.^36,37^ This may prove beneficial, as poorly absorbed antibiotics can reach high concentrations within the intestinal lumen and mucus, increasing local antimicrobial exposure.^38^ Although AZT has a relatively short plasma half-life of ∼1 hour, the intracellular half-life of its active triphosphate metabolite is substantially longer, with reported estimates ranging from ∼3–7 hours, contributing to prolonged intracellular exposure to the active metabolite.^36,39^ Clinically, AZT is no longer a first-line therapy for most adults/adolescents with HIV due to toxicity concerns and the development of newer antiretroviral agents,^40,41^ but it remains in use in specific contexts such as preventing mother-to-child HIV transmission and as part of neonatal prophylaxis regimens. Future studies should further evaluate the antibacterial activity of AZT across clinically relevant infection models and microbiome contexts, while also assessing whether structural modification or combination therapy could enhance its efficacy and reduce toxicity for potential repurposing beyond its current antiviral use.

The antimicrobial activity of AZT against members of the Enterobacteriaceae family suggests that its mechanism of nucleotide analog incorporation may have broader effects across bacterial DNA replication pathways, although these effects are not fully understood. Functional analysis of the AZT-susceptibility mutants identified in the transposon screen provides insight into the bacterial pathways that influence this response. KEGG BRITE and PANTHER GO analyses identified DNA-associated functions, carbohydrate/glycan metabolism, and cell envelope biosynthesis as major functional themes among the susceptibility mutants. The enrichment of DNA repair pathways and the GO molecular function term single-stranded DNA binding is consistent with ATZ’s activity as a thymidine analog and supports the hypothesis that disruption of DNA maintenance increases susceptibility to AZT-induced DNA stress. Likewise, the identification of exopolysaccharide synthesis, peptidoglycan biosynthesis, and glycan metabolism suggests that pathways dependent on nucleotide-activated precursors may also influence susceptibility to AZT. Interestingly, mutants with defects in nitrogen metabolism, fatty acid biosynthesis, and biotin metabolism had reduced susceptibility to AZT. One possible explanation of this finding may be that disruption of nucleotide synthesis and slow DNA replication decrease the demand for thymidine incorporation. Similarly, alterations in fatty acid biosynthesis or biotin metabolism may shift cellular metabolism or growth state in a manner that reduces susceptibility to AZT. Although these mechanisms remain speculative, these findings suggest that bacterial DNA maintenance and nucleotide-dependent metabolic pathways play important roles in determining the antibacterial activity of AZT in Kp.

Gut colonization by Kp represents an attractive target for infection prevention because colonized patients are at increased risk for subsequent extraintestinal infection, which is frequently caused by the strain already present in the gastrointestinal tract.^6^ Selective digestive decontamination with poorly absorbed antibiotics such as gentamicin has therefore been explored for patients colonized with carbapenem-resistant Kp; however, clinical studies have demonstrated incomplete or transient eradication and, importantly, the emergence of gentamicin- and colistin-resistant isolates during decolonization, highlighting resistance development as a major limitation of antibiotic-based approaches.^42,43^ Our finding that AZT retains activity against Kp in an *ex vivo* human fecal environment supports its potential repurposing as an alternative gut decolonization strategy, but the emergence of AZT resistance similarly necessitates consideration of evolutionary escape. Our identification of *tdk* mutations following AZT evolution suggests that loss of thymidine kinase activity represents an accessible resistance pathway in Kp. Importantly, loss of Tdk also eliminates thymidine salvage, potentially increasing dependence on folate-dependent *de novo* thymidylate synthesis and thereby creating a collateral vulnerability to TMP. This potential collateral sensitivity is further supported by previous work demonstrating synergy between AZT and TMP, in which TMP-mediated disruption of folate biosynthesis and nucleotide homeostasis sensitized bacteria to AZT-mediated inhibition of DNA replication.^44^ This interaction was also observed in multidrug-resistant clinical Kp isolates, supporting the potential relevance of combining AZT with TMP against Kp. This interaction may be particularly relevant clinically because TMP-sulfamethoxazole (TMP-SMX) is already routinely administered for opportunistic-infection prophylaxis in immunocompromised populations, including solid-organ and hematopoietic-cell transplant recipients, who also represent important populations at risk for Kp colonization and invasive infection.^45,46^ Thus, rather than serving simply as a second decolonizing agent, TMP-SMX could provide a clinically established partner for AZT-based decolonization that exploits the metabolic liability associated with AZT resistance. Determining whether AZT remains active against TMP-evolved populations, together with the increased TMP susceptibility predicted for AZT-evolved *tdk* mutants, will establish whether this combination not only enhances antibacterial activity but also constrains accessible evolutionary pathways to resistance.

In contrast to AZT, fingolimod and NH125 lost activity in our gut relevant settings. Nonetheless, these drugs may be useful in other infectious settings, specifically those with aerobic states, such as pneumonia. Fingolimod is FDA-approved for the treatment of multiple sclerosis and stroke through reducing circulating lymphocytes by trapping them in secondary lymphoid tissues; however, studies are looking into repurposing the use of fingolimod as an antiviral treatment to block viral entry into cells specifically in the context of HIV.^47,48^ NH125 is not FDA approved, but current studies are aiming to utilize this drug for cancer therapy by modulating various R groups to lower toxicity. Others are looking into the possible effects it has on blocking viral entry specifically for avian influenza virus (H5N1), Ebola virus, and Lassa virus.^49–52^ While the mechanism of action in terms of viral elimination is either known or being studied for these drugs, their bacterial targets remain unknown. Future studies should investigate the mechanistic basis of their antibacterial effects under infection-relevant conditions and determine whether chemical modification or combination therapy could restore or enhance their activity in gut-associated environments.

We acknowledge that our study has several limitations warrant consideration. There are thousands of potentially antibacterial compounds available, and we screened only 509; therefore, the six candidates identified here do not represent a comprehensive assessment. Nevertheless, these findings lay promising groundwork for future directions in the field, particularly given that multiple targets were identified. As mentioned throughout, urine was the most sampled site for the array of Kp clinical isolates; therefore, future studies incorporating larger and more diverse clinical cohorts, as well as clinical metadata, will be important to strengthen the rigor and interpretability of these findings. Finally, off-target effects of these drugs, including elimination of beneficial bacteria in the gut, are plausible and should be addressed in future studies through careful assessment of microbiome composition changes using sequencing-based approaches. Despite these limitations, the findings presented here demonstrate, for the first time, the feasibility of drug repurposing as an anti-Kp therapeutic strategy, with further clinical evaluation needed before translation to approved use.

## Methods

### Bacterial strains

For initial drug susceptibility screening, hypervirulent KPPR1pJL1-sfGFP was constructed with a pfL1-sfGFP plasmid (Addgene item 102634, electroporated into KPPR1) and then grown overnight from a single fluorescent colony in Luria-Bertani (LB, Miller formulation) supplemented with 40 µg/mL of kanamycin. Kp rifampin-resistant mutant, KPPR1^53^, was initially grown on LB agar overnight at 27 °C with 30 µg/mL of rifampicin. All materials and reagents were purchased from Sigma-Aldrich or Thermo Fisher Scientific unless otherwise specified.

### KPPR1 susceptibility screening

An overnight culture of KPPR1pJL1-sfGFP was prepared as described, then diluted to a final OD_600_ = 0.01. This culture was arrayed into 96-well plates, then each compound from the DiscoveryProbe™ Anti-virus Compound Library (APExBIO, Catalog No. L1050) were added to individual wells to a final concentration of 1 mM. Cultures were incubated at 37 °C with shaking (220 rpm) for 24 hours. Resulting cultures were transferred to black-walled 96 well plates, and growth was determined by measuring OD_600_ and relative fluorescent units (RFU). This screen was independently repeated three times. Then, growth was summarized as a *z*-score for each biological replicate. Growth-restrictive compounds were identified as having a calculated OD_600_ *z*-score below −2.5 and a RFU *z*-score below −1 compared to the entire 509 compound library in each of the three replicates. Resulting “hits” were then selected to proceed forward with MIC/MBC determination. Crystal violet was excluded from our hit list due to optical interference.

### MIC/MBC determination

An overnight culture of KPPR1 was prepared in LB broth and then diluted a final OD_600_ = 0.01. Antiviral drugs were initially reconstituted in DMSO, then diluted to a final concentration of 400 µM in LB broth, then subjected to ten two-fold dilutions. In a 96 well round bottom plate, KPPR1 and antiviral drugs were added. Cultures were incubated at 37 °C with shaking (220 rpm) for 24 hours. A no-growth blank control of LB with 4% DMSO and a KPPR1 only (no drug) were included. The minimum inhibitory concentration (MIC) was determined visually, then the minimum bactericidal concentration (MBC) was determined by quantifying Kp viability via serial plating two dilutions above and below the determined MIC and the controls on LB agar.

### Collection and preparation of clinical isolates

Clinical isolates were collected and de-identified by in accordance with approval by the Indiana University Institutional Review Board (protocol #16139, PI: Ryan F. Relich). These isolates were derived from a variety of infection sites (blood, urine, wound [abscess, pilonidal drainage], respiratory tract [tracheal aspirate, sputum, bronchoalveolar lavage fluid, bronchial wash], or other) and samples were de-identified. Isolates were passaged once from primary plating media (sheep blood agar or MacConkey agar) prior to storage on brain-heart infusion agar slants (ThermoFisher) at room temperature.

### Clinical isolates screening and validation

Collected Kp clinical isolates were cultured and subsequently frozen in a 96 well plate to create an array to be used for all subsequent experiments. Next, this array was cultured in LB broth in a 96 deep well plate using a pin replicator at 37 °C with shaking (220 rpm) overnight. The following day samples were mixed well, diluted 1:10, and diluted again 1:100 into a new 96 well round bottom plate supplemented with the following drugs at twice their pre-determined MIC: 3.125 µM gentamicin, 12.5 µM AZT, 12.5 µM NH125, and 100 µM fingolimod. Controls included a blank well in each plate that received no bacteria, one well in each plate with KPPR1 only, and a control plate of LB broth only. If growth was observed in the KPPR1 only well, then the entire plate was excluded from analysis. After overnight incubation at 37 °C shaking, plates were visually read as either sensitive (no growth) or resistant (growth). For validation, three Kp clinical isolates that varied in their sensitivity/resistance and their collection site were selected: JV94 - urine, JV101 - lung, and JV132 - blood, and their MICs were determined for gentamicin, AZT, NH125, and fingolimod as described above.

### Detection of *rmpA*

PCR was performed using primers rmpA_Fwd (5’-GAG TAT TGG TTG ACA GCA GGA - 3’) and rmpA_Rev (5’-AGC CGT GGA TAA TGG TTT ACA −3’). Thermocycler settings included an initial denaturation at 95 °C for 2 min., followed by 35 cycles at 95 °C for 15 sec., 59 °C for 15 sec., and 72 °C for 30 sec., with a final extension at 72 °C for 5 min. A 1.5% agarose gel with SYBR safe gel stain was cast and run at 130 volts for 30 min. then visualized using a UV light source to determine the presence or absence of a band. Positive PCR results were confirmed by string test.^54^

### Aerobic/Anaerobic *ex vivo* human fecal microbiota assays

Three male human fecal samples were obtained from Medix Biochemica USA Inc. (lot #’s 25-02-744, 25-02-770, and 25-08-1065). Each sample was weighed and diluted 1:10 in anaerobically reduced 1X PBS. Samples were then inoculated with ∼10^5^ CFU KPPR1 and 16X the MIC of gentamicin, AZT, NH125, or fingolimod. Cultures were incubated at 37 °C for 48 hours under either static anaerobic conditions in a Coy Laboratories type B vinyl anaerobic chamber or aerobic conditions with shaking (220 rpm), and KPPR1 was quantified by dilution plating on selective media.

### *In vitro* transposon screen for AZT susceptibility

An in-depth description of the condensed, arrayed KPPR1 transposon library construction can be found elsewhere.^55^ Arrayed transposon mutants were pin-replicated into deep-well plates containing 2 mL of LB supplemented with 30 µg/mL rifampicin and ¼ of the predetermined MIC (1.56 µM) of AZT. Cultures were incubated for 24 hrs at 37 °C with shaking at 220 rpm. Plates were initially scored for growth or no growth, after which cultures were sampled for OD_600_ measurements using a plate reader. VK055 gene locus tags from mapped transposon mutants were used to assign genes to Kyoto Encyclopedia of Genes and Genomes (KEGG) identifiers and KEGG BRITE functional hierarchy annotations.^56,57^ Growth phenotypes were normalized within each trial by calculating *z*-scores from blank OD_600_ values, where negative *z*-scores indicated reduced growth compared with the trial mean.

KEGG BRITE annotations were joined to the *z*-score dataset, and mutant phenotypes were summarized by Level C category. To avoid counting the same gene more than once within a category, duplicate gene-category combinations were removed. For each Level C category, the number of unique genes and mean *z*-score was calculated. Categories with fewer than two unique genes were excluded from the main summary, as were *z*-score outliers (Grubbs’ test, N = 1), and categories with an absolute mean *z*-score ≥ 0.3 were considered candidate functional groups associated with altered growth during AZT exposure.

For PANTHER GO analysis, mutants were classified as potential AZT-susceptibility hits if they had a trial-level *z*-score below −1.4 in all trials tested. These hit genes were mapped to putative *E. coli* orthologs using shared KEGG Orthology identifiers. The corresponding *E. coli* UniProt identifiers were then analyzed using the PANTHER Overrepresentation Test with *E. coli* all genes in the PANTHER database as the reference list. GO biological process complete, GO molecular function complete, and GO cellular component complete datasets were tested separately using Fisher’s exact test with false discovery rate correction. Terms with FDR ≤ 0.05 and positive fold enrichment were considered significantly overrepresented.

### KPPR1 evolution in AZT

Three independent KPPR1 colonies were selected for parallel evolution in the presence of ¼ (5 passages) then ½ (5 passages) of the KPPR1 MIC of AZT. Cultures were maintained in biological triplicate throughout the experiment. After overnight growth, aliquots of each culture were preserved as glycerol stocks at −80°C. In parallel, each biological replicate was drip plated onto LB agar supplemented with 30 µg/mL rifampicin to monitor colony morphology and culture purity. To initiate the subsequent passage, a 1:1,000 dilution of each culture was transferred into fresh LB containing 30 µg/mL rifampicin and AZT. Each transfer was considered one passage, and this serial passaging procedure was repeated for a total of 10 passages.

### Genomic DNA extraction, whole genome sequencing, and mutation identification

AZT-evolved strains were cultured overnight in LB broth prior to genomic DNA extraction. Genomic DNA was extracted using the Monarch Genomic DNA Purification Kit (New England Biolabs, Ipswich, MA) according to the manufacturer’s instructions. DNA concentration was quantified using a Qubit 4 Fluorometer with the Qubit 1x dsDNA High Sensitivity Assay Kit (Thermo Fisher Scientific, Waltham, MA). For each sample, 200 ng of genomic DNA was prepared for sequencing using the Rapid Barcoding Kit 24 V14 (SQK-RBK114.24; Oxford Nanopore Technologies, Oxford, England) according to the manufacturer’s Rapid sequencing DNA V14 barcoding protocol. Bacterial genomes were sequenced on a FLO-MIN114 flow cell using a MinION Mk1B device (Oxford Nanopore Technologies, Oxford, England). Fast basecalling and barcode trimming were performed during sequencing using MinKNOW software (v26.01.15). Sequencing was continued until a minimum of 100,000 reads per barcode was obtained. breseq^58^ was to identify mutations in AZT-evolved lineages. The KPPR1 reference genome was downloaded from The National Center for Biotechnology Information (GenBank accession CP009208.1), and ONT Nanpore reads were used to identify mutations using the “-x” and “--no-junction-prediction” options.

### Trimethoprim (TMP) killing assay

Ancestral KPPR1 and AZT-evolved KPPR1 lineages were grown overnight at 37 °C with shaking (220 rpm). Overnight cultures were standardized to a final OD_600_ of 0.01 into fresh LB broth with or without TMP at a final concentration of 524 µg/mL. Following overnight incubation, cultures were serially diluted and plated on LB agar containing rifampicin (30 µg/mL) for CFU enumeration. CFU recovery following TMP treatment was compared with the corresponding untreated LB control to determine percent recovery.

### Statistical analysis

All *in vitro* experimental replicates represent biological replicates. A *p*-value ≤ 0.05 was considered statistically significant for the above experiments unless otherwise indicated, and analysis was performed using base R version 2026.04.0+526. Data were processed, analyzed, and visualized using R packages “readxl,” “tidyverse,” “dplyr,” “vegan,” “EnvStats,” “ggplot2,” “ggrepel,”“ggtext.,” “writexl”, “KEGGREST”, “tibble”, “stringr”, and “outliers”.^59–71^

### Data availability

All raw data and analysis scripts used for this study are available at https://github.com/jayvorn/antiviral-drug-repurposing-for-Klebsiella-peumoniae. Raw sequencing data are available on the Sequence Read Archive (Bioproject PRJNA1525623).

## Acknowledgments

The authors would like to acknowledge the members of the Tilston and Vornhagen labs for their feedback on this study.

## Contributions

Conceptualization: NLA, JV

Methodology: NLA, JV

Investigation: All authors

Visualization: NLA, JV

Funding acquisition: NLT, JV

Project administration: RR, JV

Supervision: JV

Writing – original draft: NLA, KG, JV

Writing – review & editing: All authors

## Use of AI

ChatGPT was used after drafting this manuscript for grammar and clarity improvements.

## Supplemental Data

**Supplemental Table 1.** Summary of clinical *Klebsiella pneumoniae* isolates.

| Strain Number | Collection Site | Strain Number | Collection Site |
| --- | --- | --- | --- |
| 74 | Urine | 123 | Urine |
| 75 | Urine | 124 | Urine |
| 76 | Urine | 125 | Urine |
| 77 | Urine | 126 | Urine |
| 78 | Urine | 127 | Sputum |
| 79 | Urine | 128 | Urine |
| 80 | Urine | 129 | Sputum |
| 81 | Urine | 130 | Urine |
| 82 | Urine | 131 | Blood |
| 83 | Urine | 132 | Blood |
| 84 | Urine | 134 | Urine |
| 86 | Urine | 135 | Urine |
| 88 | Urine | 136 | Urine |
| 89 | Urine | 137 | Unknown |
| 90 | Urine | 138 | Urine |
| 91 | Urine | 139 | Urine |
| 92 | Urine | 140 | Urine |
| 93 | Urine | 141 | Blood |
| 94 | Urine | 142 | Blood |
| 95 | Urine | 143 | Urine |
| 96 | Urine | 144 | Urine |
| 97 | Urine | 145 | Urine |
| 98 | Urine | 146 | Blood |
| 100 | Tracheal Aspirate | 147 | Urine |
| 101 | BALF* | 148 | Tracheal Aspirate |
| 102 | Pilonidal Drainage | 149 | Urine |
| 103 | Tracheal Aspirate | 150 | Sputum |
| 104 | Urine | 151 | Blood |
| 105 | Urine | 152 | BALF* |
| 106 | Urine | 153 | Urine |
| 107 | Urine | 154 | Blood |
| 109 | Urine | 155 | Urine |
| 112 | Sputum | 156 | Urine |
| 113 | Urine | 157 | Urine |
| 114 | Urine | 158 | Urine |
| 115 | Urine | 159 | Urine |
| 116 | Urine | 160 | Sputum |
| 117 | Bronchial Wash | 161 | Abscess |
| 118 | Urine | 162 | Tracheal Aspirate |
| 119 | Tracheal Aspirate | 163 | Urine |
| 120 | Urine | 165 | Urine |
| 121 | Tracheal Aspirate | 166 | Urine |
| 122 | Blood | 167 | Urine |
|  |  | 169 | Urine |

\*Bronchoalveolar lavage fluid

**Supplemental Table 2.** MICs of clinical Kp isolates.

| Clinical isolate | Gentamicin MIC (μM)* | AZT MIC (μM)* | NH125 MIC (μM)* | Fingolimod MIC (μM)* |
| --- | --- | --- | --- | --- |
| JV94 | 1.56 (S) | 6.25 (R) | 200 (R) | 50 (S) |
| JV101 | 3.13 (S) | >400 (R) | 50 (R) | 25 (S) |
| JV132 | 6.25 (S) | >400 (R) | 12.5 (S) | 50 (S) |

\*Screen results indicated in parentheses. A result is considered valid if the MIC is ± 2-fold the screen concentration (gentamicin = 3.13 μM, AZT = 12.5 μM, NH125 = 12.5 μM, fingolimod = 100 μM).

**Supplemental Tabel 3.** *z*-scores of Kp transposon mutants.

| Locus Tag | z-score | Locus Tag | z-score | Locus Tag | z-score | Locus Tag | z-score |
| --- | --- | --- | --- | --- | --- | --- | --- |
| VK055_0011 | -0.885 | VK055_1331 | -0.189 | VK055_2629 | -0.808 | VK055_3920 | -0.290 |
| VK055_0013 | -0.295 | VK055_1332 | -0.735 | VK055_2630 | 1.076 | VK055_3921 | -0.245 |
| VK055_0016 | 0.034 | VK055_1333 | 0.143 | VK055_2631 | -0.379 | VK055_3922 | 0.127 |
| VK055_0017 | 0.427 | VK055_1334 | -0.142 | VK055_2632 | 1.660 | VK055_3923 | -0.286 |
| VK055_0018 | 0.175 | VK055_1335 | 0.505 | VK055_2633 | 0.109 | VK055_3924 | 0.189 |
| VK055_0019 | 0.562 | VK055_1337 | 0.163 | VK055_2634 | 0.148 | VK055_3925 | 0.417 |
| VK055_0020 | 0.200 | VK055_1340 | -0.753 | VK055_2635 | 0.330 | VK055_3926 | 0.259 |
| VK055_0023 | -0.538 | VK055_1341 | -0.673 | VK055_2636 | -0.835 | VK055_3927 | 0.374 |
| VK055_0024 | 1.704 | VK055_1342 | 0.374 | VK055_2637 | -0.831 | VK055_3928 | -0.306 |
| VK055_0025 | 0.502 | VK055_1344 | 0.330 | VK055_2638 | 0.671 | VK055_3929 | 0.110 |
| VK055_0026 | 0.029 | VK055_1345 | -0.975 | VK055_2639 | 0.797 | VK055_3930 | -1.470 |
| VK055_0027 | -0.731 | VK055_1347 | -1.490 | VK055_2640 | -1.397 | VK055_3931 | -0.516 |
| VK055_0028 | 0.108 | VK055_1351 | -0.287 | VK055_2641 | 0.297 | VK055_3933 | -1.186 |
| VK055_0029 | -0.156 | VK055_1352 | 0.261 | VK055_2642 | 1.099 | VK055_3935 | -0.809 |
| VK055_0032 | -0.065 | VK055_1353 | -1.016 | VK055_2643 | -1.471 | VK055_3936 | 0.136 |
| VK055_0033 | -0.505 | VK055_1354 | 1.658 | VK055_2644 | 0.952 | VK055_3937 | -1.105 |
| VK055_0038 | 0.205 | VK055_1355 | -1.055 | VK055_2645 | -0.315 | VK055_3938 | 0.474 |
| VK055_0039 | 0.179 | VK055_1356 | -0.618 | VK055_2646 | -0.144 | VK055_3939 | 0.283 |
| VK055_0040 | 0.999 | VK055_1357 | -0.172 | VK055_2649 | -0.514 | VK055_3941 | -0.623 |
| VK055_0042 | -0.433 | VK055_1358 | -0.031 | VK055_2650 | -0.720 | VK055_3942 | -0.912 |
| VK055_0044 | -1.268 | VK055_1359 | 0.310 | VK055_2651 | -0.854 | VK055_3944 | -0.489 |
| VK055_0045 | 0.277 | VK055_1360 | 0.995 | VK055_2652 | -0.989 | VK055_3945 | 0.481 |
| VK055_0046 | -1.186 | VK055_1361 | 0.501 | VK055_2653 | -0.376 | VK055_3946 | 0.477 |
| VK055_0047 | 0.139 | VK055_1362 | 0.042 | VK055_2655 | -0.521 | VK055_3947 | 0.782 |
| VK055_0049 | 0.954 | VK055_1363 | -0.647 | VK055_2656 | 1.733 | VK055_3948 | 0.042 |
| VK055_0051 | 0.590 | VK055_1364 | -1.476 | VK055_2657 | 0.440 | VK055_3949 | 0.018 |
| VK055_0052 | 0.029 | VK055_1365 | 0.426 | VK055_2659 | 0.952 | VK055_3951 | 0.737 |
| VK055_0053 | 0.182 | VK055_1366 | -0.435 | VK055_2660 | -1.434 | VK055_3952 | -0.291 |
| VK055_0054 | 0.253 | VK055_1367 | -1.215 | VK055_2661 | 1.357 | VK055_3953 | -0.975 |

| Locus Tag | Z-score | Locus Tag | Z-score | Locus Tag | Z-score | Locus Tag | Z-score |
| --- | --- | --- | --- | --- | --- | --- | --- |
| VK055_0055 | 0.648 | VK055_1370 | 0.518 | VK055_2662 | 0.363 | VK055_3954 | 0.560 |
| VK055_0056 | -0.279 | VK055_1371 | -1.080 | VK055_2663 | 1.107 | VK055_3955 | 0.513 |
| VK055_0057 | -0.168 | VK055_1372 | 2.114 | VK055_2664 | -0.648 | VK055_3956 | 0.943 |
| VK055_0058 | 0.169 | VK055_1380 | -1.443 | VK055_2665 | 0.276 | VK055_3957 | -1.483 |
| VK055_0059 | -0.486 | VK055_1381 | 0.337 | VK055_2666 | 0.356 | VK055_3958 | 0.287 |
| VK055_0060 | 0.260 | VK055_1382 | 0.302 | VK055_2667 | 1.248 | VK055_3959 | 0.135 |
| VK055_0061 | 0.072 | VK055_1384 | 0.509 | VK055_2668 | -0.153 | VK055_3960 | 0.209 |
| VK055_0062 | -0.162 | VK055_1387 | 0.525 | VK055_2669 | -1.434 | VK055_3961 | 0.450 |
| VK055_0063 | -0.748 | VK055_1388 | -0.075 | VK055_2670 | 1.752 | VK055_3963 | -0.063 |
| VK055_0064 | -0.761 | VK055_1389 | 0.232 | VK055_2671 | -0.559 | VK055_3964 | 0.300 |
| VK055_0066 | 0.350 | VK055_1390 | -1.477 | VK055_2673 | -1.476 | VK055_3965 | 0.455 |
| VK055_0073 | -0.784 | VK055_1392 | -0.273 | VK055_2674 | 0.329 | VK055_3966 | 0.003 |
| VK055_0074 | -1.476 | VK055_1393 | 0.322 | VK055_2675 | -0.286 | VK055_3967 | -0.685 |
| VK055_0075 | 0.351 | VK055_1395 | 0.235 | VK055_2676 | 0.632 | VK055_3968 | -0.014 |
| VK055_0076 | 0.762 | VK055_1396 | 0.130 | VK055_2677 | -0.362 | VK055_3969 | 0.033 |
| VK055_0077 | 0.090 | VK055_1397 | 0.062 | VK055_2678 | -0.629 | VK055_3970 | 0.865 |
| VK055_0078 | 0.490 | VK055_1399 | 0.620 | VK055_2681 | 0.756 | VK055_3971 | -0.656 |
| VK055_0079 | 0.742 | VK055_1400 | -0.367 | VK055_2682 | -0.472 | VK055_3973 | -0.934 |
| VK055_0081 | 0.879 | VK055_1401 | 0.247 | VK055_2685 | -1.347 | VK055_3975 | -1.315 |
| VK055_0082 | 0.286 | VK055_1402 | 0.098 | VK055_2686 | 0.400 | VK055_3976 | -0.733 |
| VK055_0084 | -1.054 | VK055_1403 | -0.143 | VK055_2687 | 0.101 | VK055_3977 | 1.163 |
| VK055_0086 | -1.434 | VK055_1404 | -1.462 | VK055_2688 | 0.142 | VK055_3978 | 1.282 |
| VK055_0087 | 0.176 | VK055_1405 | -0.786 | VK055_2689 | -1.051 | VK055_3979 | 1.042 |
| VK055_0088 | 0.282 | VK055_1407 | 0.925 | VK055_2690 | 0.513 | VK055_3981 | -0.587 |
| VK055_0089 | 0.280 | VK055_1408 | 0.721 | VK055_2691 | 0.508 | VK055_3982 | 0.286 |
| VK055_0091 | -0.011 | VK055_1410 | -0.557 | VK055_2692 | -0.666 | VK055_3983 | 0.324 |
| VK055_0095 | -0.510 | VK055_1411 | 0.017 | VK055_2693 | 0.100 | VK055_3984 | 0.766 |
| VK055_0096 | 0.475 | VK055_1413 | 1.242 | VK055_2694 | 0.298 | VK055_3985 | -0.312 |
| VK055_0097 | 1.582 | VK055_1414 | -0.678 | VK055_2695 | -1.497 | VK055_3989 | 0.128 |
| VK055_0098 | 0.156 | VK055_1417 | -0.013 | VK055_2700 | -0.870 | VK055_3990 | -0.478 |
| VK055_0099 | 1.721 | VK055_1419 | 0.451 | VK055_2701 | 0.429 | VK055_3991 | -0.678 |
| VK055_0100 | -0.283 | VK055_1420 | 0.095 | VK055_2702 | 1.921 | VK055_3999 | -0.830 |
| VK055_0101 | -1.491 | VK055_1424 | -0.211 | VK055_2703 | 0.318 | VK055_4000 | 0.130 |
| VK055_0103 | -0.217 | VK055_1425 | 0.989 | VK055_2704 | 0.115 | VK055_4001 | 0.251 |
| VK055_0104 | -1.184 | VK055_1426 | 0.131 | VK055_2705 | 0.200 | VK055_4002 | 0.157 |
| VK055_0105 | 0.029 | VK055_1427 | -0.271 | VK055_2706 | -0.677 | VK055_4003 | 0.311 |
| VK055_0107 | -0.587 | VK055_1428 | 0.392 | VK055_2707 | -0.069 | VK055_4004 | 0.369 |
| VK055_0109 | -0.363 | VK055_1429 | -0.748 | VK055_2708 | -1.464 | VK055_4006 | -1.483 |
| VK055_0111 | 0.045 | VK055_1430 | -0.192 | VK055_2709 | 0.548 | VK055_4009 | 1.021 |
| VK055_0112 | -0.371 | VK055_1431 | -0.342 | VK055_2710 | 0.596 | VK055_4010 | -1.081 |
| VK055_0114 | -0.198 | VK055_1432 | -0.525 | VK055_2711 | 0.142 | VK055_4011 | 0.035 |
| VK055_0115 | -0.490 | VK055_1433 | -0.562 | VK055_2712 | -1.434 | VK055_4013 | 0.711 |
| VK055_0117 | 0.874 | VK055_1434 | -0.355 | VK055_2714 | 0.307 | VK055_4016 | 0.918 |
| VK055_0118 | -1.282 | VK055_1436 | 0.561 | VK055_2715 | 0.399 | VK055_4020 | -1.478 |
| VK055_0119 | -0.650 | VK055_1437 | 0.168 | VK055_2716 | -0.101 | VK055_4021 | 0.545 |
| VK055_0120 | 0.774 | VK055_1438 | -0.947 | VK055_2717 | 0.859 | VK055_4022 | 0.612 |
| VK055_0121 | 0.679 | VK055_1439 | -0.123 | VK055_2718 | 0.402 | VK055_4023 | -0.891 |
| VK055_0122 | 0.458 | VK055_1440 | 0.110 | VK055_2722 | -1.485 | VK055_4024 | -0.355 |
| VK055_0123 | -0.350 | VK055_1441 | 0.095 | VK055_2723 | -1.497 | VK055_4026 | 0.092 |
| VK055_0124 | -0.344 | VK055_1443 | -0.316 | VK055_2724 | -0.127 | VK055_4027 | 1.610 |
| VK055_0125 | -1.497 | VK055_1445 | -0.218 | VK055_2725 | 0.409 | VK055_4028 | 0.598 |
| VK055_0127 | -0.474 | VK055_1446 | 0.060 | VK055_2726 | -0.885 | VK055_4029 | -0.258 |
| VK055_0128 | -0.667 | VK055_1448 | -0.286 | VK055_2727 | 0.230 | VK055_4030 | 0.476 |
| VK055_0129 | 0.758 | VK055_1449 | -0.697 | VK055_2728 | -0.750 | VK055_4032 | -1.446 |
| VK055_0131 | 0.504 | VK055_1450 | 0.276 | VK055_2729 | -0.978 | VK055_4033 | 1.196 |
| VK055_0132 | 0.071 | VK055_1451 | 0.004 | VK055_2730 | -0.885 | VK055_4034 | 0.253 |
| VK055_0134 | 0.128 | VK055_1453 | 1.705 | VK055_2732 | 0.327 | VK055_4035 | 0.004 |
| VK055_0136 | 0.820 | VK055_1454 | 1.704 | VK055_2733 | -0.348 | VK055_4036 | 1.116 |
| VK055_0138 | -0.017 | VK055_1455 | 0.327 | VK055_2735 | -0.144 | VK055_4039 | 0.942 |
| VK055_0139 | 0.844 | VK055_1456 | 0.288 | VK055_2736 | 0.878 | VK055_4040 | -0.148 |
| VK055_0140 | -0.572 | VK055_1457 | 0.631 | VK055_2737 | -0.755 | VK055_4041 | -0.553 |
| VK055_0141 | -0.294 | VK055_1458 | -0.274 | VK055_2738 | -0.903 | VK055_4042 | 2.474 |
| VK055_0143 | -1.072 | VK055_1459 | 0.309 | VK055_2739 | -1.194 | VK055_4043 | 0.719 |
| VK055_0144 | -1.063 | VK055_1460 | 0.149 | VK055_2740 | -0.758 | VK055_4044 | 0.075 |
| VK055_0145 | -0.625 | VK055_1461 | 0.450 | VK055_2741 | -0.669 | VK055_4045 | -0.185 |
| VK055_0147 | -0.206 | VK055_1463 | 0.871 | VK055_2742 | -0.912 | VK055_4046 | 0.005 |
| VK055_0148 | 0.478 | VK055_1464 | 0.280 | VK055_2743 | -0.800 | VK055_4048 | -0.098 |
| VK055_0149 | 0.372 | VK055_1465 | -0.497 | VK055_2744 | 0.041 | VK055_4050 | -0.424 |
| VK055_0150 | -1.442 | VK055_1466 | 1.494 | VK055_2746 | -0.951 | VK055_4051 | 0.682 |
| VK055_0151 | 0.322 | VK055_1467 | -0.242 | VK055_2748 | 0.082 | VK055_4052 | -1.155 |
| VK055_0152 | 0.223 | VK055_1468 | -0.667 | VK055_2749 | 1.478 | VK055_4053 | 0.704 |
| VK055_0153 | -1.338 | VK055_1469 | 0.052 | VK055_2750 | -0.282 | VK055_4054 | -1.476 |
| VK055_0155 | -0.836 | VK055_1470 | -0.577 | VK055_2752 | 0.350 | VK055_4055 | -1.440 |
| VK055_0156 | -0.278 | VK055_1471 | -1.242 | VK055_2753 | -1.435 | VK055_4056 | -0.385 |
| VK055_0157 | -0.646 | VK055_1472 | 0.365 | VK055_2754 | 0.130 | VK055_4057 | 1.021 |
| VK055_0161 | -1.172 | VK055_1473 | -0.530 | VK055_2759 | 0.253 | VK055_4058 | 1.091 |
| VK055_0162 | -0.017 | VK055_1474 | 0.608 | VK055_2765 | -0.092 | VK055_4059 | -0.343 |
| VK055_0164 | -0.074 | VK055_1475 | -0.350 | VK055_2767 | -0.852 | VK055_4062 | -0.493 |
| VK055_0165 | 1.915 | VK055_1476 | 0.201 | VK055_2768 | -0.370 | VK055_4064 | 0.451 |
| VK055_0166 | -0.767 | VK055_1477 | -1.440 | VK055_2769 | -0.444 | VK055_4065 | 0.003 |
| VK055_0167 | 0.256 | VK055_1478 | -0.829 | VK055_2771 | 1.489 | VK055_4066 | -0.507 |
| VK055_0168 | -1.056 | VK055_1479 | 0.297 | VK055_2773 | -0.356 | VK055_4067 | -1.291 |
| VK055_0169 | 0.053 | VK055_1480 | 0.103 | VK055_2774 | 0.261 | VK055_4068 | -0.278 |
| VK055_0170 | -0.131 | VK055_1481 | -0.342 | VK055_2775 | 0.394 | VK055_4073 | 0.539 |
| VK055_0171 | 0.250 | VK055_1482 | 0.411 | VK055_2776 | 0.038 | VK055_4074 | -0.394 |
| VK055_0172 | -0.224 | VK055_1483 | -1.305 | VK055_2777 | 0.793 | VK055_4075 | -0.690 |
| VK055_0173 | -0.342 | VK055_1485 | 0.085 | VK055_2778 | -0.084 | VK055_4077 | 0.501 |
| VK055_0174 | 1.089 | VK055_1486 | 0.075 | VK055_2779 | -0.685 | VK055_4078 | -0.511 |
| VK055_0175 | -1.396 | VK055_1487 | 0.278 | VK055_2780 | 0.106 | VK055_4079 | -0.175 |
| VK055_0176 | 0.259 | VK055_1488 | -0.066 | VK055_2781 | -0.779 | VK055_4080 | 1.076 |
| VK055_0178 | 0.706 | VK055_1490 | 1.150 | VK055_2783 | -0.706 | VK055_4082 | -0.167 |
| VK055_0179 | 0.823 | VK055_1491 | 0.064 | VK055_2787 | 0.858 | VK055_4083 | 0.113 |
| VK055_0180 | -0.486 | VK055_1492 | 0.053 | VK055_2788 | -1.180 | VK055_4084 | 0.192 |
| VK055_0182 | 0.184 | VK055_1494 | -0.524 | VK055_2790 | -0.158 | VK055_4086 | -0.392 |
| VK055_0183 | -0.204 | VK055_1495 | -1.114 | VK055_2794 | -1.459 | VK055_4088 | 0.862 |
| VK055_0184 | 0.007 | VK055_1497 | -0.065 | VK055_2797 | -0.362 | VK055_4089 | -0.043 |
| VK055_0186 | -0.374 | VK055_1498 | -0.992 | VK055_2798 | 1.096 | VK055_4091 | 0.642 |
| VK055_0187 | 0.084 | VK055_1500 | -1.462 | VK055_2799 | 0.021 | VK055_4092 | 0.083 |
| VK055_0188 | 1.058 | VK055_1501 | -0.044 | VK055_2800 | -0.515 | VK055_4093 | -0.686 |
| VK055_0189 | -1.009 | VK055_1503 | -0.074 | VK055_2801 | 0.145 | VK055_4094 | 0.561 |
| VK055_0191 | -1.199 | VK055_1505 | 0.162 | VK055_2802 | -1.135 | VK055_4096 | -0.415 |
| VK055_0193 | -0.181 | VK055_1506 | -1.114 | VK055_2803 | -0.630 | VK055_4099 | -0.418 |
| VK055_0194 | -0.188 | VK055_1507 | 0.823 | VK055_2804 | -1.147 | VK055_4100 | -0.483 |
| VK055_0195 | 0.166 | VK055_1508 | -1.081 | VK055_2805 | 0.788 | VK055_4101 | 0.922 |
| VK055_0196 | 0.568 | VK055_1509 | -1.456 | VK055_2806 | 0.083 | VK055_4102 | 0.044 |
| VK055_0197 | 0.506 | VK055_1510 | 0.181 | VK055_2807 | 0.548 | VK055_4103 | -1.242 |
| VK055_0198 | 0.950 | VK055_1511 | 0.838 | VK055_2808 | -0.833 | VK055_4105 | 0.260 |
| VK055_0199 | 0.218 | VK055_1512 | 1.915 | VK055_2810 | 0.039 | VK055_4106 | -0.189 |
| VK055_0200 | 0.662 | VK055_1513 | 0.202 | VK055_2811 | 1.317 | VK055_4107 | -0.022 |
| VK055_0201 | -0.136 | VK055_1515 | -0.193 | VK055_2812 | 0.655 | VK055_4108 | -0.773 |
| VK055_0202 | -0.654 | VK055_1517 | -0.666 | VK055_2813 | 0.886 | VK055_4109 | -1.112 |
| VK055_0203 | -0.895 | VK055_1519 | 0.229 | VK055_2814 | 0.150 | VK055_4110 | -0.592 |
| VK055_0204 | 0.646 | VK055_1520 | -0.536 | VK055_2815 | 0.139 | VK055_4111 | -0.030 |
| VK055_0206 | 0.299 | VK055_1521 | -1.469 | VK055_2816 | 0.778 | VK055_4112 | 0.214 |
| VK055_0208 | 0.418 | VK055_1522 | -0.044 | VK055_2817 | 0.133 | VK055_4113 | -1.485 |
| VK055_0209 | 0.538 | VK055_1523 | -0.163 | VK055_2818 | -1.165 | VK055_4115 | 0.720 |
| VK055_0210 | 1.297 | VK055_1524 | 1.260 | VK055_2820 | 1.367 | VK055_4116 | -1.002 |
| VK055_0211 | 0.216 | VK055_1525 | -0.175 | VK055_2822 | 0.638 | VK055_4117 | 0.376 |
| VK055_0221 | -0.010 | VK055_1527 | -0.187 | VK055_2823 | 1.438 | VK055_4118 | 0.207 |
| VK055_0222 | 0.872 | VK055_1528 | -0.154 | VK055_2824 | 0.495 | VK055_4119 | -0.812 |
| VK055_0223 | -0.511 | VK055_1530 | 0.031 | VK055_2825 | -1.249 | VK055_4120 | 0.454 |
| VK055_0225 | 0.950 | VK055_1531 | 0.001 | VK055_2826 | 0.411 | VK055_4121 | -1.177 |
| VK055_0226 | 0.811 | VK055_1532 | -0.053 | VK055_2827 | -0.871 | VK055_4122 | 0.126 |
| VK055_0227 | 0.429 | VK055_1537 | -0.644 | VK055_2829 | -0.708 | VK055_4123 | -0.277 |
| VK055_0228 | 0.528 | VK055_1538 | 0.384 | VK055_2830 | 0.754 | VK055_4124 | 0.696 |
| VK055_0229 | -0.275 | VK055_1541 | 0.692 | VK055_2831 | -0.886 | VK055_4126 | 0.653 |
| VK055_0230 | 0.175 | VK055_1544 | 0.975 | VK055_2832 | 1.287 | VK055_4129 | 2.050 |
| VK055_0231 | 1.476 | VK055_1548 | 0.058 | VK055_2833 | 0.636 | VK055_4130 | -1.096 |
| VK055_0232 | -0.631 | VK055_1549 | -0.523 | VK055_2834 | -1.462 | VK055_4131 | -1.427 |
| VK055_0233 | 0.261 | VK055_1550 | 0.085 | VK055_2835 | 0.557 | VK055_4135 | -0.590 |
| VK055_0234 | 2.173 | VK055_1551 | -0.234 | VK055_2836 | -0.733 | VK055_4136 | -0.043 |
| VK055_0235 | 0.621 | VK055_1552 | 0.152 | VK055_2838 | -0.272 | VK055_4138 | -0.550 |
| VK055_0236 | 0.567 | VK055_1553 | -0.027 | VK055_2839 | 0.021 | VK055_4141 | -0.740 |
| VK055_0237 | -0.061 | VK055_1555 | 0.161 | VK055_2840 | -0.538 | VK055_4143 | -0.570 |
| VK055_0238 | 1.904 | VK055_1556 | -0.836 | VK055_2841 | 0.028 | VK055_4144 | -0.854 |
| VK055_0239 | 1.497 | VK055_1557 | 0.460 | VK055_2842 | -0.319 | VK055_4145 | -1.485 |
| VK055_0241 | -0.285 | VK055_1558 | -1.329 | VK055_2843 | 0.952 | VK055_4149 | 0.148 |
| VK055_0242 | 0.150 | VK055_1559 | -0.013 | VK055_2844 | -0.606 | VK055_4150 | -0.825 |
| VK055_0243 | -1.464 | VK055_1561 | 0.282 | VK055_2845 | 0.168 | VK055_4152 | 1.278 |
| VK055_0246 | 0.775 | VK055_1562 | -0.763 | VK055_2846 | -0.334 | VK055_4153 | 0.294 |
| VK055_0247 | 1.663 | VK055_1564 | 0.439 | VK055_2847 | -0.601 | VK055_4154 | 0.703 |
| VK055_0248 | -0.214 | VK055_1565 | 0.020 | VK055_2848 | 0.081 | VK055_4155 | -0.021 |
| VK055_0249 | -1.470 | VK055_1566 | 1.019 | VK055_2849 | 0.169 | VK055_4157 | -0.014 |
| VK055_0250 | -0.238 | VK055_1567 | -0.028 | VK055_2850 | -0.762 | VK055_4159 | -0.647 |
| VK055_0253 | 1.108 | VK055_1568 | -1.471 | VK055_2851 | -0.069 | VK055_4161 | -0.762 |
| VK055_0254 | 0.181 | VK055_1569 | 0.450 | VK055_2852 | -0.289 | VK055_4162 | -1.015 |
| VK055_0258 | -0.771 | VK055_1571 | 1.260 | VK055_2853 | 0.939 | VK055_4165 | 0.322 |
| VK055_0259 | 0.342 | VK055_1573 | 0.346 | VK055_2854 | 0.172 | VK055_4166 | -0.097 |
| VK055_0260 | 0.057 | VK055_1574 | 0.193 | VK055_2855 | -1.491 | VK055_4167 | -0.392 |
| VK055_0261 | -0.424 | VK055_1575 | 0.336 | VK055_2856 | 0.348 | VK055_4168 | 0.716 |
| VK055_0262 | -0.235 | VK055_1576 | 1.028 | VK055_2857 | 0.943 | VK055_4169 | 0.502 |
| VK055_0263 | -0.373 | VK055_1577 | 0.437 | VK055_2859 | 0.346 | VK055_4170 | 0.285 |
| VK055_0265 | -1.428 | VK055_1578 | -1.477 | VK055_2860 | -1.213 | VK055_4171 | 1.008 |
| VK055_0267 | -0.772 | VK055_1579 | -0.110 | VK055_2861 | -0.072 | VK055_4172 | -1.242 |
| VK055_0269 | 0.117 | VK055_1580 | 0.187 | VK055_2862 | 0.030 | VK055_4173 | -0.706 |
| VK055_0270 | -0.191 | VK055_1581 | 0.246 | VK055_2863 | -0.157 | VK055_4174 | -0.395 |
| VK055_0271 | 0.510 | VK055_1583 | -1.476 | VK055_2865 | -1.292 | VK055_4175 | 0.269 |
| VK055_0274 | -1.152 | VK055_1584 | 0.410 | VK055_2870 | -0.196 | VK055_4176 | 0.115 |
| VK055_0284 | 0.351 | VK055_1585 | 0.097 | VK055_2871 | 0.218 | VK055_4177 | -0.169 |
| VK055_0285 | 0.104 | VK055_1586 | 0.392 | VK055_2872 | -1.497 | VK055_4180 | -0.417 |
| VK055_0286 | 1.009 | VK055_1587 | -0.855 | VK055_2873 | -0.673 | VK055_4182 | -0.057 |
| VK055_0287 | 0.863 | VK055_1588 | 0.185 | VK055_2874 | -0.379 | VK055_4183 | 0.071 |
| VK055_0288 | -1.383 | VK055_1590 | -0.052 | VK055_2876 | 0.877 | VK055_4184 | 0.220 |
| VK055_0289 | -0.792 | VK055_1591 | 0.077 | VK055_2877 | -0.760 | VK055_4188 | -0.127 |
| VK055_0292 | 0.017 | VK055_1592 | 1.471 | VK055_2878 | 0.080 | VK055_4189 | 0.834 |
| VK055_0294 | 0.848 | VK055_1593 | -0.678 | VK055_2880 | 0.047 | VK055_4190 | -0.697 |
| VK055_0295 | -0.278 | VK055_1594 | -0.625 | VK055_2881 | 0.016 | VK055_4192 | -0.749 |
| VK055_0297 | 0.923 | VK055_1595 | -0.550 | VK055_2882 | 1.460 | VK055_4193 | -0.636 |
| VK055_0300 | -0.386 | VK055_1596 | -0.819 | VK055_2883 | -0.785 | VK055_4194 | 1.738 |
| VK055_0301 | -0.255 | VK055_1597 | -1.134 | VK055_2884 | -0.788 | VK055_4196 | 0.261 |
| VK055_0304 | -0.414 | VK055_1598 | -0.169 | VK055_2885 | 0.751 | VK055_4200 | 0.002 |
| VK055_0305 | 0.717 | VK055_1599 | 0.467 | VK055_2886 | 0.332 | VK055_4201 | -0.650 |
| VK055_0306 | -0.241 | VK055_1602 | 0.769 | VK055_2887 | -0.665 | VK055_4202 | -0.769 |
| VK055_0307 | 0.501 | VK055_1603 | -0.223 | VK055_2888 | 0.017 | VK055_4203 | -1.238 |
| VK055_0308 | 0.823 | VK055_1604 | 0.266 | VK055_2889 | 0.427 | VK055_4204 | 0.620 |
| VK055_0309 | -0.623 | VK055_1605 | 0.810 | VK055_2890 | 0.618 | VK055_4205 | 0.557 |
| VK055_0310 | 0.577 | VK055_1606 | -0.435 | VK055_2891 | -0.658 | VK055_4207 | -0.273 |
| VK055_0311 | -0.023 | VK055_1607 | 1.179 | VK055_2892 | 0.177 | VK055_4208 | 0.026 |
| VK055_0312 | 1.104 | VK055_1608 | -0.242 | VK055_2893 | -0.823 | VK055_4209 | 1.428 |
| VK055_0314 | 0.157 | VK055_1613 | -0.898 | VK055_2894 | 0.819 | VK055_4210 | -0.592 |
| VK055_0315 | 0.345 | VK055_1615 | 0.189 | VK055_2895 | 0.380 | VK055_4211 | -0.073 |
| VK055_0320 | 0.105 | VK055_1617 | -0.319 | VK055_2896 | 2.293 | VK055_4212 | -0.330 |
| VK055_0321 | -0.762 | VK055_1618 | 0.253 | VK055_2897 | 2.231 | VK055_4213 | 0.109 |
| VK055_0322 | -0.389 | VK055_1619 | 0.586 | VK055_2899 | 0.088 | VK055_4214 | -0.808 |
| VK055_0323 | -0.872 | VK055_1620 | 0.739 | VK055_2900 | 0.966 | VK055_4215 | 0.340 |
| VK055_0324 | 0.434 | VK055_1625 | 0.057 | VK055_2904 | 0.348 | VK055_4216 | -0.884 |
| VK055_0325 | -0.906 | VK055_1626 | 0.187 | VK055_2909 | 0.301 | VK055_4217 | -0.166 |
| VK055_0327 | 0.507 | VK055_1627 | -0.719 | VK055_2910 | 0.602 | VK055_4218 | 0.941 |
| VK055_0330 | -0.991 | VK055_1628 | -1.108 | VK055_2912 | 0.367 | VK055_4219 | -0.135 |
| VK055_0331 | -0.549 | VK055_1630 | -0.418 | VK055_2913 | 0.154 | VK055_4220 | -0.078 |
| VK055_0332 | 0.411 | VK055_1631 | 1.138 | VK055_2914 | 0.054 | VK055_4221 | -0.532 |
| VK055_0333 | 0.500 | VK055_1632 | 2.324 | VK055_2915 | -1.203 | VK055_4222 | 0.170 |
| VK055_0335 | 2.107 | VK055_1633 | 0.098 | VK055_2916 | 1.903 | VK055_4224 | -1.309 |
| VK055_0338 | 0.929 | VK055_1635 | 0.222 | VK055_2917 | -1.628 | VK055_4226 | 0.325 |
| VK055_0339 | -0.068 | VK055_1636 | -0.046 | VK055_2918 | -0.825 | VK055_4227 | -1.373 |
| VK055_0340 | -0.751 | VK055_1638 | -1.035 | VK055_2921 | -0.568 | VK055_4229 | -1.497 |
| VK055_0342 | -0.652 | VK055_1639 | 0.156 | VK055_2922 | 0.184 | VK055_4230 | 0.651 |
| VK055_0343 | -0.268 | VK055_1640 | -0.651 | VK055_2923 | -0.169 | VK055_4231 | 1.203 |
| VK055_0344 | 0.529 | VK055_1641 | -1.403 | VK055_2924 | -0.015 | VK055_4235 | -1.483 |
| VK055_0345 | 0.365 | VK055_1643 | 0.109 | VK055_2925 | -0.143 | VK055_4236 | 0.487 |
| VK055_0346 | -1.497 | VK055_1647 | 0.322 | VK055_2926 | -0.621 | VK055_4237 | 0.777 |
| VK055_0347 | -0.322 | VK055_1648 | 1.018 | VK055_2927 | -1.422 | VK055_4238 | -0.001 |
| VK055_0348 | -0.482 | VK055_1649 | 1.403 | VK055_2928 | 1.938 | VK055_4239 | 0.148 |
| VK055_0349 | -0.139 | VK055_1650 | 1.495 | VK055_2929 | -1.059 | VK055_4240 | 0.005 |
| VK055_0350 | -0.010 | VK055_1652 | 0.956 | VK055_2933 | -0.232 | VK055_4242 | -0.515 |
| VK055_0351 | 0.458 | VK055_1653 | -0.772 | VK055_2935 | 1.158 | VK055_4243 | 0.104 |
| VK055_0353 | 0.047 | VK055_1654 | -1.073 | VK055_2936 | -1.440 | VK055_4244 | 0.656 |
| VK055_0354 | 0.322 | VK055_1655 | -0.423 | VK055_2937 | -0.156 | VK055_4245 | 0.200 |
| VK055_0354 | 0.037 | VK055_1657 | -0.601 | VK055_2938 | -0.187 | VK055_4246 | -0.681 |
| VK055_0355 | -0.642 | VK055_1658 | -0.342 | VK055_2939 | -0.086 | VK055_4247 | -0.393 |
| VK055_0356 | 0.568 | VK055_1659 | 0.333 | VK055_2940 | 0.416 | VK055_4249 | -0.125 |
| VK055_0357 | -0.645 | VK055_1660 | -0.120 | VK055_2942 | 0.936 | VK055_4250 | 1.002 |
| VK055_0358 | 0.731 | VK055_1662 | -0.129 | VK055_2943 | -0.581 | VK055_4251 | -0.125 |
| VK055_0360 | 0.557 | VK055_1663 | -0.807 | VK055_2944 | -1.267 | VK055_4252 | -0.735 |
| VK055_0361 | -1.034 | VK055_1664 | -0.196 | VK055_2946 | 0.330 | VK055_4253 | 1.774 |
| VK055_0363 | -1.490 | VK055_1665 | 0.356 | VK055_2948 | 0.217 | VK055_4254 | -1.223 |
| VK055_0364 | -0.485 | VK055_1666 | -0.033 | VK055_2949 | 0.057 | VK055_4255 | -0.617 |
| VK055_0366 | -0.859 | VK055_1667 | 0.483 | VK055_2950 | -0.090 | VK055_4256 | -0.776 |
| VK055_0367 | 0.419 | VK055_1669 | -1.472 | VK055_2952 | -1.359 | VK055_4258 | 0.182 |
| VK055_0368 | -0.043 | VK055_1670 | -0.246 | VK055_2954 | -0.461 | VK055_4259 | 0.483 |
| VK055_0369 | -0.538 | VK055_1671 | -0.196 | VK055_2955 | 1.187 | VK055_4260 | -0.618 |
| VK055_0370 | -0.466 | VK055_1672 | 0.338 | VK055_2957 | -0.161 | VK055_4261 | -0.102 |
| VK055_0371 | 0.068 | VK055_1673 | 0.217 | VK055_2958 | 0.067 | VK055_4262 | 0.147 |
| VK055_0372 | 0.262 | VK055_1674 | 0.294 | VK055_2960 | -0.079 | VK055_4263 | 0.158 |
| VK055_0375 | -0.947 | VK055_1675 | 0.708 | VK055_2961 | -0.842 | VK055_4264 | 0.277 |
| VK055_0376 | 0.795 | VK055_1676 | 0.445 | VK055_2962 | -0.103 | VK055_4265 | 0.005 |
| VK055_0377 | -0.139 | VK055_1677 | -0.010 | VK055_2963 | -0.840 | VK055_4266 | -0.292 |
| VK055_0379 | 0.561 | VK055_1678 | 0.519 | VK055_2965 | -0.154 | VK055_4267 | 2.325 |
| VK055_0380 | -0.265 | VK055_1679 | -0.097 | VK055_2966 | -0.058 | VK055_4268 | 0.456 |
| VK055_0382 | 0.227 | VK055_1680 | -1.490 | VK055_2967 | -0.002 | VK055_4269 | 0.017 |
| VK055_0384 | 0.240 | VK055_1681 | -0.275 | VK055_2969 | 0.760 | VK055_4270 | 1.439 |
| VK055_0385 | 1.390 | VK055_1682 | 0.013 | VK055_2970 | 0.208 | VK055_4271 | 0.167 |
| VK055_0386 | -0.415 | VK055_1684 | 0.242 | VK055_2973 | -1.020 | VK055_4272 | 0.735 |
| VK055_0387 | -0.334 | VK055_1685 | 0.172 | VK055_2974 | 0.942 | VK055_4273 | -0.362 |
| VK055_0388 | 1.344 | VK055_1687 | 0.303 | VK055_2975 | -0.467 | VK055_4274 | 0.194 |
| VK055_0389 | -0.810 | VK055_1688 | 0.770 | VK055_2976 | 0.131 | VK055_4275 | 0.245 |
| VK055_0390 | 0.414 | VK055_1689 | -0.004 | VK055_2977 | 0.877 | VK055_4276 | 1.134 |
| VK055_0391 | -0.078 | VK055_1690 | -0.597 | VK055_2978 | 0.087 | VK055_4277 | 0.258 |
| VK055_0392 | 0.170 | VK055_1692 | 0.436 | VK055_2979 | 0.276 | VK055_4278 | -0.170 |
| VK055_0397 | 0.552 | VK055_1693 | -0.031 | VK055_2980 | 0.270 | VK055_4279 | 0.457 |
| VK055_0398 | -1.440 | VK055_1694 | -0.659 | VK055_2981 | -0.183 | VK055_4280 | 0.159 |
| VK055_0400 | -0.394 | VK055_1695 | -0.058 | VK055_2982 | -0.020 | VK055_4281 | -0.670 |
| VK055_0401 | -1.470 | VK055_1696 | -0.605 | VK055_2983 | -1.013 | VK055_4282 | 0.939 |
| VK055_0402 | 0.940 | VK055_1697 | 0.329 | VK055_2984 | 0.324 | VK055_4283 | -0.580 |
| VK055_0403 | -1.457 | VK055_1699 | 0.434 | VK055_2985 | -0.718 | VK055_4284 | 0.097 |
| VK055_0404 | 1.166 | VK055_1700 | -0.016 | VK055_2986 | -0.478 | VK055_4286 | 0.082 |
| VK055_0405 | -0.542 | VK055_1701 | -0.075 | VK055_2987 | 0.259 | VK055_4287 | -0.005 |
| VK055_0406 | 0.073 | VK055_1702 | -0.774 | VK055_2988 | 0.041 | VK055_4294 | -1.434 |
| VK055_0407 | -0.008 | VK055_1703 | -0.598 | VK055_2989 | -0.898 | VK055_4295 | 0.170 |
| VK055_0408 | -0.060 | VK055_1706 | -1.409 | VK055_2990 | 0.207 | VK055_4296 | -1.485 |
| VK055_0409 | 0.000 | VK055_1709 | -0.112 | VK055_2992 | 0.888 | VK055_4297 | -0.160 |
| VK055_0411 | -0.124 | VK055_1710 | 0.161 | VK055_2993 | 0.215 | VK055_4298 | -1.000 |
| VK055_0412 | 0.264 | VK055_1711 | 1.242 | VK055_2994 | -0.386 | VK055_4299 | 0.880 |
| VK055_0415 | 0.511 | VK055_1712 | -0.175 | VK055_2995 | 0.020 | VK055_4300 | 0.713 |
| VK055_0418 | 1.132 | VK055_1713 | -0.664 | VK055_2996 | -0.516 | VK055_4301 | 0.868 |
| VK055_0420 | -0.164 | VK055_1714 | -0.623 | VK055_2997 | 0.190 | VK055_4303 | -0.653 |
| VK055_0421 | 1.828 | VK055_1715 | 0.566 | VK055_2999 | -0.720 | VK055_4305 | 0.996 |
| VK055_0423 | -0.255 | VK055_1716 | -0.174 | VK055_3000 | 1.339 | VK055_4306 | -0.091 |
| VK055_0424 | 0.049 | VK055_1717 | -0.658 | VK055_3001 | -1.491 | VK055_4307 | 0.331 |
| VK055_0427 | 0.101 | VK055_1719 | -0.697 | VK055_3003 | -0.440 | VK055_4308 | 2.387 |
| VK055_0428 | -0.480 | VK055_1721 | 0.659 | VK055_3004 | 0.323 | VK055_4310 | -0.600 |
| VK055_0429 | 0.736 | VK055_1722 | 0.763 | VK055_3006 | 0.962 | VK055_4312 | -0.181 |
| VK055_0431 | 0.052 | VK055_1723 | 0.610 | VK055_3007 | 0.336 | VK055_4313 | 0.710 |
| VK055_0432 | -0.468 | VK055_1723 | -0.277 | VK055_3010 | -0.276 | VK055_4316 | -0.251 |
| VK055_0433 | -0.933 | VK055_1724 | 0.416 | VK055_3011 | 0.420 | VK055_4317 | -1.440 |
| VK055_0434 | 1.303 | VK055_1725 | -0.759 | VK055_3012 | 0.229 | VK055_4318 | 0.304 |
| VK055_0435 | 0.672 | VK055_1726 | -1.485 | VK055_3014 | 1.785 | VK055_4319 | 0.393 |
| VK055_0436 | 0.394 | VK055_1727 | 0.057 | VK055_3015 | 1.287 | VK055_4320 | 0.434 |
| VK055_0437 | 0.983 | VK055_1728 | 0.042 | VK055_3016 | -0.692 | VK055_4321 | 0.454 |
| VK055_0438 | 0.002 | VK055_1729 | -0.631 | VK055_3017 | -0.475 | VK055_4324 | -0.822 |
| VK055_0439 | -0.683 | VK055_1730 | 1.122 | VK055_3020 | 0.698 | VK055_4325 | 1.446 |
| VK055_0440 | 0.670 | VK055_1731 | -0.269 | VK055_3022 | -0.961 | VK055_4326 | -1.249 |
| VK055_0441 | 0.903 | VK055_1732 | 1.679 | VK055_3024 | 0.582 | VK055_4327 | -0.404 |
| VK055_0442 | 0.921 | VK055_1733 | -0.855 | VK055_3026 | -0.535 | VK055_4328 | 0.008 |
| VK055_0443 | 1.120 | VK055_1734 | 0.811 | VK055_3027 | 0.788 | VK055_4332 | 0.089 |
| VK055_0444 | 1.730 | VK055_1735 | 0.295 | VK055_3029 | 0.237 | VK055_4334 | -0.189 |
| VK055_0446 | 0.790 | VK055_1736 | 0.397 | VK055_3030 | -0.026 | VK055_4335 | 0.771 |
| VK055_0447 | 0.771 | VK055_1737 | -0.075 | VK055_3031 | 0.496 | VK055_4336 | 0.103 |
| VK055_0448 | -0.522 | VK055_1738 | -0.396 | VK055_3032 | -0.867 | VK055_4337 | -0.863 |
| VK055_0450 | 0.337 | VK055_1739 | 0.052 | VK055_3033 | 1.292 | VK055_4338 | -1.196 |
| VK055_0451 | 0.739 | VK055_1740 | -0.879 | VK055_3035 | 0.649 | VK055_4339 | -0.739 |
| VK055_0452 | -1.299 | VK055_1741 | 0.579 | VK055_3036 | 0.506 | VK055_4342 | -0.005 |
| VK055_0454 | 2.399 | VK055_1742 | -0.254 | VK055_3037 | 0.402 | VK055_4343 | -0.871 |
| VK055_0455 | 0.320 | VK055_1744 | -0.743 | VK055_3038 | 0.028 | VK055_4344 | 0.466 |
| VK055_0456 | 0.023 | VK055_1745 | -0.017 | VK055_3039 | 1.431 | VK055_4347 | -1.141 |
| VK055_0457 | 0.440 | VK055_1746 | 1.249 | VK055_3041 | 1.501 | VK055_4348 | 0.235 |
| VK055_0458 | -0.202 | VK055_1749 | 0.318 | VK055_3043 | 0.088 | VK055_4350 | -0.369 |
| VK055_0459 | 0.067 | VK055_1750 | -1.075 | VK055_3046 | 0.171 | VK055_4351 | 0.046 |
| VK055_0460 | -1.057 | VK055_1751 | -0.913 | VK055_3047 | 0.119 | VK055_4353 | 0.399 |
| VK055_0462 | 0.201 | VK055_1752 | 1.261 | VK055_3048 | -0.216 | VK055_4354 | 0.506 |
| VK055_0463 | -0.201 | VK055_1754 | -1.440 | VK055_3049 | -1.470 | VK055_4355 | 0.523 |
| VK055_0466 | -1.455 | VK055_1756 | 1.096 | VK055_3051 | 0.631 | VK055_4356 | -0.755 |
| VK055_0474 | -1.171 | VK055_1757 | -0.372 | VK055_3052 | 0.842 | VK055_4357 | 0.179 |
| VK055_0475 | -0.302 | VK055_1758 | -0.281 | VK055_3053 | -0.599 | VK055_4358 | -0.149 |
| VK055_0476 | -1.478 | VK055_1759 | -0.159 | VK055_3054 | -0.556 | VK055_4359 | -0.198 |
| VK055_0478 | 0.789 | VK055_1761 | -0.184 | VK055_3055 | -0.031 | VK055_4360 | 0.348 |
| VK055_0479 | 0.688 | VK055_1762 | -0.358 | VK055_3056 | 0.898 | VK055_4361 | 0.487 |
| VK055_0480 | -1.015 | VK055_1763 | 0.369 | VK055_3057 | 0.694 | VK055_4362 | -0.612 |
| VK055_0481 | 0.162 | VK055_1764 | 0.413 | VK055_3059 | 0.386 | VK055_4363 | 1.165 |
| VK055_0482 | 1.374 | VK055_1765 | 0.720 | VK055_3060 | -0.008 | VK055_4365 | 0.267 |
| VK055_0483 | 0.643 | VK055_1766 | -0.059 | VK055_3061 | -0.833 | VK055_4367 | 1.304 |
| VK055_0485 | 0.429 | VK055_1768 | -0.705 | VK055_3062 | 0.815 | VK055_4368 | 0.134 |
| VK055_0488 | -0.072 | VK055_1769 | 0.537 | VK055_3064 | 0.768 | VK055_4369 | -0.777 |
| VK055_0489 | 1.289 | VK055_1770 | -0.051 | VK055_3065 | -0.853 | VK055_4370 | -0.900 |
| VK055_0490 | -0.188 | VK055_1771 | 1.108 | VK055_3066 | -0.107 | VK055_4371 | 0.113 |
| VK055_0492 | 0.494 | VK055_1772 | 0.165 | VK055_3067 | -1.316 | VK055_4372 | 0.725 |
| VK055_0493 | -0.825 | VK055_1780 | -1.008 | VK055_3069 | 0.013 | VK055_4373 | 0.276 |
| VK055_0494 | 0.836 | VK055_1789 | -1.497 | VK055_3070 | 0.079 | VK055_4374 | -0.282 |
| VK055_0495 | 0.459 | VK055_1790 | 0.528 | VK055_3072 | -0.603 | VK055_4375 | 0.121 |
| VK055_0496 | -0.031 | VK055_1791 | -0.487 | VK055_3073 | 0.528 | VK055_4376 | 1.084 |
| VK055_0497 | -0.909 | VK055_1793 | -0.883 | VK055_3074 | -0.090 | VK055_4379 | 0.194 |
| VK055_0500 | 0.565 | VK055_1794 | 0.249 | VK055_3075 | -0.602 | VK055_4380 | 1.360 |
| VK055_0503 | 0.151 | VK055_1797 | -0.086 | VK055_3076 | 0.868 | VK055_4381 | 0.395 |
| VK055_0504 | -0.307 | VK055_1798 | -1.182 | VK055_3077 | -0.382 | VK055_4382 | -1.340 |
| VK055_0505 | 0.555 | VK055_1799 | -0.309 | VK055_3078 | 0.308 | VK055_4383 | 0.026 |
| VK055_0506 | 0.554 | VK055_1800 | -0.847 | VK055_3079 | 1.084 | VK055_4384 | 0.446 |
| VK055_0507 | 0.149 | VK055_1802 | -0.349 | VK055_3080 | 1.510 | VK055_4386 | -0.916 |
| VK055_0508 | 0.157 | VK055_1803 | 0.274 | VK055_3081 | -1.483 | VK055_4387 | -0.673 |
| VK055_0509 | 0.122 | VK055_1804 | 0.436 | VK055_3086 | -0.015 | VK055_4388 | 0.912 |
| VK055_0510 | 0.747 | VK055_1805 | 1.391 | VK055_3087 | -1.378 | VK055_4389 | -1.226 |
| VK055_0511 | -0.225 | VK055_1806 | 0.043 | VK055_3088 | -0.824 | VK055_4392 | 0.244 |
| VK055_0512 | 0.046 | VK055_1807 | 0.363 | VK055_3089 | 0.455 | VK055_4394 | -0.097 |
| VK055_0515 | -0.874 | VK055_1808 | 0.658 | VK055_3090 | -0.865 | VK055_4395 | 0.630 |
| VK055_0516 | 0.645 | VK055_1809 | 0.194 | VK055_3091 | 2.445 | VK055_4397 | 0.788 |
| VK055_0517 | -1.497 | VK055_1810 | -1.110 | VK055_3092 | -0.837 | VK055_4398 | -0.274 |
| VK055_0518 | -0.596 | VK055_1811 | -0.896 | VK055_3093 | 1.610 | VK055_4400 | 0.578 |
| VK055_0519 | 0.172 | VK055_1812 | -0.172 | VK055_3094 | -0.468 | VK055_4401 | 0.835 |
| VK055_0521 | -0.583 | VK055_1813 | 0.306 | VK055_3096 | 0.534 | VK055_4402 | -1.320 |
| VK055_0523 | 0.712 | VK055_1814 | -1.490 | VK055_3097 | 1.654 | VK055_4403 | 1.562 |
| VK055_0524 | 0.722 | VK055_1815 | 0.178 | VK055_3098 | -0.031 | VK055_4404 | 0.172 |
| VK055_0526 | 0.037 | VK055_1816 | 0.701 | VK055_3099 | 0.763 | VK055_4405 | 0.011 |
| VK055_0527 | -0.174 | VK055_1817 | 0.619 | VK055_3100 | -0.390 | VK055_4409 | -1.399 |
| VK055_0528 | -0.121 | VK055_1818 | -0.121 | VK055_3103 | 0.091 | VK055_4410 | 0.868 |
| VK055_0529 | 0.065 | VK055_1819 | -0.486 | VK055_3104 | 0.274 | VK055_4411 | -0.383 |
| VK055_0530 | 0.709 | VK055_1820 | 1.661 | VK055_3105 | 0.513 | VK055_4412 | -0.536 |
| VK055_0531 | 1.030 | VK055_1825 | 0.664 | VK055_3106 | -0.750 | VK055_4413 | 0.701 |
| VK055_0532 | -0.817 | VK055_1826 | -0.113 | VK055_3108 | -0.092 | VK055_4414 | -0.159 |
| VK055_0533 | -0.119 | VK055_1827 | -1.483 | VK055_3131 | 0.108 | VK055_4415 | 1.058 |
| VK055_0534 | 0.008 | VK055_1828 | 1.794 | VK055_3132 | -0.511 | VK055_4416 | -0.338 |
| VK055_0535 | 1.326 | VK055_1829 | -0.150 | VK055_3134 | 0.043 | VK055_4419 | -0.470 |
| VK055_0536 | 0.290 | VK055_1831 | -0.049 | VK055_3135 | 0.828 | VK055_4421 | -0.092 |
| VK055_0537 | -0.090 | VK055_1832 | -0.172 | VK055_3136 | -0.566 | VK055_4422 | -0.214 |
| VK055_0540 | -0.435 | VK055_1833 | -0.879 | VK055_3137 | -0.768 | VK055_4423 | 0.177 |
| VK055_0542 | -0.542 | VK055_1834 | -0.154 | VK055_3139 | 0.764 | VK055_4424 | -0.719 |
| VK055_0543 | -0.285 | VK055_1835 | -0.723 | VK055_3141 | 0.979 | VK055_4425 | -0.867 |
| VK055_0546 | -1.469 | VK055_1836 | 1.657 | VK055_3142 | -0.401 | VK055_4426 | -1.114 |
| VK055_0548 | -0.679 | VK055_1845 | -0.133 | VK055_3143 | 0.761 | VK055_4427 | 0.566 |
| VK055_0550 | -0.224 | VK055_1848 | -0.628 | VK055_3148 | -1.476 | VK055_4429 | -0.254 |
| VK055_0551 | 0.093 | VK055_1850 | -0.212 | VK055_3149 | -0.410 | VK055_4430 | -0.055 |
| VK055_0552 | 1.996 | VK055_1851 | -0.483 | VK055_3150 | -0.068 | VK055_4433 | -0.239 |
| VK055_0553 | -0.024 | VK055_1852 | -0.399 | VK055_3151 | 0.026 | VK055_4434 | 2.247 |
| VK055_0554 | 0.237 | VK055_1853 | 0.606 | VK055_3152 | -1.180 | VK055_4436 | -0.038 |
| VK055_0555 | -0.292 | VK055_1855 | -0.415 | VK055_3153 | 0.451 | VK055_4437 | -1.004 |
| VK055_0556 | 0.235 | VK055_1856 | -0.762 | VK055_3154 | 0.369 | VK055_4438 | 0.329 |
| VK055_0557 | 0.741 | VK055_1857 | -0.419 | VK055_3156 | 1.351 | VK055_4439 | -0.419 |
| VK055_0558 | -0.113 | VK055_1862 | 0.770 | VK055_3157 | 0.817 | VK055_4440 | 0.485 |
| VK055_0561 | -0.065 | VK055_1868 | -0.810 | VK055_3158 | -0.087 | VK055_4441 | 0.785 |
| VK055_0563 | -0.040 | VK055_1869 | 0.281 | VK055_3159 | 0.358 | VK055_4442 | -0.614 |
| VK055_0564 | 0.863 | VK055_1870 | -1.440 | VK055_3160 | -1.484 | VK055_4443 | -1.219 |
| VK055_0566 | -0.861 | VK055_1871 | -0.333 | VK055_3161 | -0.354 | VK055_4444 | 1.498 |
| VK055_0567 | 0.655 | VK055_1872 | -0.182 | VK055_3162 | -0.277 | VK055_4445 | 0.256 |
| VK055_0570 | -0.652 | VK055_1873 | 0.636 | VK055_3163 | 1.148 | VK055_4446 | -0.244 |
| VK055_0572 | -1.228 | VK055_1874 | 0.542 | VK055_3164 | 0.965 | VK055_4448 | -0.499 |
| VK055_0573 | -1.153 | VK055_1876 | 0.526 | VK055_3166 | -0.949 | VK055_4452 | 0.559 |
| VK055_0574 | 0.337 | VK055_1877 | 0.085 | VK055_3167 | -1.224 | VK055_4454 | -0.651 |
| VK055_0576 | -0.950 | VK055_1878 | -0.419 | VK055_3168 | -0.871 | VK055_4455 | -0.786 |
| VK055_0578 | -0.146 | VK055_1879 | 0.264 | VK055_3170 | -1.424 | VK055_4456 | 0.036 |
| VK055_0581 | 0.855 | VK055_1880 | 0.974 | VK055_3173 | -1.091 | VK055_4457 | -0.779 |
| VK055_0582 | 0.030 | VK055_1881 | -1.146 | VK055_3174 | -1.497 | VK055_4458 | -0.960 |
| VK055_0583 | -0.494 | VK055_1882 | 0.660 | VK055_3179 | 0.338 | VK055_4459 | -0.762 |
| VK055_0584 | -0.024 | VK055_1884 | 1.398 | VK055_3180 | -1.216 | VK055_4460 | 1.874 |
| VK055_0585 | -0.207 | VK055_1885 | 0.618 | VK055_3181 | 1.068 | VK055_4462 | 1.551 |
| VK055_0586 | 0.447 | VK055_1886 | -0.305 | VK055_3182 | -1.085 | VK055_4463 | 0.900 |
| VK055_0587 | -1.141 | VK055_1887 | -1.096 | VK055_3183 | -0.982 | VK055_4464 | -0.252 |
| VK055_0588 | -0.018 | VK055_1888 | -0.517 | VK055_3184 | -0.899 | VK055_4465 | -0.829 |
| VK055_0591 | 0.396 | VK055_1889 | -1.060 | VK055_3185 | -0.637 | VK055_4467 | 0.412 |
| VK055_0592 | -1.434 | VK055_1891 | -0.480 | VK055_3186 | -1.171 | VK055_4468 | 0.052 |
| VK055_0593 | -0.623 | VK055_1892 | -0.051 | VK055_3187 | -0.538 | VK055_4469 | 0.279 |
| VK055_0594 | -0.145 | VK055_1893 | -0.826 | VK055_3188 | 0.285 | VK055_4470 | 0.367 |
| VK055_0595 | -1.478 | VK055_1894 | -0.610 | VK055_3189 | -0.008 | VK055_4473 | 0.208 |
| VK055_0596 | -0.679 | VK055_1895 | -1.067 | VK055_3190 | 0.518 | VK055_4474 | 0.722 |
| VK055_0597 | -1.371 | VK055_1896 | 0.528 | VK055_3191 | -1.076 | VK055_4475 | 0.385 |
| VK055_0598 | 0.023 | VK055_1898 | 0.157 | VK055_3194 | -0.576 | VK055_4478 | 0.348 |
| VK055_0599 | 0.245 | VK055_1899 | 0.274 | VK055_3195 | -0.211 | VK055_4479 | -0.843 |
| VK055_0600 | -0.010 | VK055_1900 | 0.626 | VK055_3196 | -1.230 | VK055_4480 | -0.493 |
| VK055_0602 | 0.029 | VK055_1902 | 1.137 | VK055_3197 | -0.626 | VK055_4482 | 0.267 |
| VK055_0603 | -0.046 | VK055_1904 | 0.635 | VK055_3198 | -0.020 | VK055_4483 | -1.264 |
| VK055_0604 | -0.468 | VK055_1905 | 0.855 | VK055_3200 | -0.139 | VK055_4485 | -0.734 |
| VK055_0605 | 0.126 | VK055_1906 | -0.448 | VK055_3202 | -0.197 | VK055_4486 | 0.547 |
| VK055_0606 | 1.101 | VK055_1907 | 0.334 | VK055_3203 | 0.628 | VK055_4495 | 2.425 |
| VK055_0607 | 0.052 | VK055_1908 | -0.072 | VK055_3204 | 0.057 | VK055_4496 | 1.086 |
| VK055_0608 | 0.552 | VK055_1911 | -0.583 | VK055_3205 | 0.003 | VK055_4497 | 0.402 |
| VK055_0610 | 0.016 | VK055_1912 | -0.122 | VK055_3206 | 1.892 | VK055_4498 | -1.244 |
| VK055_0611 | -0.762 | VK055_1913 | -0.434 | VK055_3207 | -0.652 | VK055_4500 | -0.811 |
| VK055_0612 | -0.080 | VK055_1914 | 0.038 | VK055_3208 | 0.411 | VK055_4501 | -0.536 |
| VK055_0613 | 0.332 | VK055_1915 | 0.917 | VK055_3210 | 0.428 | VK055_4502 | 0.459 |
| VK055_0614 | 1.657 | VK055_1916 | 0.742 | VK055_3211 | -0.106 | VK055_4503 | 0.095 |
| VK055_0615 | -1.029 | VK055_1917 | 0.728 | VK055_3212 | 0.027 | VK055_4504 | 0.435 |
| VK055_0616 | -0.180 | VK055_1918 | -0.057 | VK055_3220 | 0.740 | VK055_4505 | -0.621 |
| VK055_0617 | -0.833 | VK055_1919 | -0.375 | VK055_3221 | -1.283 | VK055_4506 | -1.483 |
| VK055_0619 | 0.141 | VK055_1920 | -0.985 | VK055_3222 | 0.297 | VK055_4507 | -0.253 |
| VK055_0622 | 0.715 | VK055_1921 | -0.271 | VK055_3223 | -0.976 | VK055_4508 | -0.042 |
| VK055_0624 | -0.179 | VK055_1922 | 0.284 | VK055_3224 | 0.886 | VK055_4510 | -1.469 |
| VK055_0625 | 0.285 | VK055_1923 | 2.560 | VK055_3225 | -1.440 | VK055_4511 | 0.843 |
| VK055_0626 | 0.566 | VK055_1924 | -0.801 | VK055_3226 | -0.344 | VK055_4512 | -0.347 |
| VK055_0627 | -0.486 | VK055_1925 | 0.251 | VK055_3227 | -0.591 | VK055_4514 | -0.210 |
| VK055_0628 | 0.173 | VK055_1926 | -0.766 | VK055_3228 | -0.678 | VK055_4515 | -0.596 |
| VK055_0629 | -0.125 | VK055_1927 | -1.440 | VK055_3229 | 0.817 | VK055_4516 | 1.174 |
| VK055_0630 | -0.191 | VK055_1928 | 1.242 | VK055_3230 | 0.763 | VK055_4517 | 0.147 |
| VK055_0631 | 0.859 | VK055_1931 | -0.716 | VK055_3231 | 0.240 | VK055_4518 | -0.306 |
| VK055_0632 | 0.762 | VK055_1932 | -1.044 | VK055_3232 | 1.303 | VK055_4519 | -0.231 |
| VK055_0635 | 0.454 | VK055_1933 | 0.762 | VK055_3233 | -0.097 | VK055_4523 | -0.166 |
| VK055_0636 | -0.905 | VK055_1934 | 0.044 | VK055_3234 | -1.067 | VK055_4525 | -1.087 |
| VK055_0639 | 0.548 | VK055_1935 | 0.491 | VK055_3235 | 0.818 | VK055_4526 | -1.434 |
| VK055_0640 | 0.810 | VK055_1936 | -0.461 | VK055_3236 | -0.088 | VK055_4527 | 0.408 |
| VK055_0642 | 1.622 | VK055_1937 | -1.461 | VK055_3237 | -0.049 | VK055_4528 | 0.496 |
| VK055_0645 | -0.310 | VK055_1938 | 0.414 | VK055_3238 | -0.028 | VK055_4529 | 1.369 |
| VK055_0648 | 0.280 | VK055_1939 | -0.218 | VK055_3239 | 0.514 | VK055_4530 | 0.281 |
| VK055_0651 | -0.715 | VK055_1940 | -0.798 | VK055_3240 | -0.263 | VK055_4531 | 0.058 |
| VK055_0653 | 0.760 | VK055_1941 | -0.371 | VK055_3241 | -0.204 | VK055_4532 | 0.666 |
| VK055_0655 | -0.522 | VK055_1942 | 1.242 | VK055_3242 | 0.377 | VK055_4533 | -0.359 |
| VK055_0656 | 0.395 | VK055_1943 | -0.265 | VK055_3243 | -0.809 | VK055_4534 | -0.580 |
| VK055_0657 | -0.659 | VK055_1944 | 0.798 | VK055_3244 | -1.220 | VK055_4535 | -0.382 |
| VK055_0658 | -0.065 | VK055_1945 | -0.496 | VK055_3245 | -0.191 | VK055_4537 | 1.156 |
| VK055_0660 | -0.455 | VK055_1946 | 0.427 | VK055_3248 | -0.703 | VK055_4539 | -1.025 |
| VK055_0661 | 0.390 | VK055_1947 | -0.988 | VK055_3249 | 0.203 | VK055_4540 | -0.107 |
| VK055_0663 | -0.777 | VK055_1948 | 0.129 | VK055_3250 | -0.314 | VK055_4542 | 0.276 |
| VK055_0664 | -1.455 | VK055_1950 | -0.143 | VK055_3251 | -0.467 | VK055_4543 | 1.020 |
| VK055_0665 | 0.701 | VK055_1952 | -0.039 | VK055_3252 | -1.491 | VK055_4544 | 0.117 |
| VK055_0666 | 0.333 | VK055_1953 | -0.051 | VK055_3253 | 1.984 | VK055_4545 | -0.266 |
| VK055_0667 | 0.505 | VK055_1954 | 1.772 | VK055_3254 | -0.141 | VK055_4546 | 0.326 |
| VK055_0668 | 1.494 | VK055_1955 | 0.313 | VK055_3255 | 1.096 | VK055_4548 | 0.213 |
| VK055_0670 | -0.582 | VK055_1956 | -0.291 | VK055_3256 | -0.214 | VK055_4549 | -0.033 |
| VK055_0671 | -0.823 | VK055_1957 | -0.682 | VK055_3257 | -1.477 | VK055_4550 | 0.746 |
| VK055_0672 | -1.458 | VK055_1958 | -0.666 | VK055_3258 | -0.008 | VK055_4551 | -0.683 |
| VK055_0673 | -1.433 | VK055_1960 | -0.399 | VK055_3259 | -0.398 | VK055_4552 | -1.433 |
| VK055_0674 | -0.277 | VK055_1961 | 0.955 | VK055_3260 | 0.276 | VK055_4554 | -1.428 |
| VK055_0676 | 0.042 | VK055_1962 | -0.027 | VK055_3261 | 0.303 | VK055_4555 | -1.267 |
| VK055_0678 | 0.390 | VK055_1964 | 1.103 | VK055_3262 | 0.858 | VK055_4557 | 0.567 |
| VK055_0682 | 0.235 | VK055_1965 | 0.113 | VK055_3263 | -1.453 | VK055_4558 | -0.168 |
| VK055_0683 | 0.381 | VK055_1966 | -0.126 | VK055_3264 | 0.806 | VK055_4560 | -0.098 |
| VK055_0686 | 0.188 | VK055_1967 | -0.513 | VK055_3265 | -0.633 | VK055_4561 | -0.311 |
| VK055_0687 | -0.943 | VK055_1968 | 0.873 | VK055_3266 | 0.625 | VK055_4562 | -1.440 |
| VK055_0688 | 1.290 | VK055_1969 | 0.420 | VK055_3267 | 1.491 | VK055_4565 | 1.713 |
| VK055_0692 | 0.855 | VK055_1970 | -1.144 | VK055_3268 | -0.283 | VK055_4566 | 0.801 |
| VK055_0693 | -0.819 | VK055_1971 | -0.258 | VK055_3269 | 0.933 | VK055_4572 | -0.560 |
| VK055_0694 | -0.982 | VK055_1972 | -1.171 | VK055_3270 | 0.720 | VK055_4573 | 0.134 |
| VK055_0695 | 0.885 | VK055_1973 | 0.461 | VK055_3271 | -0.150 | VK055_4574 | 0.225 |
| VK055_0696 | 1.408 | VK055_1974 | -0.684 | VK055_3272 | 0.043 | VK055_4575 | -0.187 |
| VK055_0697 | 0.286 | VK055_1976 | 0.204 | VK055_3273 | -1.365 | VK055_4576 | -0.010 |
| VK055_0698 | -0.672 | VK055_1977 | 0.301 | VK055_3275 | 0.359 | VK055_4577 | -0.051 |
| VK055_0701 | -0.287 | VK055_1978 | -0.592 | VK055_3276 | 1.436 | VK055_4578 | 0.862 |
| VK055_0702 | -0.146 | VK055_1979 | -0.705 | VK055_3277 | -0.498 | VK055_4579 | -0.164 |
| VK055_0703 | 0.355 | VK055_1980 | -0.818 | VK055_3279 | 0.161 | VK055_4580 | -0.090 |
| VK055_0704 | -0.212 | VK055_1983 | 0.595 | VK055_3280 | -0.053 | VK055_4581 | 1.350 |
| VK055_0705 | -0.621 | VK055_1984 | 1.434 | VK055_3281 | -0.078 | VK055_4583 | 0.779 |
| VK055_0706 | 0.376 | VK055_1985 | -0.132 | VK055_3282 | -0.000 | VK055_4584 | -0.103 |
| VK055_0707 | 0.845 | VK055_1987 | -0.604 | VK055_3283 | 0.125 | VK055_4585 | -0.366 |
| VK055_0708 | 1.746 | VK055_1991 | 0.054 | VK055_3284 | -1.218 | VK055_4591 | 0.222 |
| VK055_0710 | -0.838 | VK055_1993 | -1.152 | VK055_3285 | -1.133 | VK055_4594 | -0.095 |
| VK055_0712 | 0.476 | VK055_1994 | -1.385 | VK055_3286 | 0.219 | VK055_4595 | 0.402 |
| VK055_0713 | 0.239 | VK055_1995 | 0.235 | VK055_3287 | -0.627 | VK055_4596 | 0.869 |
| VK055_0715 | 0.645 | VK055_1996 | -0.026 | VK055_3288 | 1.547 | VK055_4597 | 0.452 |
| VK055_0716 | 0.014 | VK055_1997 | -1.147 | VK055_3289 | -1.356 | VK055_4598 | -0.895 |
| VK055_0718 | 0.337 | VK055_1999 | 0.339 | VK055_3291 | 0.700 | VK055_4599 | 0.183 |
| VK055_0719 | 1.045 | VK055_2000 | -0.035 | VK055_3292 | 0.924 | VK055_4600 | 0.743 |
| VK055_0721 | -0.077 | VK055_2001 | 0.667 | VK055_3293 | -0.921 | VK055_4601 | -0.193 |
| VK055_0722 | 0.013 | VK055_2002 | -0.281 | VK055_3295 | -0.549 | VK055_4602 | 0.474 |
| VK055_0724 | 0.797 | VK055_2003 | -0.315 | VK055_3296 | 0.114 | VK055_4604 | -0.773 |
| VK055_0725 | -1.029 | VK055_2004 | 1.381 | VK055_3297 | -0.302 | VK055_4605 | 0.421 |
| VK055_0727 | 0.559 | VK055_2005 | -0.295 | VK055_3298 | 0.144 | VK055_4611 | -0.384 |
| VK055_0729 | -0.197 | VK055_2006 | 0.176 | VK055_3299 | -1.440 | VK055_4614 | -0.539 |
| VK055_0730 | -1.645 | VK055_2008 | 0.294 | VK055_3301 | 0.112 | VK055_4615 | -0.313 |
| VK055_0731 | -1.465 | VK055_2009 | -0.358 | VK055_3302 | -1.360 | VK055_4617 | -0.156 |
| VK055_0734 | -0.895 | VK055_2012 | 0.654 | VK055_3303 | -0.951 | VK055_4619 | 1.166 |
| VK055_0735 | -0.013 | VK055_2013 | -0.929 | VK055_3304 | 0.657 | VK055_4620 | -0.999 |
| VK055_0736 | 0.657 | VK055_2014 | 0.999 | VK055_3305 | 0.453 | VK055_4622 | 0.331 |
| VK055_0737 | -0.889 | VK055_2015 | -0.830 | VK055_3306 | 0.449 | VK055_4623 | -0.225 |
| VK055_0738 | 0.317 | VK055_2016 | 0.108 | VK055_3312 | 1.004 | VK055_4624 | 0.560 |
| VK055_0739 | 0.482 | VK055_2017 | -1.073 | VK055_3313 | -1.628 | VK055_4625 | 0.049 |
| VK055_0740 | -0.821 | VK055_2018 | 0.151 | VK055_3314 | -0.906 | VK055_4628 | 0.937 |
| VK055_0741 | -0.250 | VK055_2019 | 0.560 | VK055_3315 | -1.320 | VK055_4629 | 0.430 |
| VK055_0743 | 0.421 | VK055_2020 | -0.514 | VK055_3316 | -0.372 | VK055_4630 | -0.434 |
| VK055_0744 | 0.057 | VK055_2021 | 1.452 | VK055_3317 | 0.586 | VK055_4631 | -1.429 |
| VK055_0745 | -0.245 | VK055_2023 | -0.871 | VK055_3318 | -0.797 | VK055_4632 | 1.038 |
| VK055_0746 | 2.422 | VK055_2025 | 0.211 | VK055_3319 | -1.440 | VK055_4633 | 0.639 |
| VK055_0749 | -0.497 | VK055_2026 | 1.836 | VK055_3320 | -1.066 | VK055_4635 | 0.647 |
| VK055_0750 | 0.261 | VK055_2027 | 0.961 | VK055_3322 | -0.823 | VK055_4637 | 1.158 |
| VK055_0753 | -0.474 | VK055_2029 | 0.112 | VK055_3323 | 0.263 | VK055_4646 | -0.111 |
| VK055_0754 | -0.446 | VK055_2031 | 0.922 | VK055_3324 | 0.097 | VK055_4647 | 0.304 |
| VK055_0755 | 0.073 | VK055_2033 | 0.253 | VK055_3325 | -0.342 | VK055_4648 | 1.114 |
| VK055_0756 | -0.664 | VK055_2034 | 0.029 | VK055_3326 | 0.603 | VK055_4649 | 0.152 |
| VK055_0757 | 0.773 | VK055_2035 | 0.330 | VK055_3327 | -0.548 | VK055_4650 | -0.019 |
| VK055_0758 | -0.375 | VK055_2036 | 1.041 | VK055_3328 | 0.780 | VK055_4651 | 0.153 |
| VK055_0761 | 0.382 | VK055_2037 | 0.884 | VK055_3338 | 0.434 | VK055_4653 | -0.251 |
| VK055_0762 | -0.756 | VK055_2040 | -0.146 | VK055_3342 | -0.329 | VK055_4658 | 0.097 |
| VK055_0763 | 0.707 | VK055_2041 | -1.230 | VK055_3344 | -1.497 | VK055_4660 | 0.987 |
| VK055_0764 | 0.147 | VK055_2043 | -0.180 | VK055_3347 | 1.271 | VK055_4662 | -0.718 |
| VK055_0765 | -0.088 | VK055_2046 | -0.176 | VK055_3348 | -0.078 | VK055_4663 | 0.124 |
| VK055_0767 | 0.116 | VK055_2047 | -0.765 | VK055_3349 | -0.765 | VK055_4664 | -1.216 |
| VK055_0770 | -0.494 | VK055_2048 | 0.114 | VK055_3350 | -0.095 | VK055_4665 | 0.131 |
| VK055_0773 | -1.044 | VK055_2049 | -1.159 | VK055_3351 | -0.902 | VK055_4667 | 0.896 |
| VK055_0775 | 1.242 | VK055_2050 | 0.322 | VK055_3352 | -1.428 | VK055_4668 | -0.231 |
| VK055_0776 | -0.792 | VK055_2051 | 0.333 | VK055_3353 | 0.950 | VK055_4669 | -0.502 |
| VK055_0777 | -1.477 | VK055_2052 | -0.605 | VK055_3354 | -0.315 | VK055_4671 | 0.382 |
| VK055_0778 | -0.222 | VK055_2053 | -0.628 | VK055_3355 | -0.190 | VK055_4672 | -0.965 |
| VK055_0779 | -1.464 | VK055_2055 | 1.080 | VK055_3356 | 0.044 | VK055_4673 | 1.474 |
| VK055_0780 | -0.162 | VK055_2056 | -0.158 | VK055_3357 | -0.326 | VK055_4674 | -0.455 |
| VK055_0781 | 0.922 | VK055_2057 | -0.463 | VK055_3361 | -0.631 | VK055_4676 | 0.983 |
| VK055_0782 | -0.554 | VK055_2058 | -1.437 | VK055_3363 | -0.702 | VK055_4677 | -1.165 |
| VK055_0784 | -0.567 | VK055_2059 | 0.256 | VK055_3364 | -1.065 | VK055_4678 | -0.891 |
| VK055_0785 | -0.045 | VK055_2060 | 0.563 | VK055_3365 | 0.467 | VK055_4679 | 0.868 |
| VK055_0786 | 0.894 | VK055_2061 | -0.239 | VK055_3366 | 0.126 | VK055_4680 | 1.151 |
| VK055_0787 | 0.346 | VK055_2065 | -0.677 | VK055_3367 | -0.254 | VK055_4681 | -1.470 |
| VK055_0788 | 0.086 | VK055_2068 | 0.070 | VK055_3368 | 0.337 | VK055_4682 | -1.477 |
| VK055_0789 | 0.183 | VK055_2070 | 0.393 | VK055_3369 | -1.452 | VK055_4683 | 0.524 |
| VK055_0790 | -1.457 | VK055_2071 | -1.203 | VK055_3370 | 0.797 | VK055_4684 | -0.063 |
| VK055_0791 | -0.536 | VK055_2072 | -0.920 | VK055_3371 | 0.115 | VK055_4685 | 6.350 |
| VK055_0792 | 0.081 | VK055_2073 | -1.458 | VK055_3372 | 0.192 | VK055_4686 | 1.436 |
| VK055_0793 | 0.291 | VK055_2075 | -0.104 | VK055_3373 | -0.096 | VK055_4687 | -0.435 |
| VK055_0794 | -0.093 | VK055_2076 | 0.102 | VK055_3376 | -0.405 | VK055_4688 | -0.291 |
| VK055_0795 | -0.208 | VK055_2077 | 1.783 | VK055_3377 | 0.271 | VK055_4689 | 1.558 |
| VK055_0798 | -0.308 | VK055_2078 | -0.749 | VK055_3378 | -1.409 | VK055_4690 | 0.175 |
| VK055_0799 | 0.643 | VK055_2079 | 0.779 | VK055_3379 | 0.132 | VK055_4691 | 0.598 |
| VK055_0800 | -0.554 | VK055_2080 | 0.212 | VK055_3380 | 1.017 | VK055_4693 | 0.229 |
| VK055_0802 | -0.113 | VK055_2081 | 0.853 | VK055_3381 | 0.299 | VK055_4694 | -0.153 |
| VK055_0803 | -0.547 | VK055_2082 | -0.367 | VK055_3383 | 0.865 | VK055_4695 | -0.674 |
| VK055_0807 | 0.086 | VK055_2084 | -0.210 | VK055_3384 | -0.598 | VK055_4696 | -0.748 |
| VK055_0808 | -0.406 | VK055_2085 | -0.778 | VK055_3385 | 1.657 | VK055_4697 | -0.298 |
| VK055_0809 | -0.474 | VK055_2087 | -0.057 | VK055_3386 | 0.076 | VK055_4698 | 1.348 |
| VK055_0810 | -0.703 | VK055_2088 | -1.479 | VK055_3387 | 0.584 | VK055_4700 | -0.706 |
| VK055_0811 | -0.412 | VK055_2089 | -0.567 | VK055_3388 | -0.727 | VK055_4701 | -0.337 |
| VK055_0812 | 0.245 | VK055_2090 | 0.096 | VK055_3389 | 0.320 | VK055_4702 | -0.497 |
| VK055_0814 | 1.317 | VK055_2091 | -0.389 | VK055_3391 | -0.182 | VK055_4703 | 0.231 |
| VK055_0815 | 1.027 | VK055_2094 | 0.317 | VK055_3392 | 0.529 | VK055_4707 | -0.514 |
| VK055_0816 | -0.203 | VK055_2098 | 0.227 | VK055_3393 | -0.838 | VK055_4708 | -0.450 |
| VK055_0817 | -0.519 | VK055_2100 | -0.833 | VK055_3395 | -0.272 | VK055_4709 | -0.855 |
| VK055_0818 | 0.730 | VK055_2101 | -0.150 | VK055_3396 | 1.091 | VK055_4710 | 0.572 |
| VK055_0819 | 0.275 | VK055_2102 | -0.446 | VK055_3397 | 0.103 | VK055_4711 | -0.250 |
| VK055_0821 | 0.055 | VK055_2103 | 0.046 | VK055_3399 | 0.101 | VK055_4712 | 1.057 |
| VK055_0822 | 0.263 | VK055_2104 | 0.885 | VK055_3400 | 1.506 | VK055_4713 | -1.416 |
| VK055_0823 | 0.108 | VK055_2104 | -0.057 | VK055_3401 | 0.748 | VK055_4714 | 0.606 |
| VK055_0824 | 0.801 | VK055_2105 | 0.255 | VK055_3402 | -0.239 | VK055_4716 | -1.260 |
| VK055_0826 | -0.518 | VK055_2106 | 0.678 | VK055_3403 | -0.928 | VK055_4718 | -0.412 |
| VK055_0827 | 0.287 | VK055_2107 | 0.326 | VK055_3404 | -0.115 | VK055_4720 | -0.653 |
| VK055_0828 | -0.152 | VK055_2108 | 0.985 | VK055_3405 | 0.468 | VK055_4721 | 0.847 |
| VK055_0829 | 0.041 | VK055_2110 | -1.353 | VK055_3406 | 0.047 | VK055_4722 | -1.497 |
| VK055_0830 | -1.096 | VK055_2111 | -0.873 | VK055_3407 | -0.431 | VK055_4723 | 0.697 |
| VK055_0831 | -1.461 | VK055_2112 | 0.074 | VK055_3409 | 1.177 | VK055_4724 | 0.276 |
| VK055_0833 | -1.457 | VK055_2113 | 0.823 | VK055_3410 | -0.072 | VK055_4725 | 0.247 |
| VK055_0834 | 0.293 | VK055_2114 | -0.394 | VK055_3411 | -0.227 | VK055_4726 | -0.289 |
| VK055_0835 | 1.304 | VK055_2115 | -1.025 | VK055_3412 | 0.244 | VK055_4727 | 0.262 |
| VK055_0837 | 0.950 | VK055_2117 | 0.361 | VK055_3413 | 1.086 | VK055_4728 | -0.110 |
| VK055_0838 | 0.048 | VK055_2118 | 0.504 | VK055_3414 | -0.581 | VK055_4729 | -0.412 |
| VK055_0839 | 0.211 | VK055_2119 | 0.603 | VK055_3415 | -0.735 | VK055_4731 | -1.422 |
| VK055_0840 | 0.485 | VK055_2120 | -0.405 | VK055_3416 | 1.051 | VK055_4732 | -0.530 |
| VK055_0841 | 0.428 | VK055_2121 | -0.086 | VK055_3417 | -0.161 | VK055_4733 | -1.149 |
| VK055_0842 | -0.336 | VK055_2122 | -0.616 | VK055_3419 | -0.152 | VK055_4734 | 0.863 |
| VK055_0849 | -0.319 | VK055_2123 | 0.100 | VK055_3420 | 1.064 | VK055_4737 | 1.541 |
| VK055_0850 | -1.031 | VK055_2124 | 0.779 | VK055_3421 | 0.728 | VK055_4738 | 0.746 |
| VK055_0852 | -0.362 | VK055_2125 | -1.217 | VK055_3422 | -0.410 | VK055_4739 | -0.480 |
| VK055_0854 | -0.718 | VK055_2127 | 0.149 | VK055_3423 | -0.806 | VK055_4740 | 1.485 |
| VK055_0855 | -0.034 | VK055_2129 | 0.080 | VK055_3424 | 0.614 | VK055_4741 | 1.079 |
| VK055_0856 | 0.057 | VK055_2130 | -0.118 | VK055_3425 | -0.432 | VK055_4742 | -0.767 |
| VK055_0860 | 0.260 | VK055_2131 | -0.257 | VK055_3426 | -0.254 | VK055_4743 | 0.139 |
| VK055_0861 | -0.660 | VK055_2132 | 1.055 | VK055_3427 | -0.019 | VK055_4744 | -0.071 |
| VK055_0862 | 0.502 | VK055_2134 | -0.713 | VK055_3429 | -0.281 | VK055_4745 | 0.550 |
| VK055_0863 | -0.060 | VK055_2135 | 0.519 | VK055_3430 | -0.514 | VK055_4746 | -0.667 |
| VK055_0864 | -0.065 | VK055_2136 | -0.714 | VK055_3431 | 0.294 | VK055_4748 | -0.913 |
| VK055_0865 | -0.084 | VK055_2137 | -0.794 | VK055_3432 | 0.484 | VK055_4749 | 1.444 |
| VK055_0866 | -1.236 | VK055_2138 | -0.545 | VK055_3433 | 0.160 | VK055_4750 | -0.492 |
| VK055_0867 | -1.269 | VK055_2139 | 0.151 | VK055_3434 | -0.932 | VK055_4751 | -0.305 |
| VK055_0868 | 0.076 | VK055_2140 | 0.503 | VK055_3436 | 0.755 | VK055_4752 | -0.816 |
| VK055_0870 | -0.035 | VK055_2141 | 1.049 | VK055_3437 | -0.346 | VK055_4755 | 0.887 |
| VK055_0871 | 1.872 | VK055_2142 | -0.090 | VK055_3438 | -0.748 | VK055_4756 | 1.471 |
| VK055_0874 | -0.565 | VK055_2143 | 1.308 | VK055_3439 | 0.444 | VK055_4757 | -0.846 |
| VK055_0875 | -0.096 | VK055_2144 | -0.178 | VK055_3440 | 0.689 | VK055_4764 | 0.314 |
| VK055_0876 | 1.124 | VK055_2145 | 4.107 | VK055_3441 | -0.145 | VK055_4767 | -0.762 |
| VK055_0878 | -0.065 | VK055_2146 | -0.488 | VK055_3443 | 2.567 | VK055_4768 | -0.027 |
| VK055_0880 | 0.091 | VK055_2147 | 0.017 | VK055_3444 | 2.166 | VK055_4769 | -0.320 |
| VK055_0882 | 0.441 | VK055_2150 | 1.020 | VK055_3445 | 1.463 | VK055_4770 | 0.936 |
| VK055_0883 | -1.471 | VK055_2152 | -1.497 | VK055_3448 | 0.799 | VK055_4771 | -1.079 |
| VK055_0884 | 0.803 | VK055_2153 | 0.194 | VK055_3449 | 0.667 | VK055_4772 | -0.927 |
| VK055_0885 | 0.555 | VK055_2154 | -0.113 | VK055_3450 | 0.918 | VK055_4773 | 0.235 |
| VK055_0887 | 0.523 | VK055_2155 | 0.303 | VK055_3451 | -0.240 | VK055_4774 | 0.055 |
| VK055_0888 | 1.139 | VK055_2157 | 0.886 | VK055_3452 | -0.464 | VK055_4775 | 0.536 |
| VK055_0889 | -0.008 | VK055_2159 | -0.263 | VK055_3453 | 0.002 | VK055_4776 | 0.250 |
| VK055_0891 | 0.383 | VK055_2160 | -0.118 | VK055_3454 | 0.394 | VK055_4778 | -0.723 |
| VK055_0892 | 0.627 | VK055_2161 | 0.121 | VK055_3456 | 0.900 | VK055_4779 | 0.544 |
| VK055_0893 | -0.385 | VK055_2162 | -0.359 | VK055_3457 | 1.128 | VK055_4780 | -1.491 |
| VK055_0894 | -1.256 | VK055_2164 | -0.356 | VK055_3458 | 0.047 | VK055_4781 | -0.293 |
| VK055_0895 | 0.920 | VK055_2165 | -0.683 | VK055_3460 | 0.530 | VK055_4782 | 0.478 |
| VK055_0896 | -0.345 | VK055_2167 | 1.308 | VK055_3461 | -1.216 | VK055_4783 | -1.483 |
| VK055_0897 | -0.275 | VK055_2170 | 0.332 | VK055_3462 | 0.844 | VK055_4784 | 1.888 |
| VK055_0898 | 0.978 | VK055_2171 | -0.340 | VK055_3463 | 0.494 | VK055_4785 | -0.584 |
| VK055_0899 | -1.253 | VK055_2172 | 0.355 | VK055_3465 | -0.651 | VK055_4786 | 0.922 |
| VK055_0900 | 0.647 | VK055_2173 | 0.228 | VK055_3466 | 0.721 | VK055_4787 | -0.392 |
| VK055_0907 | -0.118 | VK055_2174 | -0.347 | VK055_3467 | -1.497 | VK055_4789 | 1.134 |
| VK055_0909 | -1.416 | VK055_2176 | 0.629 | VK055_3468 | -0.294 | VK055_4790 | -1.476 |
| VK055_0910 | -0.449 | VK055_2177 | -1.292 | VK055_3469 | -0.077 | VK055_4793 | 0.032 |
| VK055_0911 | -0.172 | VK055_2179 | 1.144 | VK055_3470 | -0.846 | VK055_4794 | 0.169 |
| VK055_0913 | 0.371 | VK055_2182 | -0.926 | VK055_3473 | 0.515 | VK055_4795 | -0.324 |
| VK055_0914 | 2.025 | VK055_2183 | -1.201 | VK055_3474 | -1.490 | VK055_4796 | 0.060 |
| VK055_0915 | 0.577 | VK055_2190 | -0.000 | VK055_3475 | -0.191 | VK055_4797 | 0.198 |
| VK055_0917 | 0.426 | VK055_2191 | 0.546 | VK055_3476 | -0.080 | VK055_4799 | -0.244 |
| VK055_0918 | 0.028 | VK055_2192 | 0.736 | VK055_3477 | 0.078 | VK055_4801 | 0.020 |
| VK055_0919 | 0.752 | VK055_2193 | 0.473 | VK055_3478 | 0.232 | VK055_4803 | -1.032 |
| VK055_0920 | -0.036 | VK055_2194 | -0.863 | VK055_3480 | 0.578 | VK055_4804 | -0.419 |
| VK055_0921 | 0.577 | VK055_2197 | 1.397 | VK055_3482 | 0.724 | VK055_4805 | 1.238 |
| VK055_0922 | 0.190 | VK055_2198 | -0.062 | VK055_3483 | -0.292 | VK055_4806 | 0.647 |
| VK055_0924 | 0.561 | VK055_2199 | 1.402 | VK055_3484 | -0.794 | VK055_4811 | 0.008 |
| VK055_0925 | -0.191 | VK055_2200 | -0.800 | VK055_3485 | 0.112 | VK055_4813 | 0.807 |
| VK055_0927 | 0.888 | VK055_2201 | 0.239 | VK055_3486 | -0.777 | VK055_4814 | 1.177 |
| VK055_0928 | 0.937 | VK055_2202 | -0.478 | VK055_3487 | -0.777 | VK055_4815 | 0.026 |
| VK055_0929 | -0.061 | VK055_2203 | -0.200 | VK055_3488 | 0.564 | VK055_4816 | -0.144 |
| VK055_0930 | -0.050 | VK055_2204 | -1.057 | VK055_3491 | 0.549 | VK055_4817 | -1.195 |
| VK055_0933 | 0.071 | VK055_2205 | -0.152 | VK055_3493 | 1.170 | VK055_4820 | -0.241 |
| VK055_0934 | -0.357 | VK055_2206 | -0.899 | VK055_3494 | 0.226 | VK055_4822 | 2.448 |
| VK055_0936 | 1.000 | VK055_2207 | 1.128 | VK055_3498 | -0.034 | VK055_4823 | 0.660 |
| VK055_0938 | -0.009 | VK055_2209 | -1.491 | VK055_3503 | -1.156 | VK055_4824 | -0.295 |
| VK055_0939 | -0.815 | VK055_2210 | 0.424 | VK055_3509 | 0.249 | VK055_4825 | 0.997 |
| VK055_0941 | -0.206 | VK055_2211 | 0.289 | VK055_3510 | -1.155 | VK055_4826 | -0.908 |
| VK055_0942 | 0.667 | VK055_2212 | 0.122 | VK055_3511 | 0.904 | VK055_4827 | -1.434 |
| VK055_0944 | 0.396 | VK055_2213 | 0.567 | VK055_3512 | -0.532 | VK055_4828 | -0.030 |
| VK055_0945 | 2.225 | VK055_2215 | -0.616 | VK055_3513 | -1.143 | VK055_4829 | 0.005 |
| VK055_0946 | 0.414 | VK055_2216 | 0.776 | VK055_3514 | -0.288 | VK055_4831 | 1.337 |
| VK055_0948 | -1.295 | VK055_2217 | -0.006 | VK055_3515 | -0.369 | VK055_4832 | -0.743 |
| VK055_0949 | -1.451 | VK055_2218 | -0.024 | VK055_3519 | -0.326 | VK055_4833 | 0.043 |
| VK055_0950 | -0.520 | VK055_2221 | 0.563 | VK055_3521 | -1.465 | VK055_4834 | 0.957 |
| VK055_0951 | 0.077 | VK055_2222 | -0.452 | VK055_3522 | 0.211 | VK055_4835 | 0.905 |
| VK055_0952 | 0.093 | VK055_2224 | 1.253 | VK055_3523 | 0.437 | VK055_4836 | 0.816 |
| VK055_0953 | -0.473 | VK055_2225 | -0.180 | VK055_3525 | -0.225 | VK055_4837 | -1.490 |
| VK055_0954 | -0.221 | VK055_2226 | -0.228 | VK055_3527 | -0.304 | VK055_4838 | -0.449 |
| VK055_0955 | -0.896 | VK055_2227 | -0.903 | VK055_3528 | 0.749 | VK055_4839 | 0.553 |
| VK055_0956 | 0.358 | VK055_2228 | 0.081 | VK055_3531 | -0.399 | VK055_4840 | 0.979 |
| VK055_0957 | -0.073 | VK055_2229 | 1.388 | VK055_3533 | -0.233 | VK055_4842 | 0.850 |
| VK055_0958 | 0.775 | VK055_2231 | -0.080 | VK055_3534 | 1.314 | VK055_4843 | 0.281 |
| VK055_0959 | -0.622 | VK055_2232 | -0.447 | VK055_3535 | -1.469 | VK055_4844 | 0.196 |
| VK055_0960 | -0.124 | VK055_2233 | -0.616 | VK055_3536 | -1.605 | VK055_4845 | 0.094 |
| VK055_0961 | -0.734 | VK055_2234 | -0.859 | VK055_3537 | -0.312 | VK055_4846 | 1.185 |
| VK055_0963 | 0.503 | VK055_2235 | 1.384 | VK055_3538 | 0.557 | VK055_4847 | 0.368 |
| VK055_0965 | -0.994 | VK055_2237 | -0.682 | VK055_3539 | 1.730 | VK055_4848 | -0.023 |
| VK055_0966 | 0.209 | VK055_2238 | -0.467 | VK055_3540 | -0.544 | VK055_4849 | -0.186 |
| VK055_0967 | 0.027 | VK055_2239 | 0.008 | VK055_3541 | -0.175 | VK055_4850 | 0.237 |
| VK055_0968 | 0.638 | VK055_2241 | -0.667 | VK055_3542 | -0.080 | VK055_4851 | 0.478 |
| VK055_0969 | 0.219 | VK055_2242 | 3.834 | VK055_3543 | 0.102 | VK055_4852 | -0.063 |
| VK055_0970 | -0.685 | VK055_2243 | 0.146 | VK055_3544 | 0.538 | VK055_4853 | -0.786 |
| VK055_0971 | 0.111 | VK055_2245 | 0.158 | VK055_3545 | -0.268 | VK055_4854 | -0.593 |
| VK055_0972 | 0.445 | VK055_2246 | -0.153 | VK055_3549 | -0.455 | VK055_4855 | -0.207 |
| VK055_0975 | 0.657 | VK055_2247 | 0.279 | VK055_3550 | -1.440 | VK055_4857 | 0.101 |
| VK055_0977 | -0.736 | VK055_2248 | 0.450 | VK055_3551 | -1.445 | VK055_4858 | -1.054 |
| VK055_0978 | -0.270 | VK055_2249 | -0.307 | VK055_3552 | 0.847 | VK055_4859 | 0.956 |
| VK055_0980 | -0.410 | VK055_2250 | 1.298 | VK055_3553 | 0.519 | VK055_4860 | 1.097 |
| VK055_0981 | 0.893 | VK055_2251 | 0.690 | VK055_3554 | 1.267 | VK055_4861 | -0.028 |
| VK055_0982 | 0.738 | VK055_2252 | -1.478 | VK055_3555 | 0.827 | VK055_4862 | 0.681 |
| VK055_0983 | -0.783 | VK055_2253 | -1.273 | VK055_3556 | 0.336 | VK055_4863 | -0.843 |
| VK055_0984 | -0.096 | VK055_2254 | 0.697 | VK055_3558 | 0.058 | VK055_4864 | 0.346 |
| VK055_0985 | 0.896 | VK055_2255 | -0.602 | VK055_3559 | 0.904 | VK055_4865 | 0.112 |
| VK055_0986 | 0.147 | VK055_2256 | 0.664 | VK055_3561 | 0.133 | VK055_4866 | -0.817 |
| VK055_0988 | -0.019 | VK055_2257 | -0.943 | VK055_3562 | -1.478 | VK055_4867 | 0.541 |
| VK055_0989 | 0.001 | VK055_2258 | -0.639 | VK055_3563 | 0.377 | VK055_4868 | -1.309 |
| VK055_0990 | -0.018 | VK055_2259 | 0.211 | VK055_3564 | 0.225 | VK055_4869 | -0.153 |
| VK055_0992 | -0.162 | VK055_2261 | -0.403 | VK055_3565 | -0.603 | VK055_4870 | -0.183 |
| VK055_0993 | 1.675 | VK055_2262 | -0.558 | VK055_3566 | 0.616 | VK055_4877 | 0.025 |
| VK055_0995 | 0.650 | VK055_2263 | -0.840 | VK055_3568 | 0.204 | VK055_4878 | 0.191 |
| VK055_0996 | 0.394 | VK055_2264 | -0.780 | VK055_3569 | 0.142 | VK055_4879 | 0.136 |
| VK055_1000 | -0.979 | VK055_2265 | -0.464 | VK055_3570 | 2.505 | VK055_4880 | 0.568 |
| VK055_1001 | -1.422 | VK055_2266 | 0.294 | VK055_3571 | -0.676 | VK055_4881 | -0.595 |
| VK055_1002 | -1.408 | VK055_2267 | 1.634 | VK055_3572 | -0.275 | VK055_4882 | 0.458 |
| VK055_1004 | 0.097 | VK055_2268 | -0.205 | VK055_3575 | 0.523 | VK055_4884 | -0.487 |
| VK055_1005 | 0.892 | VK055_2270 | 0.916 | VK055_3576 | -0.761 | VK055_4885 | 0.152 |
| VK055_1006 | 0.042 | VK055_2271 | 0.666 | VK055_3577 | 0.417 | VK055_4886 | -0.739 |
| VK055_1007 | 1.215 | VK055_2275 | -0.817 | VK055_3578 | 0.378 | VK055_4887 | 0.019 |
| VK055_1009 | 0.119 | VK055_2277 | 0.187 | VK055_3579 | -1.428 | VK055_4889 | 1.298 |
| VK055_1010 | -0.058 | VK055_2278 | 1.111 | VK055_3580 | -0.643 | VK055_4890 | -0.338 |
| VK055_1011 | 0.403 | VK055_2279 | 0.440 | VK055_3581 | -0.126 | VK055_4891 | -0.181 |
| VK055_1012 | -0.836 | VK055_2280 | 0.641 | VK055_3582 | -0.691 | VK055_4892 | 0.250 |
| VK055_1013 | -1.462 | VK055_2281 | 0.226 | VK055_3583 | 1.898 | VK055_4893 | 0.146 |
| VK055_1014 | 0.168 | VK055_2285 | 0.890 | VK055_3585 | -0.464 | VK055_4894 | 0.530 |
| VK055_1015 | 0.570 | VK055_2286 | 0.790 | VK055_3586 | -0.030 | VK055_4896 | 0.750 |
| VK055_1016 | 1.491 | VK055_2287 | -0.347 | VK055_3587 | -0.612 | VK055_4900 | 0.871 |
| VK055_1018 | 0.310 | VK055_2288 | -0.053 | VK055_3588 | 0.330 | VK055_4901 | -0.062 |
| VK055_1019 | 0.195 | VK055_2292 | -1.139 | VK055_3589 | -0.250 | VK055_4902 | -1.307 |
| VK055_1020 | -0.479 | VK055_2293 | -0.726 | VK055_3590 | 0.474 | VK055_4904 | 0.391 |
| VK055_1021 | -0.213 | VK055_2294 | -0.518 | VK055_3591 | -0.124 | VK055_4905 | -0.712 |
| VK055_1022 | 0.649 | VK055_2295 | -0.039 | VK055_3592 | -0.150 | VK055_4906 | -0.165 |
| VK055_1023 | -0.145 | VK055_2296 | 1.438 | VK055_3593 | -0.341 | VK055_4907 | 1.413 |
| VK055_1024 | -1.153 | VK055_2298 | -0.157 | VK055_3594 | -0.369 | VK055_4908 | 0.574 |
| VK055_1026 | 0.621 | VK055_2300 | -0.358 | VK055_3595 | 0.528 | VK055_4909 | -1.464 |
| VK055_1027 | 0.092 | VK055_2301 | 0.023 | VK055_3596 | -0.489 | VK055_4910 | 0.001 |
| VK055_1029 | 0.918 | VK055_2302 | 0.513 | VK055_3597 | -0.760 | VK055_4911 | 0.346 |
| VK055_1031 | 2.478 | VK055_2303 | 1.704 | VK055_3598 | 0.032 | VK055_4912 | -0.168 |
| VK055_1032 | -1.440 | VK055_2304 | -0.857 | VK055_3600 | -1.206 | VK055_4913 | -0.011 |
| VK055_1033 | 2.399 | VK055_2305 | 0.018 | VK055_3601 | 0.354 | VK055_4914 | 0.186 |
| VK055_1034 | -0.669 | VK055_2306 | -0.011 | VK055_3603 | -0.419 | VK055_4915 | -1.457 |
| VK055_1035 | -0.264 | VK055_2307 | -1.277 | VK055_3604 | -0.554 | VK055_4917 | -0.802 |
| VK055_1038 | -0.806 | VK055_2308 | 1.372 | VK055_3605 | -0.332 | VK055_4918 | -0.135 |
| VK055_1039 | -0.014 | VK055_2309 | 0.567 | VK055_3606 | -0.273 | VK055_4919 | 0.366 |
| VK055_1040 | -0.580 | VK055_2310 | 0.146 | VK055_3607 | -1.192 | VK055_4920 | -0.485 |
| VK055_1042 | 0.299 | VK055_2311 | -0.739 | VK055_3608 | 1.368 | VK055_4921 | 0.678 |
| VK055_1043 | 1.746 | VK055_2312 | 1.183 | VK055_3609 | 1.181 | VK055_4923 | -0.550 |
| VK055_1044 | -0.380 | VK055_2313 | -1.440 | VK055_3610 | 0.764 | VK055_4924 | -0.611 |
| VK055_1045 | 0.015 | VK055_2315 | 0.943 | VK055_3611 | 0.010 | VK055_4925 | -1.434 |
| VK055_1046 | 0.655 | VK055_2317 | -0.567 | VK055_3612 | -1.362 | VK055_4926 | -0.502 |
| VK055_1048 | 0.753 | VK055_2319 | 0.554 | VK055_3613 | -0.659 | VK055_4927 | 0.234 |
| VK055_1049 | 0.163 | VK055_2322 | 0.002 | VK055_3614 | -1.152 | VK055_4928 | -0.745 |
| VK055_1050 | -1.046 | VK055_2323 | 0.483 | VK055_3615 | -1.497 | VK055_4930 | 1.566 |
| VK055_1052 | -0.156 | VK055_2325 | 0.883 | VK055_3617 | -1.404 | VK055_4933 | 0.675 |
| VK055_1053 | 1.080 | VK055_2326 | -0.203 | VK055_3618 | -0.392 | VK055_4934 | 0.716 |
| VK055_1056 | 0.106 | VK055_2327 | 0.240 | VK055_3619 | -0.018 | VK055_4935 | 0.347 |
| VK055_1057 | 0.433 | VK055_2328 | -0.289 | VK055_3620 | -0.074 | VK055_4936 | -0.297 |
| VK055_1058 | 0.012 | VK055_2329 | -1.162 | VK055_3621 | 0.037 | VK055_4937 | 0.078 |
| VK055_1061 | -0.314 | VK055_2331 | -1.446 | VK055_3622 | -0.740 | VK055_4938 | 1.387 |
| VK055_1062 | -0.630 | VK055_2332 | 0.761 | VK055_3623 | -1.061 | VK055_4939 | 0.305 |
| VK055_1063 | 1.080 | VK055_2333 | 0.338 | VK055_3624 | 0.862 | VK055_4940 | 1.297 |
| VK055_1064 | -0.303 | VK055_2334 | 0.394 | VK055_3625 | -1.103 | VK055_4941 | 0.306 |
| VK055_1065 | -0.501 | VK055_2337 | 0.613 | VK055_3626 | -0.378 | VK055_4942 | 1.049 |
| VK055_1067 | -0.604 | VK055_2339 | 0.032 | VK055_3627 | -0.208 | VK055_4943 | 0.273 |
| VK055_1068 | 0.983 | VK055_2340 | 0.680 | VK055_3628 | -0.646 | VK055_4944 | 0.607 |
| VK055_1069 | 0.421 | VK055_2341 | 0.225 | VK055_3629 | -0.940 | VK055_4945 | 0.176 |
| VK055_1070 | -0.038 | VK055_2342 | 0.843 | VK055_3630 | -0.051 | VK055_4946 | 0.063 |
| VK055_1071 | -0.335 | VK055_2343 | 0.220 | VK055_3631 | 0.019 | VK055_4948 | -0.080 |
| VK055_1072 | 1.477 | VK055_2344 | 1.039 | VK055_3632 | 1.335 | VK055_4949 | -1.382 |
| VK055_1073 | -0.220 | VK055_2345 | -0.624 | VK055_3633 | 0.718 | VK055_4950 | 0.351 |
| VK055_1074 | -0.322 | VK055_2352 | -1.369 | VK055_3635 | -1.445 | VK055_4951 | -0.654 |
| VK055_1075 | -0.112 | VK055_2353 | 0.731 | VK055_3636 | 1.299 | VK055_4952 | -0.382 |
| VK055_1076 | 0.092 | VK055_2354 | -0.717 | VK055_3637 | 1.375 | VK055_4953 | 0.727 |
| VK055_1078 | 0.396 | VK055_2355 | -0.135 | VK055_3639 | 0.337 | VK055_4954 | 0.080 |
| VK055_1079 | -1.636 | VK055_2356 | 0.760 | VK055_3644 | 0.859 | VK055_4955 | -0.529 |
| VK055_1081 | 0.169 | VK055_2357 | 0.969 | VK055_3646 | 0.292 | VK055_4957 | -1.216 |
| VK055_1082 | -0.410 | VK055_2359 | 0.433 | VK055_3647 | -0.038 | VK055_4958 | 0.344 |
| VK055_1084 | 0.335 | VK055_2360 | -0.270 | VK055_3648 | -0.658 | VK055_4960 | -0.118 |
| VK055_1085 | -0.722 | VK055_2362 | 0.599 | VK055_3650 | -0.485 | VK055_4961 | -0.167 |
| VK055_1086 | -0.895 | VK055_2365 | 0.122 | VK055_3651 | 0.906 | VK055_4962 | -0.269 |
| VK055_1088 | -1.263 | VK055_2366 | -1.127 | VK055_3652 | -0.788 | VK055_4963 | 0.101 |
| VK055_1090 | -1.380 | VK055_2373 | -0.607 | VK055_3653 | 1.360 | VK055_4964 | 0.434 |
| VK055_1091 | -0.187 | VK055_2384 | 0.490 | VK055_3654 | 1.176 | VK055_4966 | -1.140 |
| VK055_1092 | 0.367 | VK055_2385 | -0.166 | VK055_3655 | 0.482 | VK055_4967 | -0.090 |
| VK055_1093 | -0.647 | VK055_2390 | -0.311 | VK055_3656 | 0.448 | VK055_4968 | 0.263 |
| VK055_1094 | 0.144 | VK055_2392 | -0.241 | VK055_3657 | -0.215 | VK055_4969 | 0.227 |
| VK055_1095 | 0.599 | VK055_2395 | 0.289 | VK055_3658 | 0.910 | VK055_4970 | -0.838 |
| VK055_1096 | 0.542 | VK055_2397 | 0.065 | VK055_3659 | -1.416 | VK055_4971 | 1.831 |
| VK055_1097 | 0.538 | VK055_2399 | 0.327 | VK055_3660 | 1.017 | VK055_4972 | -0.734 |
| VK055_1099 | 0.101 | VK055_2400 | 0.267 | VK055_3661 | -0.112 | VK055_4973 | -0.235 |
| VK055_1100 | -1.139 | VK055_2401 | 0.045 | VK055_3662 | -0.296 | VK055_4977 | 1.799 |
| VK055_1101 | -0.574 | VK055_2402 | -1.504 | VK055_3663 | -0.484 | VK055_4978 | 0.306 |
| VK055_1102 | 0.375 | VK055_2404 | 0.110 | VK055_3664 | 0.853 | VK055_4979 | 1.047 |
| VK055_1103 | -0.626 | VK055_2405 | 0.272 | VK055_3665 | 0.669 | VK055_4980 | -0.565 |
| VK055_1104 | 0.937 | VK055_2407 | -0.387 | VK055_3666 | 0.177 | VK055_4981 | 0.174 |
| VK055_1105 | 0.543 | VK055_2408 | -0.662 | VK055_3667 | 1.354 | VK055_4982 | 0.520 |
| VK055_1106 | 0.974 | VK055_2409 | -0.813 | VK055_3668 | -0.272 | VK055_4983 | 0.382 |
| VK055_1108 | 0.619 | VK055_2410 | 0.372 | VK055_3669 | 0.067 | VK055_4985 | -0.146 |
| VK055_1109 | -1.409 | VK055_2412 | 0.272 | VK055_3670 | -1.022 | VK055_4986 | 0.395 |
| VK055_1110 | -0.184 | VK055_2415 | 0.140 | VK055_3671 | -0.197 | VK055_4987 | -0.716 |
| VK055_1111 | 0.395 | VK055_2416 | -0.377 | VK055_3673 | -0.576 | VK055_4988 | 0.800 |
| VK055_1113 | 0.895 | VK055_2418 | -0.035 | VK055_3674 | 0.612 | VK055_4989 | -1.491 |
| VK055_1114 | 0.987 | VK055_2420 | 1.193 | VK055_3675 | 0.613 | VK055_4990 | 0.281 |
| VK055_1115 | 0.161 | VK055_2422 | 0.425 | VK055_3676 | 0.372 | VK055_4991 | -0.283 |
| VK055_1117 | -0.511 | VK055_2423 | 0.011 | VK055_3677 | -0.129 | VK055_4992 | 0.045 |
| VK055_1118 | 0.000 | VK055_2424 | -0.014 | VK055_3678 | -0.551 | VK055_4994 | 0.093 |
| VK055_1119 | -0.337 | VK055_2427 | -0.350 | VK055_3679 | -0.340 | VK055_4995 | -0.342 |
| VK055_1120 | 1.394 | VK055_2428 | 0.588 | VK055_3680 | 1.409 | VK055_4996 | 1.644 |
| VK055_1121 | 0.012 | VK055_2429 | 0.244 | VK055_3681 | 0.740 | VK055_4997 | 0.630 |
| VK055_1122 | 1.468 | VK055_2430 | -0.137 | VK055_3682 | -0.748 | VK055_4998 | -0.815 |
| VK055_1123 | 0.478 | VK055_2432 | 1.133 | VK055_3683 | 0.403 | VK055_4999 | 0.411 |
| VK055_1124 | 0.311 | VK055_2434 | 0.074 | VK055_3684 | -0.053 | VK055_5000 | -0.459 |
| VK055_1126 | -1.476 | VK055_2435 | 0.250 | VK055_3685 | 0.574 | VK055_5001 | 0.432 |
| VK055_1127 | 0.170 | VK055_2437 | 1.042 | VK055_3686 | 1.265 | VK055_5002 | 0.405 |
| VK055_1128 | -0.096 | VK055_2438 | -0.725 | VK055_3687 | 0.525 | VK055_5004 | 0.057 |
| VK055_1130 | -0.729 | VK055_2440 | -0.661 | VK055_3688 | 0.151 | VK055_5005 | 0.279 |
| VK055_1131 | -0.223 | VK055_2441 | -0.013 | VK055_3691 | -1.294 | VK055_5006 | -0.088 |
| VK055_1136 | -0.597 | VK055_2442 | -0.561 | VK055_3692 | 0.842 | VK055_5007 | -0.069 |
| VK055_1137 | -0.144 | VK055_2443 | -0.982 | VK055_3694 | 1.244 | VK055_5008 | -0.888 |
| VK055_1138 | 1.092 | VK055_2444 | 0.686 | VK055_3695 | 0.675 | VK055_5009 | -0.043 |
| VK055_1139 | 0.637 | VK055_2445 | -1.310 | VK055_3696 | 0.137 | VK055_5010 | 0.765 |
| VK055_1140 | -0.353 | VK055_2447 | 0.895 | VK055_3697 | 1.075 | VK055_5011 | -0.561 |
| VK055_1141 | 0.318 | VK055_2448 | 1.433 | VK055_3698 | 0.080 | VK055_5012 | 0.572 |
| VK055_1142 | -0.827 | VK055_2449 | 0.726 | VK055_3699 | -0.092 | VK055_5013 | 1.880 |
| VK055_1143 | 1.288 | VK055_2451 | -0.953 | VK055_3700 | -0.912 | VK055_5014 | -0.170 |
| VK055_1145 | 0.280 | VK055_2452 | 0.782 | VK055_3701 | 0.091 | VK055_5015 | -0.381 |
| VK055_1146 | -0.778 | VK055_2453 | -0.832 | VK055_3703 | 0.135 | VK055_5025 | -1.443 |
| VK055_1149 | 0.502 | VK055_2455 | -0.012 | VK055_3704 | 0.277 | VK055_5026 | 1.381 |
| VK055_1151 | 0.396 | VK055_2456 | 0.751 | VK055_3705 | -0.057 | VK055_5032 | 0.568 |
| VK055_1152 | 0.458 | VK055_2457 | 0.420 | VK055_3706 | 0.774 | VK055_5034 | -0.513 |
| VK055_1153 | 0.278 | VK055_2458 | -1.497 | VK055_3708 | 0.564 | VK055_5035 | -0.300 |
| VK055_1155 | 0.418 | VK055_2460 | 0.611 | VK055_3709 | 0.887 | VK055_5036 | 0.321 |
| VK055_1156 | 1.234 | VK055_2462 | -0.083 | VK055_3710 | 0.288 | VK055_5037 | 0.768 |
| VK055_1157 | -0.722 | VK055_2463 | 0.231 | VK055_3711 | 0.470 | VK055_5038 | -1.205 |
| VK055_1158 | -0.478 | VK055_2464 | -0.062 | VK055_3712 | 0.479 | VK055_5039 | -0.174 |
| VK055_1161 | 0.615 | VK055_2466 | 0.034 | VK055_3713 | -0.029 | VK055_5040 | -0.026 |
| VK055_1162 | -0.661 | VK055_2467 | -0.397 | VK055_3714 | -0.788 | VK055_5041 | 0.036 |
| VK055_1163 | -0.057 | VK055_2468 | 0.826 | VK055_3715 | -0.008 | VK055_5042 | 0.635 |
| VK055_1165 | 0.133 | VK055_2488 | 1.482 | VK055_3716 | -0.395 | VK055_5043 | -1.446 |
| VK055_1168 | 0.131 | VK055_2489 | -0.653 | VK055_3718 | 0.503 | VK055_5044 | 0.423 |
| VK055_1174 | 0.608 | VK055_2490 | 0.272 | VK055_3720 | 1.689 | VK055_5045 | -0.363 |
| VK055_1177 | -0.124 | VK055_2493 | -0.046 | VK055_3721 | 0.164 | VK055_5046 | -0.673 |
| VK055_1179 | 0.998 | VK055_2494 | -0.683 | VK055_3722 | 0.168 | VK055_5047 | -1.465 |
| VK055_1181 | -0.663 | VK055_2495 | 0.139 | VK055_3723 | 0.112 | VK055_5048 | 0.293 |
| VK055_1183 | 0.472 | VK055_2496 | -1.279 | VK055_3724 | -0.637 | VK055_5049 | 0.444 |
| VK055_1184 | -0.795 | VK055_2498 | 0.093 | VK055_3725 | -0.447 | VK055_5050 | 0.934 |
| VK055_1186 | -0.395 | VK055_2499 | -1.257 | VK055_3727 | 1.322 | VK055_5051 | -1.440 |
| VK055_1188 | -1.440 | VK055_2501 | 0.095 | VK055_3728 | 0.991 | VK055_5052 | 0.094 |
| VK055_1190 | -0.509 | VK055_2502 | 0.741 | VK055_3729 | 0.044 | VK055_5053 | -0.525 |
| VK055_1191 | -0.512 | VK055_2503 | 0.149 | VK055_3730 | 0.098 | VK055_5054 | -1.187 |
| VK055_1193 | -1.021 | VK055_2505 | -1.226 | VK055_3731 | -1.108 | VK055_5055 | -0.291 |
| VK055_1194 | 0.290 | VK055_2506 | -0.495 | VK055_3732 | 0.874 | VK055_5056 | -0.082 |
| VK055_1195 | 0.680 | VK055_2507 | -0.687 | VK055_3733 | -0.463 | VK055_5057 | 0.424 |
| VK055_1196 | 0.068 | VK055_2508 | 0.434 | VK055_3734 | 0.338 | VK055_5058 | 0.162 |
| VK055_1198 | 0.103 | VK055_2509 | 0.285 | VK055_3735 | -0.756 | VK055_5059 | 0.447 |
| VK055_1199 | -0.047 | VK055_2512 | 1.330 | VK055_3736 | 0.875 | VK055_5060 | -0.071 |
| VK055_1202 | -0.273 | VK055_2513 | -0.042 | VK055_3737 | -0.142 | VK055_5061 | -0.256 |
| VK055_1204 | -1.464 | VK055_2514 | 0.495 | VK055_3738 | -1.187 | VK055_5062 | 0.315 |
| VK055_1207 | -0.177 | VK055_2515 | 0.208 | VK055_3739 | 0.856 | VK055_5063 | -0.664 |
| VK055_1208 | -1.255 | VK055_2516 | 0.427 | VK055_3740 | 0.100 | VK055_5064 | 0.184 |
| VK055_1209 | 0.411 | VK055_2517 | -0.085 | VK055_3742 | -0.135 | VK055_5065 | -0.780 |
| VK055_1211 | 0.415 | VK055_2518 | 0.639 | VK055_3751 | 0.622 | VK055_5066 | 0.189 |
| VK055_1212 | 0.015 | VK055_2519 | 0.686 | VK055_3752 | -0.111 | VK055_5069 | 3.115 |
| VK055_1213 | -1.248 | VK055_2520 | 0.286 | VK055_3753 | -0.148 | VK055_5070 | 0.170 |
| VK055_1214 | -1.465 | VK055_2521 | 0.577 | VK055_3781 | -0.796 | VK055_5071 | 1.042 |
| VK055_1215 | 0.043 | VK055_2524 | -0.394 | VK055_3782 | 1.567 | VK055_5072 | -0.723 |
| VK055_1216 | 0.507 | VK055_2525 | -0.375 | VK055_3783 | 0.194 | VK055_5074 | 0.346 |
| VK055_1217 | 0.744 | VK055_2526 | -0.698 | VK055_3784 | -1.341 | VK055_5076 | 0.990 |
| VK055_1218 | 1.313 | VK055_2528 | -0.387 | VK055_3787 | -0.699 | VK055_5077 | 0.843 |
| VK055_1219 | 0.391 | VK055_2530 | 0.361 | VK055_3788 | 1.812 | VK055_5079 | 0.527 |
| VK055_1221 | -0.660 | VK055_2531 | -1.205 | VK055_3791 | 0.451 | VK055_5080 | -0.312 |
| VK055_1222 | -0.830 | VK055_2532 | 0.636 | VK055_3792 | -0.275 | VK055_5084 | 0.169 |
| VK055_1223 | 0.478 | VK055_2533 | 0.652 | VK055_3794 | 0.662 | VK055_5085 | -0.406 |
| VK055_1224 | 0.187 | VK055_2534 | -0.574 | VK055_3801 | 0.686 | VK055_5086 | -0.768 |
| VK055_1226 | -1.462 | VK055_2537 | -1.471 | VK055_3802 | 0.334 | VK055_5087 | -0.505 |
| VK055_1229 | 0.612 | VK055_2538 | 1.399 | VK055_3803 | -1.469 | VK055_5089 | 0.958 |
| VK055_1233 | -0.416 | VK055_2539 | 0.124 | VK055_3804 | 0.426 | VK055_5090 | -0.614 |
| VK055_1234 | -1.419 | VK055_2540 | -0.022 | VK055_3808 | 1.008 | VK055_5091 | 0.501 |
| VK055_1235 | 0.909 | VK055_2541 | -0.096 | VK055_3809 | -0.504 | VK055_5093 | 0.541 |
| VK055_1236 | 0.250 | VK055_2542 | -1.278 | VK055_3814 | 0.408 | VK055_5095 | -0.426 |
| VK055_1237 | -0.613 | VK055_2544 | -1.483 | VK055_3815 | -0.679 | VK055_5096 | 0.180 |
| VK055_1238 | 0.288 | VK055_2545 | 0.402 | VK055_3816 | -0.732 | VK055_5097 | 0.641 |
| VK055_1239 | 0.211 | VK055_2546 | 0.097 | VK055_3817 | -0.529 | VK055_5101 | 0.124 |
| VK055_1242 | 0.841 | VK055_2547 | -1.380 | VK055_3821 | 1.176 | VK055_5102 | 0.842 |
| VK055_1243 | -1.008 | VK055_2548 | 0.786 | VK055_3822 | 0.120 | VK055_5103 | -0.763 |
| VK055_1245 | -0.589 | VK055_2549 | 0.126 | VK055_3823 | -0.091 | VK055_5104 | -1.176 |
| VK055_1247 | -0.862 | VK055_2550 | -0.946 | VK055_3824 | 1.320 | VK055_5105 | 1.311 |
| VK055_1248 | -0.089 | VK055_2551 | 1.003 | VK055_3825 | 0.412 | VK055_5106 | -0.629 |
| VK055_1249 | -0.905 | VK055_2557 | 1.217 | VK055_3826 | 1.301 | VK055_5107 | -0.299 |
| VK055_1250 | -0.467 | VK055_2558 | -1.089 | VK055_3827 | -0.545 | VK055_5109 | -0.336 |
| VK055_1251 | -0.255 | VK055_2559 | -1.485 | VK055_3828 | 1.066 | VK055_5110 | -0.410 |
| VK055_1252 | 0.068 | VK055_2561 | 0.307 | VK055_3829 | -0.049 | VK055_5111 | 0.800 |
| VK055_1253 | -0.098 | VK055_2562 | -0.670 | VK055_3830 | 0.247 | VK055_5113 | 1.048 |
| VK055_1254 | -0.157 | VK055_2563 | -0.447 | VK055_3831 | -0.031 | VK055_5114 | -0.096 |
| VK055_1255 | -1.179 | VK055_2564 | -0.146 | VK055_3833 | -0.178 | VK055_5115 | -0.945 |
| VK055_1256 | 0.411 | VK055_2565 | -1.201 | VK055_3834 | 0.107 | VK055_5116 | 0.671 |
| VK055_1258 | 0.034 | VK055_2566 | 0.708 | VK055_3835 | 0.790 | VK055_5117 | -0.900 |
| VK055_1259 | 2.058 | VK055_2567 | 0.309 | VK055_3836 | -0.641 | VK055_5118 | -0.572 |
| VK055_1260 | -0.277 | VK055_2569 | -1.305 | VK055_3837 | 1.653 | VK055_5119 | 0.372 |
| VK055_1261 | 1.634 | VK055_2570 | -0.388 | VK055_3838 | 0.282 | VK055_5120 | 0.624 |
| VK055_1262 | 0.267 | VK055_2571 | 0.763 | VK055_3841 | -0.335 | VK055_5121 | -0.428 |
| VK055_1263 | -1.431 | VK055_2572 | 0.156 | VK055_3843 | -0.133 | VK055_5122 | 1.252 |
| VK055_1264 | -0.819 | VK055_2573 | 0.563 | VK055_3844 | 0.956 | VK055_5123 | -0.232 |
| VK055_1265 | 0.392 | VK055_2574 | -0.719 | VK055_3845 | 0.885 | VK055_5124 | -0.213 |
| VK055_1266 | 0.563 | VK055_2576 | -0.251 | VK055_3846 | -1.182 | VK055_5125 | 0.722 |
| VK055_1267 | -0.056 | VK055_2577 | -0.358 | VK055_3849 | -0.069 | VK055_5126 | -0.046 |
| VK055_1268 | 1.922 | VK055_2578 | 1.526 | VK055_3850 | -0.795 | VK055_5127 | -1.428 |
| VK055_1269 | 0.273 | VK055_2579 | 0.155 | VK055_3851 | -0.583 | VK055_5130 | -0.255 |
| VK055_1270 | 0.315 | VK055_2580 | 0.084 | VK055_3852 | -0.551 | VK055_5132 | 0.585 |
| VK055_1271 | 1.125 | VK055_2581 | -0.023 | VK055_3853 | -0.078 | VK055_5133 | 0.160 |
| VK055_1274 | -0.032 | VK055_2583 | -0.814 | VK055_3854 | 0.779 | VK055_5134 | -0.619 |
| VK055_1275 | -0.273 | VK055_2584 | 0.593 | VK055_3855 | -0.445 | VK055_5135 | 0.175 |
| VK055_1276 | 0.549 | VK055_2585 | -1.472 | VK055_3856 | 0.064 | VK055_5137 | 0.537 |
| VK055_1277 | 0.725 | VK055_2586 | -0.865 | VK055_3857 | 1.177 | VK055_5139 | 0.460 |
| VK055_1278 | 0.707 | VK055_2587 | -0.054 | VK055_3858 | -0.093 | VK055_5140 | 0.220 |
| VK055_1279 | 1.084 | VK055_2588 | 0.756 | VK055_3859 | 0.558 | VK055_5141 | 0.856 |
| VK055_1280 | -0.617 | VK055_2589 | 0.858 | VK055_3860 | -0.409 | VK055_5142 | 0.629 |
| VK055_1281 | 0.715 | VK055_2590 | 0.808 | VK055_3862 | -0.256 | VK055_5146 | -0.740 |
| VK055_1283 | -0.930 | VK055_2591 | 0.273 | VK055_3863 | 0.529 | VK055_5147 | 1.022 |
| VK055_1284 | 0.260 | VK055_2592 | 1.321 | VK055_3864 | -0.579 | VK055_5148 | 0.914 |
| VK055_1285 | -1.285 | VK055_2593 | 0.613 | VK055_3865 | -0.776 | VK055_5149 | 0.710 |
| VK055_1286 | -0.373 | VK055_2594 | 0.641 | VK055_3870 | 0.427 | VK055_5150 | 0.314 |
| VK055_1289 | 0.844 | VK055_2595 | -0.229 | VK055_3871 | -0.642 | VK055_5151 | 1.264 |
| VK055_1290 | 0.720 | VK055_2596 | 0.901 | VK055_3872 | -0.215 | VK055_5152 | -0.298 |
| VK055_1294 | 0.167 | VK055_2597 | 0.352 | VK055_3873 | -0.202 | VK055_5153 | 0.548 |
| VK055_1296 | 0.287 | VK055_2598 | -0.110 | VK055_3874 | -0.351 | VK055_5155 | 0.901 |
| VK055_1297 | 0.912 | VK055_2599 | -0.065 | VK055_3875 | -0.048 | VK055_5157 | -0.798 |
| VK055_1298 | 0.012 | VK055_2600 | -0.706 | VK055_3879 | -0.606 | VK055_5158 | -0.157 |
| VK055_1300 | -0.363 | VK055_2601 | -0.069 | VK055_3885 | -0.433 | VK055_5159 | 0.540 |
| VK055_1301 | 1.061 | VK055_2602 | -1.455 | VK055_3887 | -0.516 | VK055_5160 | 0.346 |
| VK055_1302 | -0.734 | VK055_2604 | 0.017 | VK055_3894 | -0.960 | VK055_5161 | -0.336 |
| VK055_1303 | -0.593 | VK055_2608 | 0.773 | VK055_3895 | -0.964 | VK055_5162 | -0.170 |
| VK055_1305 | -1.476 | VK055_2609 | -0.547 | VK055_3897 | 2.406 | VK055_5164 | 0.620 |
| VK055_1307 | -0.506 | VK055_2610 | 0.954 | VK055_3898 | -0.956 | VK055_5165 | 0.296 |
| VK055_1308 | 0.337 | VK055_2611 | -0.667 | VK055_3904 | 1.476 | VK055_5166 | -1.091 |
| VK055_1309 | 1.129 | VK055_2612 | -1.161 | VK055_3906 | -0.340 | VK055_5167 | -0.875 |
| VK055_1310 | -1.437 | VK055_2614 | 0.410 | VK055_3907 | -0.837 | VK055_5168 | 0.158 |
| VK055_1314 | -0.293 | VK055_2615 | -0.994 | VK055_3908 | 0.332 | VK055_5169 | 0.191 |
| VK055_1316 | 2.179 | VK055_2616 | 0.489 | VK055_3909 | 0.768 | VK055_5171 | 0.780 |
| VK055_1317 | 0.563 | VK055_2617 | 0.475 | VK055_3910 | -1.242 | VK055_5172 | 0.337 |
| VK055_1319 | 0.186 | VK055_2619 | 0.400 | VK055_3911 | 0.406 | VK055_5173 | 0.702 |
| VK055_1324 | 0.870 | VK055_2622 | -0.190 | VK055_3912 | 1.079 | VK055_5174 | 0.116 |
| VK055_1326 | -0.309 | VK055_2624 | 0.499 | VK055_3913 | 0.320 | VK055_5175 | 0.202 |
| VK055_1327 | 0.927 | VK055_2625 | 0.345 | VK055_3914 | 1.008 | VK055_5176 | 1.108 |
| VK055_1328 | -0.291 | VK055_2626 | 0.684 | VK055_3915 | 0.419 | VK055_5177 | -1.225 |
| VK055_1329 | 0.924 | VK055_2627 | 0.329 | VK055_3918 | 0.023 | VK055_5182 | 0.381 |
| VK055_1330 | 0.343 | VK055_2628 | 0.747 | VK055_3919 | 0.410 |  |  |

**Supplemental Figure 1.**
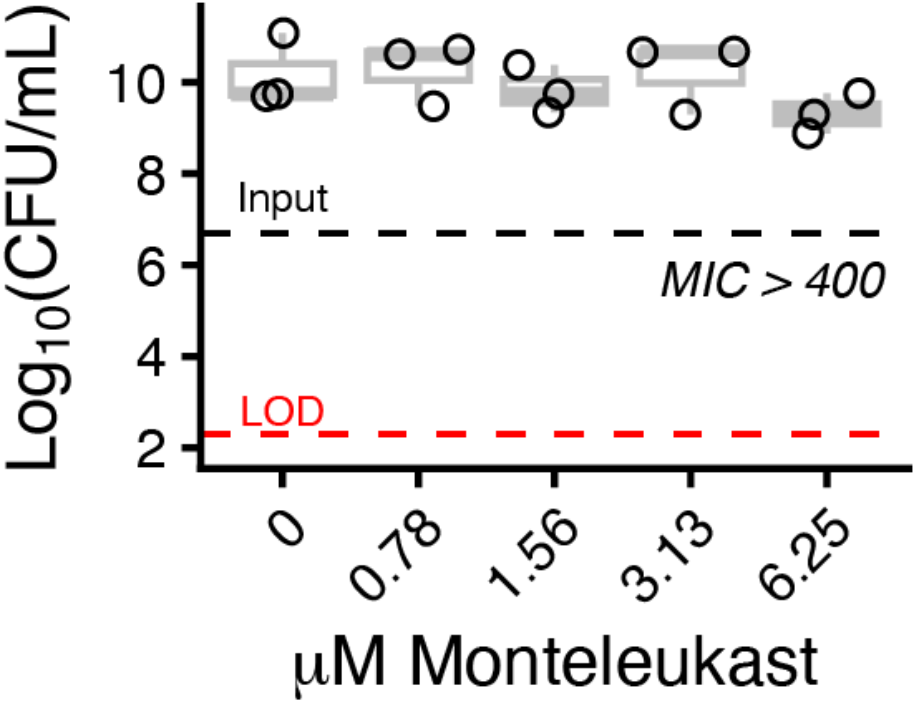
Montelukast MIC/MBC. The minimum inhibitory concentrations (MIC) and minimum bactericidal concentrations (MBC) was determined for montelukast. The black dashed line represents the input CFU/mL concentration, the red dashed line represents the limit of detection (LOD) for the assay, each datapoint represents a single biological replicate, and the boxplots shown the mean and interquartile range, with tails indicating the range.

**Supplemental Figure 2.**
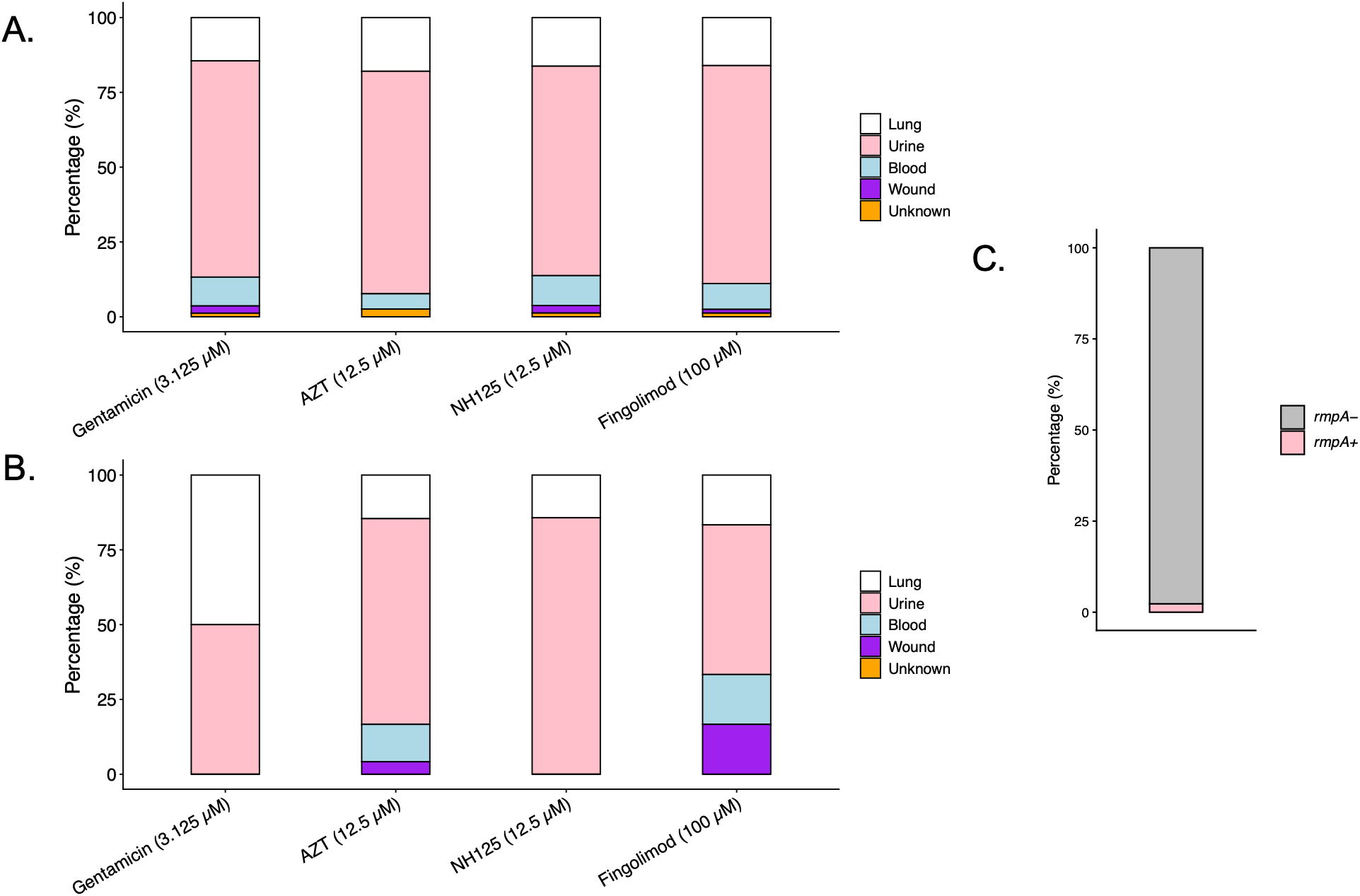
Kp clinical isolate susceptibility to candidate compounds stratified by collection site and presence of *rmpA*. Susceptible (**A**) or resistant (**B**) isolates categorized by their collection site (**Supplemental Table 1**) or presence/absence of *rmpA* (**C**). The wound category included both abscess and pilonidal drainage collection sites. The lung category was composed of Tracheal Aspirate, BALF, Sputum and Bronch Wash.

**Supplemental Figure 3.**
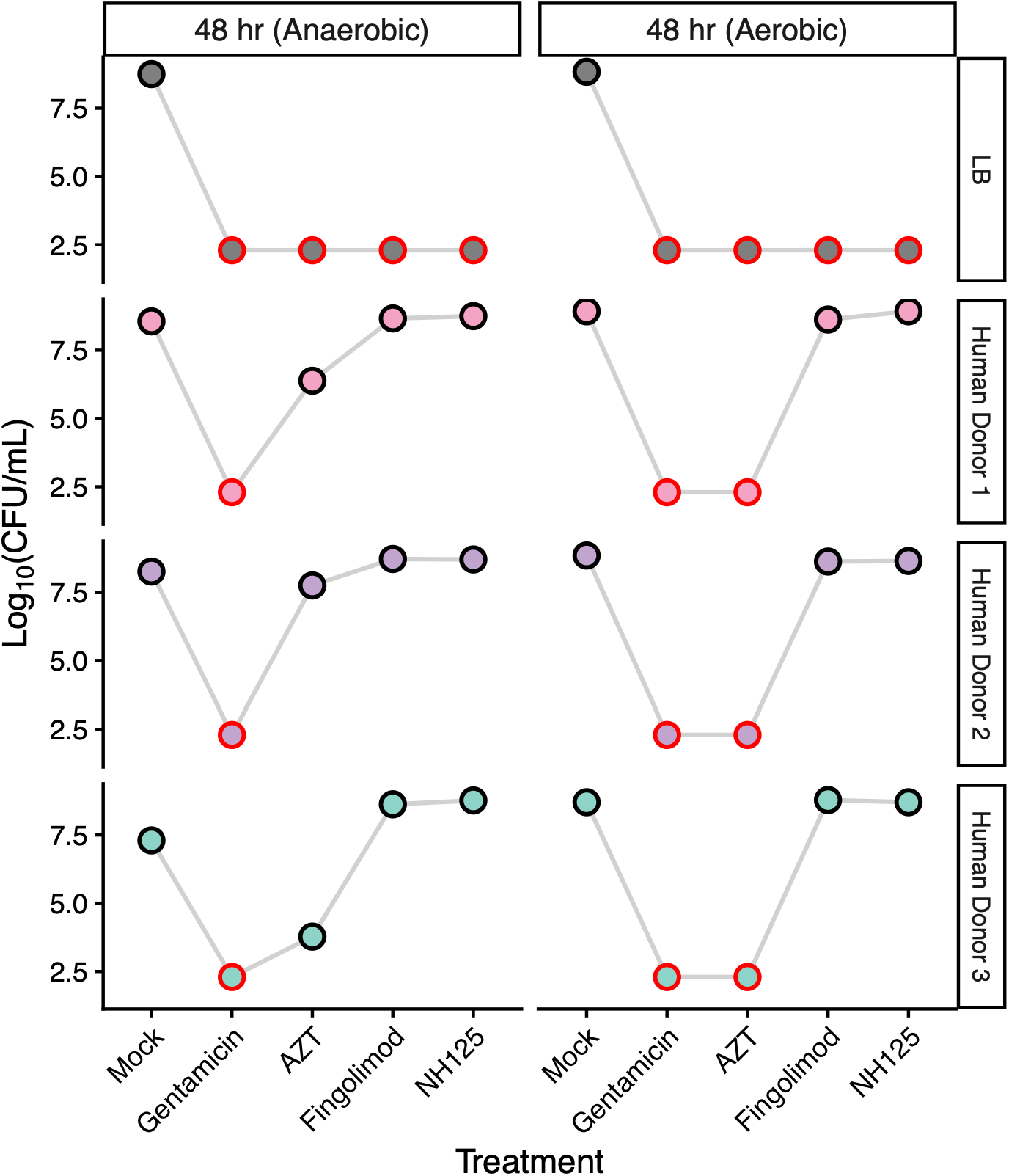
KPPR1 CFUs in human fecal material or in LB under aerobic and anaerobic conditions. KPPR1 was grown overnight at 37° C with shaking (220 rpm) and used the following day for *ex vivo* culture with fecal material from three human donors or in LB medium, in the presence of compounds at 16X their respective MIC values. Cultures were incubated under aerobic or anaerobic conditions and quantified at 48 hours. A red circle indicates the data point is at the limit of detection for this assay.

**Supplemental Dataset 1. L1050 screen data.**

**Supplemental Dataset 2. Raw AZT transposon screen data.**

**Supplemental Dataset 3. Raw study data.**

